# Microtubule detyrosination alters nuclear mechanotransduction and leads to pro-hypertrophic signaling in hypertrophic cardiomyopathy

**DOI:** 10.1101/2025.03.28.646061

**Authors:** Inez Duursma, Linda R. Micali, Edgar E. Nollet, Valentijn Jansen, Janaya A. Malone, Jet S. Bloem, Kenneth Bedi, Kenneth B. Margulies, Stephan A.C. Schoonvelde, Michelle Michels, Nicole N. van der Wel, Jolanda van der Velden, Tyler J. Kirby, Diederik W.D. Kuster

## Abstract

**Background:** Hypertrophic cardiomyopathy (HCM) is characterized by left ventricular hypertrophy and diastolic dysfunction and is accompanied by extensive cytoskeletal remodeling, including increased protein levels of desmin, tubulin and detyrosinated tubulin. We have previously demonstrated that tubulin detyrosination contributes to diastolic dysfunction. Microtubules are connected to the nucleus by the LINC complex, yet the role of cytoskeleton-nucleus interactions in HCM remain poorly understood.

**Objectives:** We investigated whether cytoskeletal remodeling in HCM alters nuclear morphology and mechanics, and modulates hypertrophy-associated signaling pathways.

**Methods:** Nuclear morphology was assessed by electron microscopy and immunofluorescence in cardiac septal tissue from obstructive HCM patients (*N*=19) and non-failing donors (NF, *N*=8), as well as in Wild-Type (WT, *N*=18) and homozygous HCM-associated *Mybpc3* c.2373InsG mice (*Mybpc3*^c.2373InsG^, *N*=19). A novel live-cell imaging approach was used to study nuclear deformation during cardiomyocyte contraction. YAP1 nuclear translocation was measured to evaluate downstream mechanosensitive signaling and detyrosination inhibitor epoY was used to test causality.

**Results:** Nuclei were highly invaginated and enlarged in HCM in both patients and mice. Nuclear deformation during contraction was restricted in HCM cardiomyocytes, indicating altered mechanotransduction. These changes were associated with increased YAP1 nuclear localization and induction of YAP1 target and hypertrophic genes. Inhibition of microtubule detyrosination reduced nuclear invaginations, restored nuclear deformation and decreased YAP1 nuclear translocation.

**Conclusion:** Cytoskeletal remodeling in HCM is associated with altered nuclear morphology and mechanotransduction, accompanied by YAP1 translocation which may contribute to hypertrophic remodeling. Targeting microtubule detyrosination rescues this phenotype, identifying nuclear mechanotransduction as a potential therapeutic target for HCM.

**Unstructured abstract:** Hypertrophic cardiomyopathy (HCM) is characterized by left ventricular hypertrophy and diastolic dysfunction and is accompanied by extensive cytoskeletal remodeling, including increased desmin, tubulin, and detyrosinated tubulin levels, which we previously showed contribute to impaired relaxation. Since microtubules connect to the nucleus via the LINC complex, we investigated whether cytoskeletal remodeling alters nuclear morphology, mechanics, and hypertrophy-associated signaling. Using electron microscopy and immunofluorescence in cardiac septal tissue from obstructive HCM patients and non-failing donors, and immunofluorescence and live-cell imaging in septal tissue from Wild-Type and homozygous HCM-associated *Mybpc3* c.2373InsG mice, we found that nuclei were enlarged and highly invaginated, with restricted nuclear deformation during contraction, indicating altered mechanotransduction. These changes were associated with increased YAP1 nuclear localization and increased expression of YAP1 target and hypertrophic genes. Inhibiting microtubule detyrosination reduced nuclear abnormalities, restored nuclear deformation, and decreased YAP1 nuclear translocation, identifying nuclear mechanotransduction as a potential therapeutic target in HCM.

**Visual abstract:** 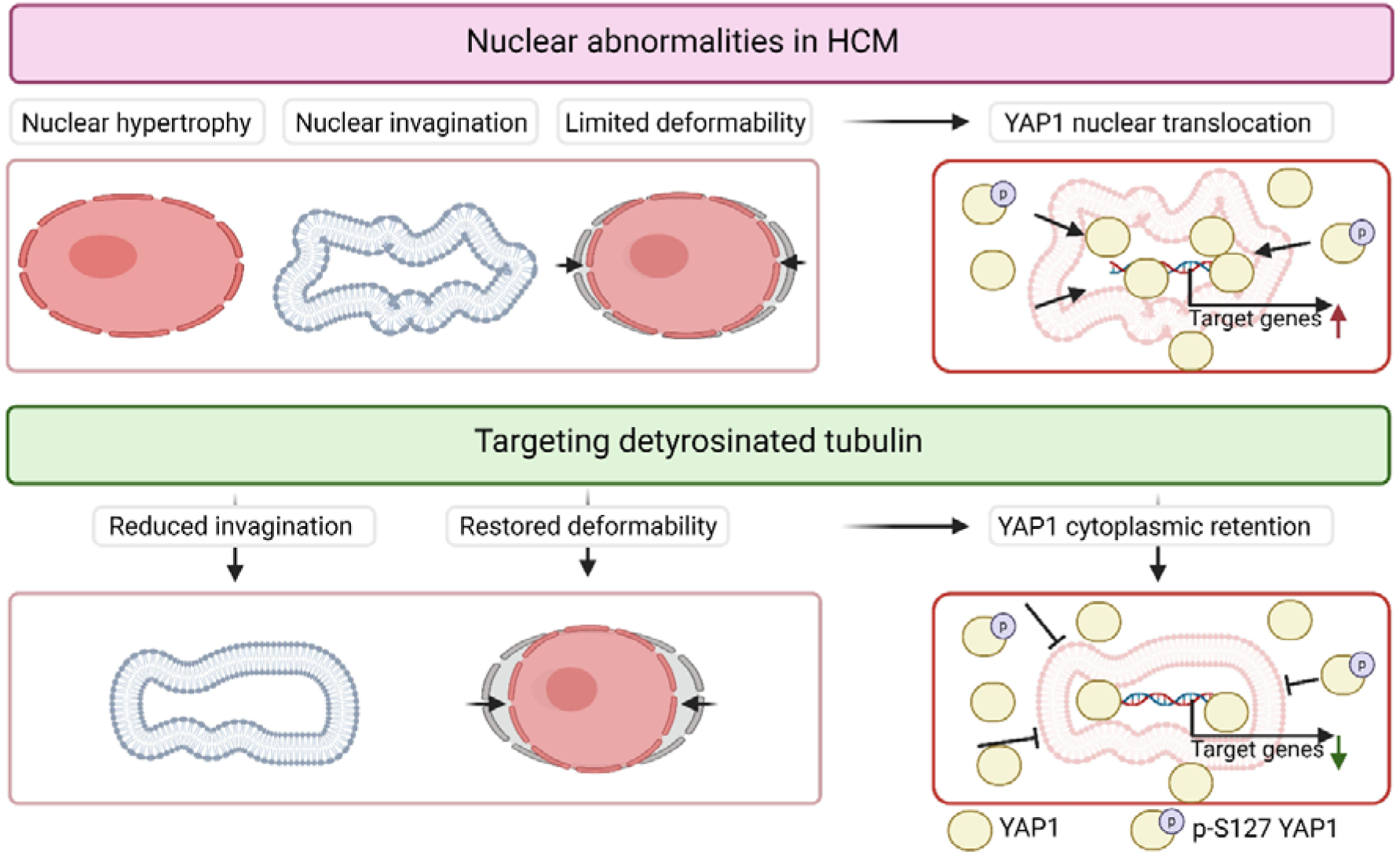

**Highlights:**

- Cytoskeletal remodeling contributes to nuclear abnormalities and impaired mechanotransduction in hypertrophic cardiomyopathy.
- Microtubule detyrosination promotes nuclear invaginations and increased YAP1 signaling in hypertrophic cardiomyopathy.
- Pharmacological reduction of detyrosinated tubulin normalizes nuclear structure and mechanosignaling.
- Targeting nuclear–cytoskeletal coupling may provide new therapeutic strategies for hypertrophic cardiomyopathy.

## Introduction

Hypertrophic cardiomyopathy (HCM) has an estimated prevalence of 1:200-500 and is clinically defined by abnormal thickening of the left ventricular (LV) wall, which cannot be explained by abnormal loading conditions ^1–4^. Besides hypertrophy, HCM is characterized by diastolic dysfunction and an increased risk of arrhythmias. In about half of the patients, HCM is caused by a mutation in one of the genes encoding for sarcomere proteins (Genotype positive [G_positive_]) ^4–6^. In this group, mutations in *MYBPC3*, encoding cardiac myosin-binding protein C (cMyBP-C) are the most common cause. Recently, a growing group of patients in which no known causative pathogenic mutation has been identified (Genotype negative [G_negative_]) ^7,8^. In this group of patients, the underlying etiology is largely unknown.

The pathological mechanisms that lead to hypertrophy, diastolic dysfunction and increased risk of arrhythmias are not completely understood. Recent research performed by our group on HCM patient myocardial tissue has revealed cytoskeletal remodeling with a robust increase in the levels of tubulin, detyrosinated and, acetylated α-tubulin, and desmin ^9–11^. Microtubule remodeling, especially detyrosination of α-tubulin, contributes to slower kinetics by interacting through desmin to the Z-disc and providing a viscoelastic load ^12–14^. Reducing levels of detyrosinated α-tubulin improved relaxation in HCM mouse models in both isolated cells ^9,15^ and in vivo ^16^. In other cardiovascular diseases, the link between microtubule remodeling and cardiac dysfunction is observed ^14,17–24^.

Besides the impact on contractility, microtubules also play an important role in modulating intracellular signaling and gene expression through their interaction with the nucleus. Microtubules are connected to and form a cage around the nucleus by interacting with the Linker of Nucleoskeleton and Cytoskeleton (LINC) complex, which transmits mechanical forces directly to the nucleus ^25–27^. Disturbing the balance of forces on the nucleus imposed by microtubules and desmin alters nuclear structure and genome organization in cardiomyocytes^28^. Indeed, nuclear alterations in the form of nuclear hypertrophy, severe wrinkling of the nuclear membrane and chromatin clumping have already been observed in HCM years ago ^29,30^, although these abnormalities have not been linked to microtubule remodeling. Little is known about the interaction of the microtubules with the nucleus during contraction and the implications this may have for the HCM pathophysiology.

The main goal of this study was to determine whether microtubule changes lead to alterations in nuclear morphology, deformability and mechanotransduction and to identify therapeutic potential in HCM. Therefore, we first studied baseline morphology and function in HCM myocardium and applied pharmacological tools to disrupt microtubule detyrosination and assessed functional consequences. We show that in HCM nuclei were considerably larger, highly invaginated, less deformable and more permeable to transcription factors that shuttle between cytosol and the nucleus. These nuclear abnormalities could be rescued by using microtubule-modifying drugs that specifically decrease microtubule detyrosination levels. Our study shows that microtubules, through mechanotransduction, play a central role in HCM and that intervening in the process might be a promising therapeutic strategy.

## Methods

### Human myocardial tissue samples

Cardiac interventricular septum tissue was acquired from 19 HCM patients during a septal myectomy surgery, of which 10 were G_positive_ and 9 G_negative_. This was approved by the medical ethics review committee of the Erasmus Medical Center and written informed consent was obtained from each patient before surgery. An overview of all known clinical parameters is provided in Table S1-S5. As controls, the left ventricular free wall tissue of 8 non-failing donors without any history of cardiac disease was obtained during organ donation from brain-dead donors. This was in accordance with protocols and ethical regulations approved by Institutional Review Boards at the University of Pennsylvania and the Gift-of-Life Donor Program (PA, USA). All tissue samples were kept cold on ice until flash freezing in liquid nitrogen for protein analysis, fixation in 2.5% (v/v) glutaraldehyde, 2% (v/v) paraformaldehyde in 0.1 mol/L phosphate buffer (EM fixation buffer) for transmission electron microscopy (TEM) or fixation in 4% paraformaldehyde for histological analysis.

### Transmission electron microscopy

Small pieces (2-4 mm^3^) of fresh left ventricular free wall tissue of non-failing donors (N=4) and interventricular septum tissue of HCM patients (N=19) (Table S2) were collected during myectomy surgery, fixed in EM fixation buffer overnight and preserved in 1% (v/v) paraformaldehyde at 4°C until further use. Tissue processing was performed as previously described ^31^. Images of 5-17 cardiomyocyte nuclei per sample were taken with a magnification of 6800-11000x using a FEI Tecnai T12 Transmission Electron Microscope G2 Spirit Biotwin using Veleta camera Plus integrated software and analysed using Fiji ^32^. To determine the amount of invaginations in each nucleus, nuclei were segmented manually and invaginations were counted by an observer blinded to the sample ID.

### Western blot

Homogenates of human cardiac interventricular septum tissue of HCM patients (N=19) and left ventricular tissue of non-failing donors (N=8) (Table S5) were made as previously described ^9^. A total of 4.5 µg of protein was loaded onto 4-15% precast Criterion™ gradient gels (Bio-Rad Laboratories Inc.). Samples from HCM patients and non-failing donors were loaded in a random order to avoid bias. Electrophoresis was performed at 100V in sodium dodecyl sulfate (SDS) electrophoresis buffer until the dye reached the bottom of the gel. Wet tank transfer to a PVDF membrane was performed at 0.3 A for 120 min, after which the membranes were blocked in 5% (w/v) milk or 3% (w/v) bovine serum albumin (BSA) in tris-buffered saline with 0.1% (v/v) tween (TBS-T). Primary antibodies were incubated in 3% (w/v) BSA in TBS-T overnight at 4°C. Antibody concentrations can be found in Table S6. Membranes were washed in TBS-T and incubated with the secondary antibody in 3% (w/v) BSA in TBS-T for 1h at room temperature (RT). Membranes were washed in TBS-T, incubated with enhanced chemiluminescent detection reagent (Amersham) and imaged using the Amersham Imager 600 (GE Healthcare Bio-Sciences AB). Protein levels were quantified using ImageQuant (Cytiva, USA) and normalized to GAPDH protein levels. Normalization over multiple blots was done using the normalizing control sample.

### Quantitative Real-Time PCR

Total RNA was isolated from heart tissue of non-failing donors (N=4) and HCM patients (N=10) (Table S4) using the miRNeasy Tissue/Cells Advanced Micro Kit (Qiagen) according to manufacturer’s protocol. 1 µg of total RNA was reverse transcribed using the iScript cDNA synthesis kit (BioRad) and quantitative real-time PCR was performed with iQ SYBR Green Supermix (BioRad) with the CFX384 Real Time PCR Detection System (BioRad). Gene expression levels were normalized to PSMB2 and OARD1, previously described as stable genes in non-failing donor and HCM heart tissue ^33^, and evaluated using the 2^-ΔCT^ method. The primer sequences are listed in Table S7.

### Mybpc3_c.2373InsG_ mouse model

An established heterozygous (*Mybpc3^+/InsG^*) and homozygous (*Mybpc3^InsG/InsG^) MYBPC3_c.2373InsG_* mouse model was used as previously described ^9,15^. The *Mybpc3^+/InsG^* mice do not develop an HCM-like phenotype, while the *Mybpc3^InsG/InsG^* mice do develop an HCM-like phenotype, including cardiac hypertrophy and left ventricular systolic and diastolic dysfunction ^15^. Additionally, *Mybpc3^InsG/InsG^*mice present with increased levels of detyrosinated microtubules and desmin ^15^. There are no sex differences ^15^. 24 wild-type (WT), 6 Mybpc3^+/InsG^ and 25 Mybpc3^InsG/InsG^ mice of both sexes, aged 4-6 months, were used for all studies. Mouse characteristics and proof of model consistency of mice used for the embedding of whole mouse hearts can be found in Table S8 and S9. Mouse characteristics of mice used for cardiomyocyte isolation and the number of mice per study can be found in Table S10.

Animal experiments were conducted following the Guide for the Animal Care and Use Committee of the VU University Medical Center and with approval of the Guide for the Animal Care and Use Committee of the VU University Medical Center.

### Immunofluorescence labeling

Six whole mouse hearts per group from 4-6-month-old Wild-Type (WT), *Mybpc3^+/InsG^* mice and *Mybpc3^InsG/InsG^*mice and human cardiac interventricular septum tissue of HCM patients (N=11) and non-failing donors (N=6) (Table S3) were collected and fixed in 4% paraformaldehyde for at least 24h to ensure complete infiltration of the fixative into the tissue. Next, the tissues were dehydrated using an automated tissue processor and embedded in paraffin. Paraffin-embedded mouse hearts were sectioned using a microtome (SLEE) at a thickness of 6 µm for 2-dimensional (2D) applications and a thickness of 20 µm for 3-dimensional (3D) analysis. Slides were deparaffinized in xylene and rehydrated using a hydration series of 100%, 95%, 70% (v/v) ethanol followed by dH_2_O. Antigen retrieval was performed in citrate buffer at 96°C, after which slides were cooled in phosphate-buffered saline with 1% (v/v) tween (PBS-T).

Immunofluorescent staining was performed on preprocessed mouse heart sections and plated isolated cardiomyocytes fixed in 4% paraformaldehyde. Slides were permeabilized with 0.25% (v/v) Triton X-100 in PBS for 15 min and blocked in 1% (w/v) BSA in PBS (blocking buffer) for 30 min. Primary antibodies were incubated in blocking buffer overnight at 4°C. An overview of all used antibody concentrations is provided in Table S11. After washing in PBS, slides were incubated with the secondary antibodies in blocking buffer for 1h at RT. Slides were then incubated with DAPI (1:1000) (Sigma-Aldrich) and/or WGA (1:400) (Invitrogen) for 15 min and washed in PBS. Heart sections were mounted with mowiol and isolated cardiomyocytes were preserved in PBS. Fluorescent images were collected using the Nikon A1R confocal microscope (60x magnification) and analyzed.

### Cardiomyocyte isolation and live cell imaging of nuclear deformation

Cardiomyocytes were isolated from hearts of 4-6-month-old mice (8 WT, 9 *Mybpc3^InsG/InsG^*) for live-cell imaging or staining following a previously described protocol ^9,34^. After isolation, cardiomyocytes were plated on laminin-coated dishes (10 μg/ml, Sigma-Aldrich) in Medium 199 (Lonza), supplemented with 1% penicillin/streptomycin and 5% bovine serum for 1h at 37°C and 5% (v/v) CO_2_, after which the cells are fixed or used for live cell imaging.

For live cell imaging medium was changed to culture medium consisting of medium 199 (Lonza), 1% penicillin/streptomycin, 1× insulin-transferrinLJsodium selenite supplement (Sigma-Aldrich), and 0.5 µmol/l cytochalasin D (Life Technologies). The culture medium was supplemented with live cell dye SPY505-DNA (1:1000, Spirochrome) to stain the nucleus. The cells were incubated at 37°C and 5% (v/v) CO_2_ for at least 1h. Fifteen min before imaging, culture medium was replaced by modified Tyrode’s solution (10 mmol/l HEPES, 133.5 mmol/l NaCl, 5 mmol/l KCl, 1.2 mmol/l NaH_2_PO_4_, 1.2 mmol/l MgSO_4_, 11.1 mmol/l glucose, 5 mmol/l sodium pyruvate; pH 7.4 at 37°C) supplemented with SPY505-DNA (1:1000). The dishes were paced at 1 Hz, 8 V at intervals of 20-30s and imaged (20 cardiomyocytes per mouse, time-lapse, single stack of middle of nucleus) using the Nikon A1R with climate control (37°C and 5% (v/v) CO_2_) (Figure 3A).

### Pharmacological manipulation of microtubules for nuclear deformability experiments

To study nuclear deformation of isolated cardiomyocytes (5 WT, 3 *Mybpc3^InsG/InsG^*) with microtubule modifying compounds, cells were treated with vehicle dimethyl sulfoxide (DMSO, 0.1% v/v) for 1h, epoY (20 µmol/l, Sigma-Aldrich) for 2h or Nocodazole (10 µmol/l, Cayman Chemicals) 1h in culture medium at 37°C and 5% (v/v) CO_2_. After incubation, cells were live-imaged as described previously.

To investigate the effect of epoY and nocodazole on nuclear invagination and Yes-axxociated protein 1 (YAP1) localization, the above-mentioned culture conditions were repeated, followed by 5 minutes of pacing at 1Hz, fixing and staining for lamin A/C and YAP1 as described above (4 WT, 4 *Mybpc3^InsG/InsG^*).

### Image analysis

Before analysis, images were deconvolved and post-processed (e.g. contrasting, filtering) using NIS-elements (Nikon) to improve the analysis workflows.

#### Nuclear morphology

To determine nuclear parameters in 2D and 3D from mouse and 2D human FFPE heart tissue, an automated analysis pipeline was generated in NIS-elements (Nikon), which segments the nucleus based on the DAPI signal and cell based on the WGA signal, thereby generating nuclear and cellular morphology parameters, such as nuclear and cellular volume, width and length.

#### Nuclear invagination and lamin A/C deposition

Nuclear invagination length was determined using Fiji ^32^ and the Ridge detection Fiji plugin ^35^. This plugin generates segments based on the lamin A/C signal from 2D maximum intensity projection images of isolated cardiomyocytes (Figure 2E). All segments within the cell were considered an invagination. All separate lengths of the invaginations were summed to generate the total invagination length. Furthermore, lamin A/C deposition was determined using NIS elements. This pipeline segments the nucleus based on the DAPI signal and generates the Lamin A/C intensity at the outside of the nucleus, thereby providing lamin A/C deposition.

#### Chromatin condensation

Chromatin condensation was determined using an automated pipeline generated with NIS-elements. Images were generated from the nuclear deformation videos, taking a frame when the cell is not contracting. The nucleus is segmented based on thresholding of the SPY505-DNA dye signal and filled to determine total nuclear area. Consequently, the densely packed chromatin is segmented using the SPY505-DNA dye signal with the detect regional maxima and signal threshold node to generate the dense chromatin area of the nucleus. SPY505-DNA dye stains mostly densely packed chromatin. The chromatin condensation is calculated by dividing the dense chromatin area by the total nuclear area.

#### YAP1 localization

To assess YAP1 localization, a pipeline was made in Fiji for human or NIS-elements for mouse images of FFPE heart tissue stained for YAP1, WGA, DAPI and lamin A/C. A Z-stack through the tissue’s thickness was acquired from which a single Z-stack was selected at the point where the nucleus was biggest for the cell being analyzed. The nucleus was manually segmented based on the DAPI signal (human) or lamin A/C signal (mouse), and the cell was manually segmented based on the WGA signal. The cytoplasmic compartment was created by subtracting the nuclear area from the cell area. This resulted in both morphological parameters and YAP1 intensity and integrated density in the nucleus and the cytoplasm.

#### Nuclear deformation

Nuclear deformation was analyzed with NIS-elements. The analysis pipeline automatically segmented the nucleus generating nuclear morphological parameters for every frame of the time-lapse. Contraction was analyzed using the brightfield channel with the SarcOptiM plugin for Fiji ^36^ by drawing a line over the sarcomeres near the nucleus, which generated a sarcomere length per frame. The nuclear deformation parameters match the sarcomere shortening over time.

### Statistical analysis

Data is presented as mean±standard error of the mean (SEM) in superplots ^37^. Statistics was performed on the average per human/mouse data points using GraphPad Prism 9.1.0 and 10.2.0 (GraphPad Software, USA). Data were tested for normality and accordingly analyzed using a Mann–Whitney test or an unpaired Student’s t-test when comparing 2 groups and a Kruskal–Wallis test with Dunn’s multiple comparisons test or a one-way analysis of variance (ANOVA) with Tukey’s multiple comparisons test when comparing >2 groups. The effect of treatments was tested using a paired two-way ANOVA. Correlations were shown by Pearson’s R (two-tailed, 95% confidence interval) or linear regression. Data was considered significant with a *p* < 0.05.

## Results

### HCM patients have nuclear abnormalities

To assess nuclear morphology in HCM myocardium, TEM was performed on septal myectomy tissue from HCM patients and non-failing donors. In a previous study, we found abnormal mitochondrial morphology and organization in HCM samples in TEM images ^31^. Here, we report on abnormal nuclear morphology in HCM patients (Figure 1A). HCM patients displayed more invaginations per nucleus than non-failing donors, independent of mutation status or sex (Figure 1B, S2A). Immunofluorescent images showed an increased nuclear area, increased nuclear perimeter and decreased nuclear circularity that were not influenced by genotype or sex (Figure 1C, 1D, S1, S2). Cell area in longitudinal-sectioned tissue tended to increase but was not significantly larger (Figure 1E), unlike cellular hypertrophy observed in transversal sections ^38,39^. The number of invaginations in the nuclei per HCM patient was correlated to their clinical parameters (Figure 1F, Table S12), which reveals a relationship (*R*=0.59, *p*=0.008) between the degree of hypertrophy, i.e. interventricular septum thickness (IVS), and the number of nuclear invaginations (Figure 1G). Together, this data revealed that HCM nuclei have abnormalities, which are present in all patient populations and might, therefore, contribute to or be a consequence of the development of HCM.

**Figure 1:**
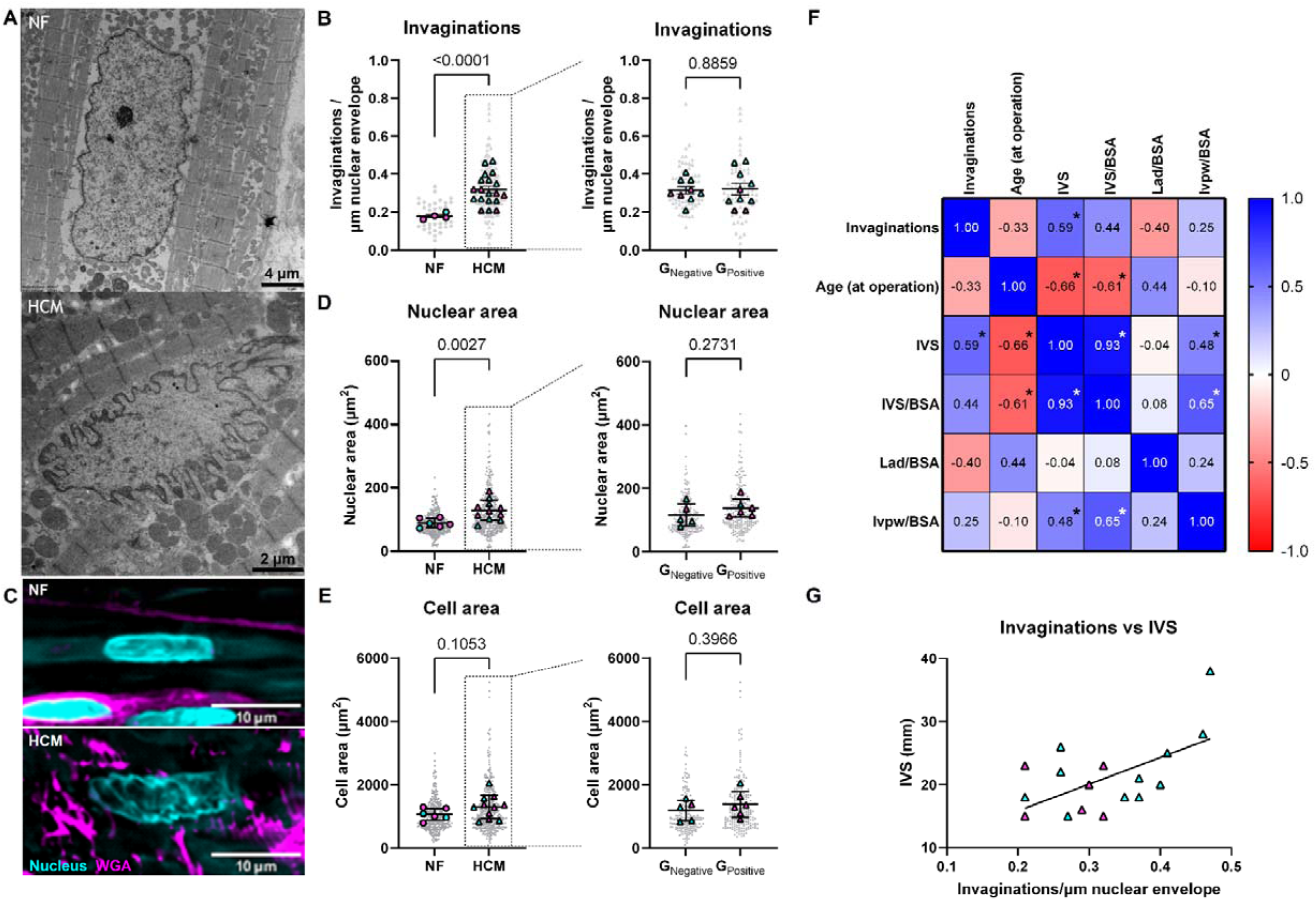
Nuclear abnormalities in human patients with hypertrophic cardiomyopathy. Representative EM images of cardiomyocyte nuclei of non-failing donors (*N*=4) and HCM patients (*N*=19) (A). Quantification of the number of invagination per nucleus normalized for the length of the nuclear envelope to correct for size in NF donors and HCM patients (left panel) and G_Negative_ and G_Positive_ (right panel) (B). Representative fluorescent images of FFPE non-failing donor (*N*=6) and HCM patient (*N*=11) tissue stained for the nucleus (cyan) and WGA (magenta) (C). Quantification of the nuclear area of NF donors and HCM patients (left panel) and G_Negative_ and G_Positive_ (right panel) (D) Quantification of cell area of NF donors and HCM patients (left panel) and G_Negative_ and G_Positive_ (right panel) (E). Pearson R correlations of nuclear invaginations with several clinical parameters (F, *=p<0.05). Simple linear regression of the number of invaginations versus IVS (G, *p*=0.0083, *r*^2^=0.3437) Data are expressed as mean ± standard error of the mean. Every grey symbol represents the value of a single nucleus, *n*, and every colored symbol represents the average value per non-failing donor or HCM patient, *N*. Female are shown in magenta, male are shown in cyan, NF are shown as circles and HCM are shown as triangles. (B) NF: *N*=4 *n*=44; HCM: *N*=19 *n*=147; G_Negative_: *N*=9 *n*=75; G_Positive_: *N*=10 *n*=72, (D) NF: *N*=6 *n*=281; HCM: *N*=11 *n*=410; G_Negative_: *N*=5 *n*=190; G_Positive_: *N*=6 *n*=220, (E) NF: *N*=6 *n*=281; HCM: *N*=11 *n*=420; G_Negative_: *N*=5 *n*=190; G_Positive_: *N*=6 *n*=230. Statistical tests: Welch’s t-test and unpaired t-test (B, D, E); simple linear regression (G). IVS, interventricular septum thickness; BSA, body surface area; Lad, left atrial diameter; lvpw, left ventricular posterior wall thickness.

### *Mybpc3^insG/insG^* mice also possess nuclear abnormalities

To further investigate the mechanisms underlying the nuclear abnormalities, and the potential implications for HCM pathogenesis, we utilized the *Mybpc3^insG/insG^* mouse model we generated ^15^. First, we assessed whether nuclear morphology in the mouse model recapitulates our observations in human HCM tissue. Cardiomyocytes from *Mybpc3^insG/insG^* mice showed both cellular hypertrophy and nuclear hypertrophy, with a relatively larger increase in nuclear area (1.80 fold) compared to cell area (1.47 fold) (Figure 2A-D). This increase in nuclear area resulted from an increase in both nuclear length and width (Figure S3). Similar results were found in 3-dimensional nuclear analysis (Figure S4A). Nuclear volume was increased (Figure S4B), mainly caused by an increase in nuclear width (Figure S4D), resulting in a decrease in nuclear elongation (Figure S4C). Nuclear parameters correlate with the clinical parameters of the mice obtained with echocardiography (Figure S5, Table S13, Figure S6, Table S14). Similar to human HCM patients, *Mybpc3^insG/insG^*mice displayed an increase in the nuclear envelope invaginations (Figure 2E). Nuclear invagination length was positively correlated with nuclear area (Figure S7A,B; Tables S15, S16). To account for this relationship, we stratified nuclei into three bins based on nuclear area and compared invagination length between genotypes within each bin. In all three area ranges, nuclear invagination length was consistently increased in *Mybpc3^insG/insG^*mice compared to WT (Figure S7C). These results indicate that the *Mybpc3^insG/insG^*mouse model is relevant to study nuclear abnormalities in HCM.

**Figure 2:**
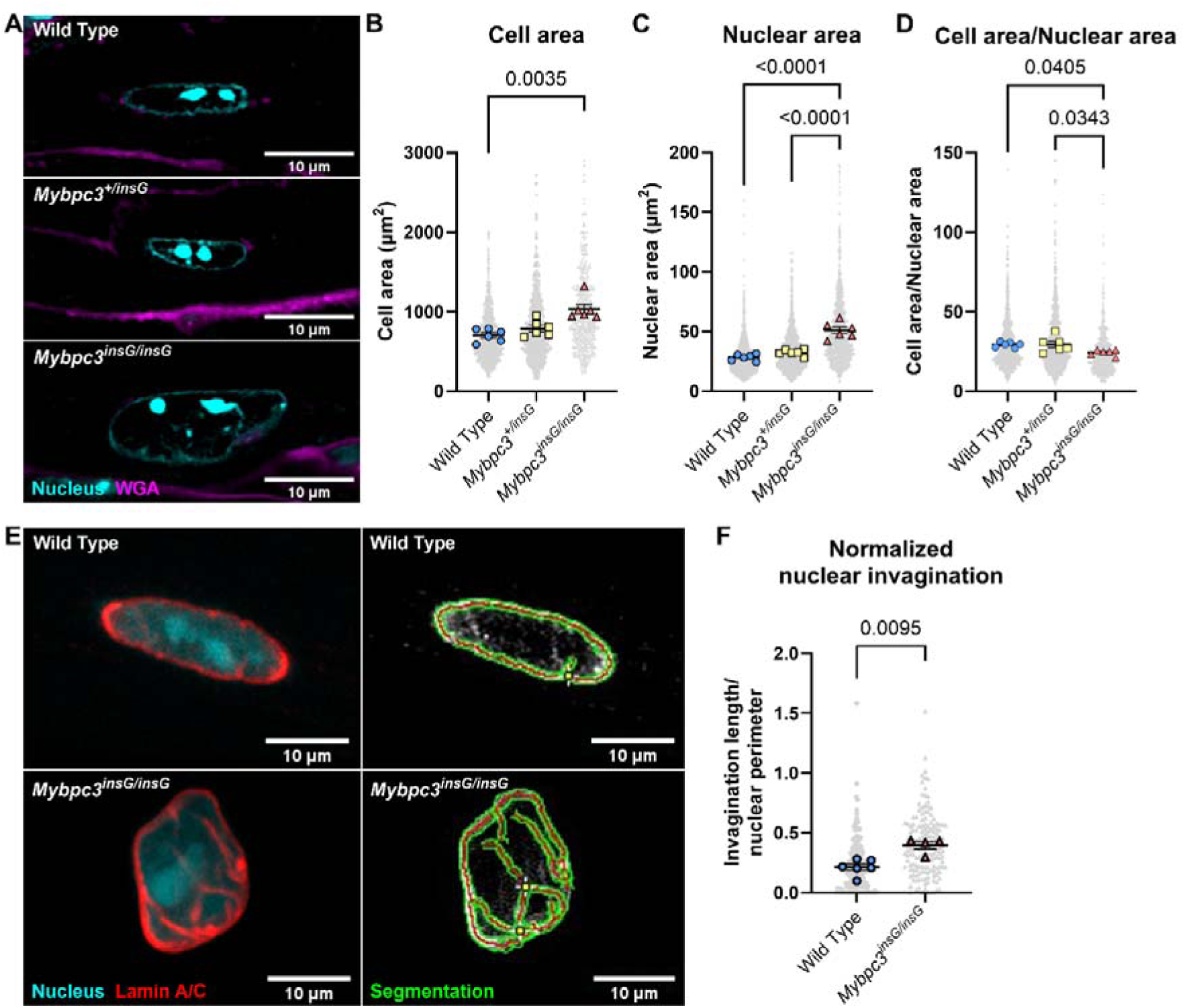
2D nuclear morphology of WT, *Mybpc3^+/insG^* and *Mybpc3^insG/insG^*mice and nuclear invaginations in nuclei of WT *and Mybpc3^insG/insG^*mice. Representative images of embedded mouse heart tissue of Wild Type (*N*=6), *Mybpc3^+/insG^* (*N*=6) and *Mybpc3^insG/insG^* (*N*=6) mice stained with DAPI (cyan) and WGA (magenta) (A). Quantification of the cell area and nuclear area in Wild Type, *Mybpc3^+/insG^* and *Mybpc3^insG/insG^* tissue. Nuclear area of both mononucleated and multinucleated cells were included, since all groups contain ∼90% mononucleated cells and ∼10% binucleated cells (B, C). The cell to nuclear area ratio was calculated by dividing the cell area by the nuclear area. Only nuclei for which also cell size was known could be included. (D). Representative images (left) and segmentation (right) of Lamin A/C (red) in maximum intensity z-projection images of isolated cardiomyocytes of WT (*N*=6) and *Mybpc3^insG/insG^* (*N*=4) (E). Quantification of the normalized nuclear invagination calculated as the sum of every separate length of lamin A/C protruding into the nucleus divided by the Lamin A/C perimeter length (F). Data are expressed as mean ± standard error of the mean. Every grey symbol represents the value of a single cell (B), nucleus (C, F) or ratio of cell to nucleus (D), *n*, and every colored symbol represents the average value per single mouse, *N*. (B) Wild Type: *N*=6 *n*=828; *Mybpc3^+/insG^*: *N*=6 *n*=803; *Mybpc3^insG/insG^*: *N*=6 *n*=524, (C) Wild Type: *N*=6 *n*=1371; *Mybpc3^+/insG^*: *N*=6 *n*=1371; *Mybpc3^insG/insG^*: *N*=6 *n*=1064, (D) Wild Type: *N*=6 *n*=900; *Mybpc3^+/insG^*: *N*=6 *n*=896; *Mybpc3^insG/insG^*: *N*=6 *n*=581, (F) Wild Type: *N*=6 *n*=205; *Mybpc3^insG/insG^*: *N*=4 *n*=155. Statistical tests: Kruskal-Wallis test (B), one-way ANOVA (C, D) and Mann-Whitney test (F).

One explanation for the increase in nuclear hypertrophy would be an increase in the ploidy of nuclei during cardiac hypertrophy ^40^. The fluorescence intensity of DNA intercalating dyes (i.e. DAPI) can be used to assess ploidy ^41^. While the integrated density of the DAPI nuclear fluorescent signal increased due to increased nuclear volume (Figure S4F), the mean nuclear fluorescent intensity remained unchanged (Figure S4E). A linear correlation was observed between integrated density and nuclear volume (Figure S4G). It was shown before that an increase in DAPI fluorescent intensity relates to an increase in DNA content ^40–42^. This suggests that nuclear hypertrophy is associated with an increase in ploidy.

### Nuclear deformation is restricted in *Mybpc3^insG/insG^* mice

The heart continuously undergoes contraction-relaxation cycles. The sarcomere is physically coupled to microtubules at the Z-disk, when α-tubulin is detyrosinated ^12,43^ and transmits mechanical and biochemical signals to the nucleus. Increased levels of detyrosination of microtubules facilitate the coupling of the microtubules to the sarcomere, leading to microtubule buckling and cardiomyocyte stiffening ^12^. Our *Mybpc3^insG/insG^* mouse hearts possess higher levels of detyrosinated microtubules ^15^, probably resulting in more microtubules coupled to the Z-disk, which can induce altered nuclear mechanotransduction ^44^. Therefore, nuclear deformability was assessed during cardiomyocyte contraction and relaxation (Figure 3A, 3B, Video S1). We observed similar sarcomere shortening and contraction kinetics in WT and *Mybpc3^insG/insG^* cardiomyocytes (Figure 3C, 3I, S8), while *Mybpc3^insG/insG^* cardiomyocytes displayed prolonged relaxation typical for HCM (Figure S8). During contraction, nuclear area and length decreased to a lesser extent in *Mybpc3^insG/insG^* cardiomyocytes (Figure 3D, 3E, 3J, 3K), with no change in nuclear width during contraction (Figure 3F, 3L). Importantly, there was no relationship between the nuclear deformability and starting morphology (Figure S9, Table S17, S18). Sarcomere-nuclear strain maps were generated by plotting the change in sarcomere length versus the change in nuclear length during systolic compression and diastolic re-lengthening (Figure 3G). The strain curve highlighted a mismatch in sarcomere-nuclear coupling in *Mybpc3^insG/insG^* mice (Figure 3G). After 1/3^rd^ of maximal sarcomere shortening, the nucleus did not deform any further, while the sarcomere continued to shorten. This indicates there is no linear relationship between sarcomere and nuclear shortening as observed in the WT. Furthermore, there seems to be a quicker re-lengthening of the nucleus during relaxation in *Mybpc3^insG/insG^*, as can be seen by the smaller change in the shortening of the nucleus during relaxation at similar changes of sarcomere shortening during contraction and relaxation (Figure 3G). This is quantified by the hysteresis of the nuclear length deformation (Figure 3H). This suggests that cardiomyocytes from HCM tissue show impaired nuclear mechanotransduction during contraction, possibly due to nuclear stiffening or abnormal nuclear-cytoskeletal coupling.

**Figure 3:**
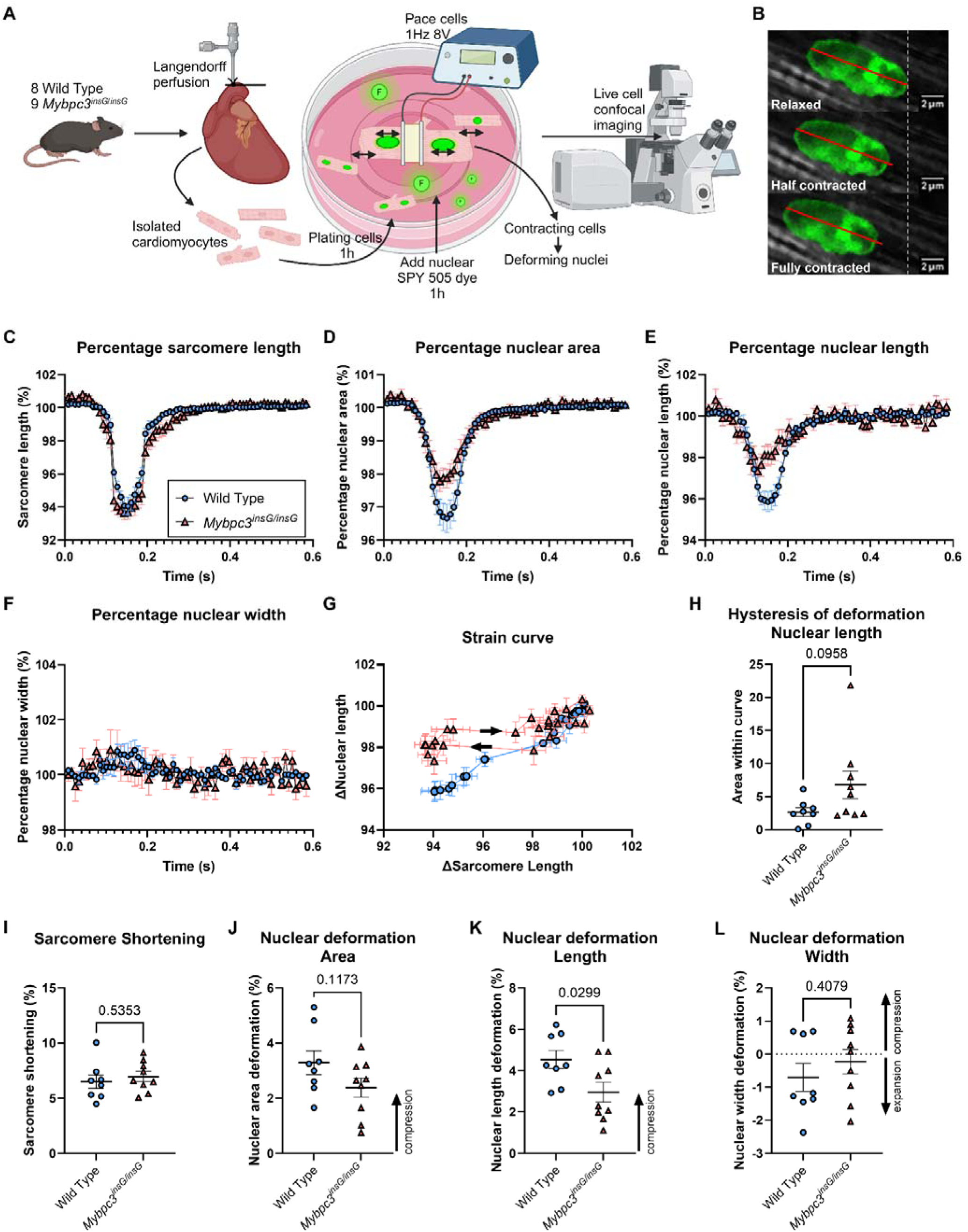
Nuclear deformation during contraction in WT and *Mybpc3^insG/insG^* mice. A schematic of the experimental set-up (A) and immunofluorescent images during a contraction showing nuclear deformation (B). The red line depicts the length of the nucleus at full relaxation of the cardiomyocyte. The white dotted line shows the starting position of the nucleus at full relaxation of the cardiomyocyte indicating movement of the nucleus during contraction (B). Representation of the sarcomere length (C), nuclear area (D), nuclear length (E) and nuclear width (F) over time in WT and *Mybpc3^insG/insG^* cardiomyocytes. The sarcomere-nuclear strain curve was generated by calculating the change in sarcomere length versus the change in nuclear length (G). Quantification of the hysteresis in the deformation of the nuclear length during a contraction. The hysteresis was determined by calculating the area within the strain curve (H). Quantification of the maximum sarcomere shortening (I), nuclear area deformation (J), nuclear length deformation (K) and nuclear width deformation (L). Every colored symbol represents the value of the average of the mice in their group and the error bars depict the standard error of the mean (C-G). For the quantifications every colored symbol represents the average value per single mouse, *N,* and data are expressed as mean ± standard error of the mean (H-L). Wild Type: *N*=8 *n*=223; *Mybpc3^insG/insG^*: *N*=9 *n*=145, Statistical tests: Unpaired t-test (H-L).

### YAP1 localizes to the nucleus irrespective of Hippo signaling thereby inducing abnormal signaling in HCM patients

A stiffer, malformed nucleus and/or impaired mechanotransduction to the nucleus during contraction could lead to changes in signaling events that depend on cytoplasmic/nuclear translocation. YAP1 is a mechanosensitive transcriptional regulator that is known to play a role in the induction of cardiac hypertrophy ^45,46^. Regulation of YAP1 is responsive to stiffness and cytoskeletal tension and dependent on the phosphorylation of YAP1. An increase in stiffness and cytoskeletal tension and a decrease in YAP1 phosphorylation leads to nuclear translocation of YAP1 and downstream target gene transcription ^44,46^. In addition, YAP1 nuclear translocation can occur as the direct result of increased nuclear tension ^47^. We hypothesized that the observed increase in nuclear invagination in HCM (Figure 1A, 1C, 2E, 2G), resulting in many sites of high nuclear membrane curvature, could lead to YAP1 nuclear import ^48^.

Indeed, HCM patients showed an increased YAP1 nuclear localization, compared to NF donors independent of genotype along with a higher YAP1 nuclear and cytoplasmic integrated density indicating higher protein levels of YAP1 per cardiomyocyte with a relatively higher increase in the nucleus (Figure 4A-D). We also observed increased YAP1 nuclear expression in *Mybpc3^insG/insG^*compared to WT mice (Figure S10). To determine whether the increased YAP1 nuclear localization was the result of increased canonical Hippo pathway activity ^49^, western blots for YAP1 and p-S127 YAP1 were performed. There was no difference in YAP1 and relative YAP1 phosphorylation between NF donor and HCM tissue (Figure 4E-G, Figure S11). In addition, we examined whether nuclear invagination length was associated with YAP1 nuclear accumulation. In a subset of donor and HCM samples previously shown to have a high nuclear YAP1 signal (Figure 1C), we quantified normalized nuclear invagination length and correlated this with YAP1 nuclear integrated density. In FFPE tissue, we again observed increased nuclear invaginations in HCM (Figure 4H). Furthermore, normalized invagination length correlated with YAP1 nuclear integrated density (*r*^2^=0.2979, *p*=<0.0001, Figure 4I). This illustrates that the elevated nuclear YAP1 levels observed in HCM are unlikely to be explained by canonical Hippo pathway regulation and instead may relate to altered nuclear mechanics, potentially involving increased nuclear tension.

**Figure 4:**
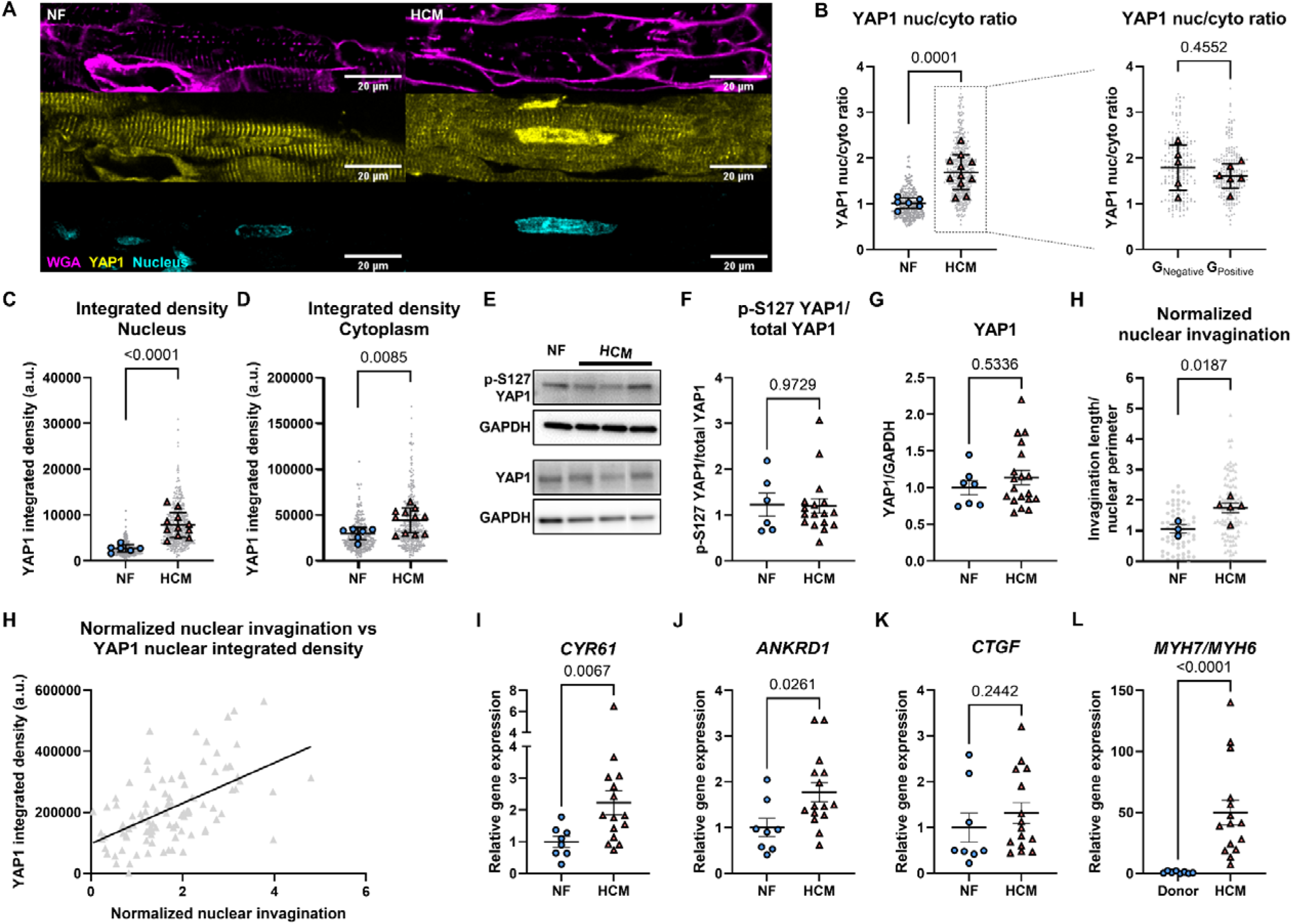
YAP1 localization, protein expression and target gene expression in NF donors and HCM patients. Representative images of intensity and localization of YAP1 in FFPE heart tissue of NF donors and HCM patients (A). Quantification of YAP1 nuclear to cytoplasm ratio in NF donor and HCM patient tissue (left panel) and G_Negative_ and G_Positive_ (right panel). The nuc/cyto ratio was calculated by dividing the mean intensity of YAP1 in the nucleus by the mean intensity of YAP1 in the cytoplasm (B). Quantification of the YAP1 integrated density in the nucleus and cytoplasm (C, D). Representative images of western blots (E). Quantification of the ratio of p-S127 YAP1 and total YAP1 in NF donors and HCM patients. Both p-S127 YAP1 and YAP1 were normalized to GAPDH expression after which the ratio was determined. Every value is the average of two measurements per donor/patient (F). Quantification of YAP1 protein expression normalized to GAPDH protein expression in NF donors and HCM patients. Every value is the average of two measurements per donor/patient (G). Quantification of the normalized nuclear invagination of a subset of donor and HCM samples. Normalized nuclear invagination is calculated by dividing the total invagination length by the nuclear perimeter (H). Simple linear regression of the normalized nuclear invagination and YAP1 nuclear integrated density of a subset of the HCM samples (I, *p*<0.0001, *r*^2^=2979). Quantification of gene expression determined by RT-qPCR of *CYR61* (J), *ANKRD1* (K), *CTGF* (L) and the *MYH7/MYH6* ratio (M). Gene expression is normalized to the housekeeping genes *PSMB2* and *OARD1* J-M). Data are expressed as mean ± standard error of the mean. Every grey symbol represents the value of a single cell and/or nucleus, *n*, and every colored symbol represents the average value per single donor or patient, *N* (B) NF: *N*=6 *n*=281; HCM: *N*=11 *n*=410; G_Negative_ *N*=5 *n*=190; G_Positive_ *N*=6 *n*=220, (C, D) NF: *N*=6 *n*=281; HCM: *N*=11 *n*=410, (F) NF: *N*=6; HCM: *N*=16, (G) NF: *N*=7; HCM: *N*=19, (H) NF: *N*=3 *n*=60; HCM: *N*=5 *n*=100, (I) HCM: *N*=5, *n*=100; (J-M) NF: *N*=8; HCM: *N*=15. Statistical tests: Welch’s t-test (B left panel, C, D, H), unpaired t-test (B right panel, K), Mann-Whitney test (F, G, J, L, M)

To assess if the translocated YAP1 is transcriptionally active, we checked the expression of YAP1 target genes (*CYR61, ANKRD1* and *CTGF*) in HCM tissue by RT-qPCR. The expression of *YAP1*, *CYR61* and *ANKRD1* was increased in HCM tissue (Figure S12A, 4J, 4K), while CTGF expression remained unaltered (Figure 4M). YAP1 also has hypertrophic genes as downstream targets ^46,50,51^. Therefore, we determined their gene expression and showed that *MYH6* gene expression was lower and hypertrophic genes *MYH7, NPPA* and *NPPB* expression was higher (Figure 4M, S12B-E). This suggests that nuclear abnormalities as a result of impaired mechanotransduction lead to nuclear accumulation of YAP1, where it induced a pro-hypertrophic signaling cascade.

### Nuclear-cytoskeletal coupling is identified as the potential culprit of impaired nuclear mechanotransduction in HCM rather than nuclear stiffness

We next sought to identify which cellular components contribute to this lack of nuclear deformability in *Mybpc3^insG/insG^* cardiomyocytes. The stiffness of the nucleus is dependent on the protein levels of Lamin A/C ^52,53^ and chromatin condensation ^54–56^. Furthermore, cytoskeletal stiffness contributes to the deformation of the nucleus ^28,57,58^. Together they determine the plasticity and sensitivity of the nucleus to strain. In *Mybpc3^insG/insG^* cardiomyocytes, there seemed to be a decreased lamin A/C expression and chromatin condensation (Figure 13A, S13B) rather than the expected increase that would reduce deformability, suggesting that the reduced nuclear deformability in *Mybpc3^insG/insG^* cardiomyocytes is unlikely due to increased nuclear stiffness.

Microtubule detyrosination is higher in *Mybpc3^insG/insG^*mice ^15^ and in HCM patients ^9,59,60^. Since microtubules are connected to the nucleus via the LINC complex, we investigated the link between abnormal nuclear morphology and microtubules. Therefore, the microtubule code was assessed in the same patient samples that were used for TEM. HCM patients showed higher α-tubulin, detyrosinated α-tubulin, acetylated α-tubulin and desmin protein expression (Figure 5A-D, 5F, Figure S14-S17). The inner nuclear membrane protein Lamin A/C, which is often a key protein in diseases involving nuclear abnormalities (e.g. laminopathies), was not differently expressed in HCM patients (Figure 5E, 5F, S18).

**Figure 5:**
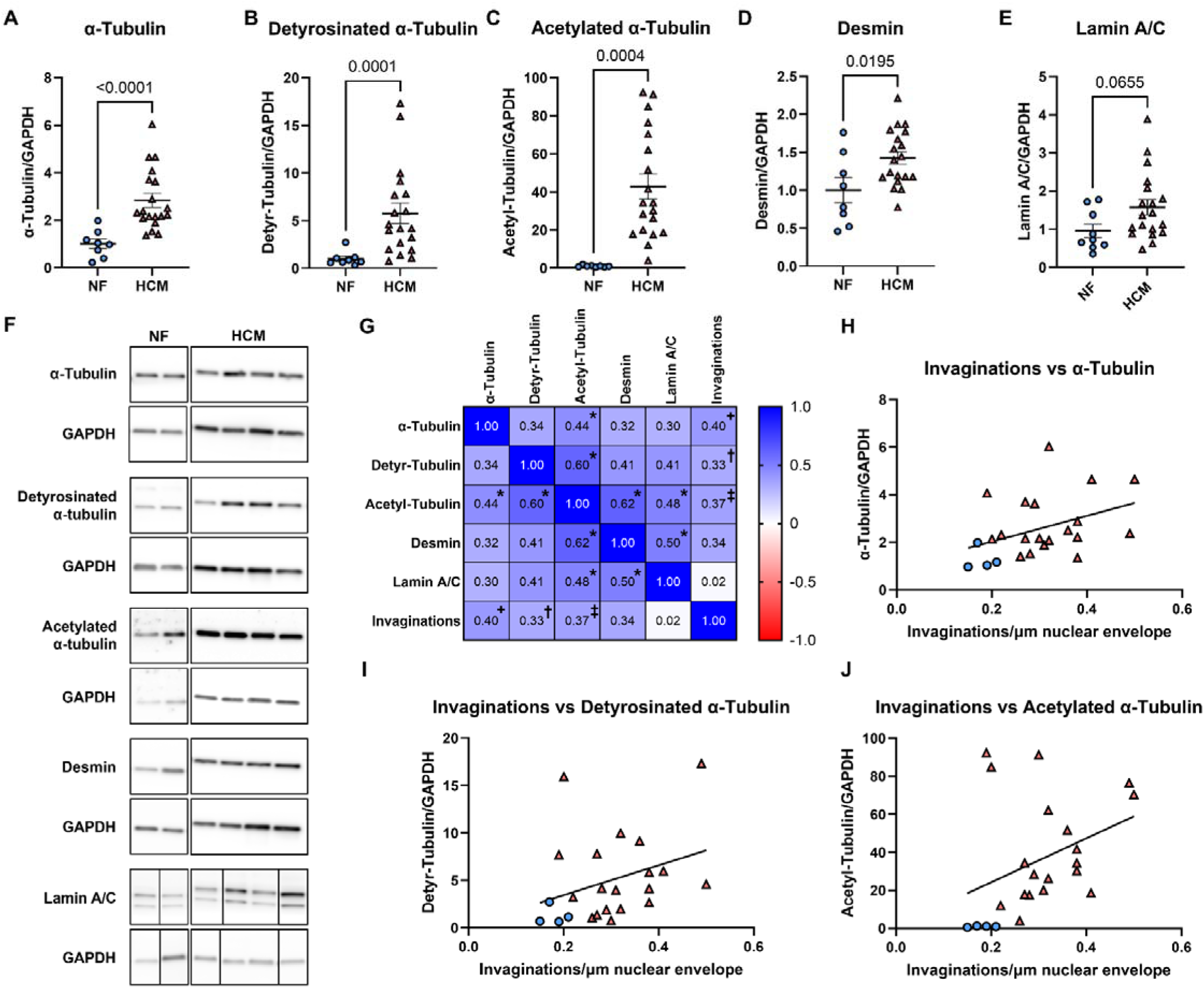
Cytoskeletal protein expression in HCM and its correlation with nuclear parameters. Quantification of protein levels of α-tubulin (A), detyrosinated α-tubulin (B), acetylated α-tubulin (C), desmin (D) and Lamin A/C (E) in NF donors and HCM patients. Protein levels were normalized to GAPDH. Every value is the average of two measurements per donor/patient (A-E). Representative western blots of NF donors and HCM patients (F). Pearson R correlation of non-failing donors and HCM patients (G, *= *p*<0.05, += *p*=0.06, †=*p*= 0.12, ‡= *p*=0.09). Simple linear regression of α-tubulin with nuclear invaginations in non-failing donors and HCM patients (H, *p*=0.06, *r*^2^=0.16). Simple linear regression of detyrosinated α-tubulin with nuclear invaginations in non-failing donors and HCM patients (I, *p*=0.12, *r*^2^=0.11). Simple linear regression of acetylated α-tubulin with nuclear invaginations in non-failing donors and HCM patients (J, *p*=0.09, *r*^2^=0.13). Data are expressed as mean ± standard error of the mean. Every colored symbol represents the average value per single donor or patient, *N*. (A-E, G-J) NF: *N*=8; HCM: *N*=19. Statistical tests: Mann-Whitney test (A,B), unpaired t-test (C-E).

Pearson’s correlation analysis revealed associations between nuclear invagination length and total microtubules, as well as detyrosinated and acetylated microtubules (Figure 5G, Table S19). Linear regression analyses further demonstrated moderate correlations between nuclear invagination length and α-tubulin, detyrosinated α-tubulin, and acetylated α-tubulin levels (Figure 5H–J). When grouping patients by genotype, we observed strong correlations in G_Negative_ patients, whereas correlations were weak to moderate in G_Positive_ patients (Figure S19). This difference may reflect the impact of sarcomere mutations as a contributing parameter in these associations. Together, these findings indicate a link between microtubule remodeling and nuclear invagination formation in HCM patients, highlighting microtubules as a potential therapeutic target to restore impaired nuclear–cytoskeletal coupling.

### Microtubule modifying compounds rescue restricted nuclear deformation in Mybpc3^insG/insG^ *mice*

To show a causal relationship between microtubules and nuclear morphology and deformability, we used microtubule detyrosination inhibitor epoY ^61^ and complete microtubule destabilizer nocodazole. EpoY did not affect resting sarcomere length and sarcomere shortening, whereas it shortened contraction time in both WT and *Mybpc3^insG/insG^* mice and restored impaired relaxation in *Mybpc3^insG/insG^* mice (Figure S20). On the other hand, nocodazole did not affect resting sarcomere length, lowered sarcomere shortening, did not affect contraction kinetics, and also seemed to restore impaired relaxation in *Mybpc3^insG/insG^* mice (Figure S21). EpoY did not affect nuclear size parameters at full relaxation (Figure S22), while nocodazole significantly increased nuclear area (Figure S23).

Treatment with epoY resulted in a similar nuclear area and nuclear length deformation in WT and *Mybpc3^insG/insG^* mice (Figure 6A, 6B). Small changes in nuclear length (Figure 6E) and nuclear width deformation (Figure 6F) led to a significant effect of epoY on nuclear area deformation (Figure 6D), thereby restoring nuclear deformability in *Mybpc3^insG/insg^* mice to similar levels as in WT mice (Figure S24, S25, S26). Restored sarcomere-nuclear coupling can also be observed in the similar strain curves between WT and *Mybpc3^insG/insG^* mice (Figure 6C). EpoY treatment prevented the hysteresis in nuclear length deformation during contraction that was observed in *Mybpc3^insG/insG^* cardiomyocytes (Figure 6G). Nocodazole treatment also resulted in similar nuclear area deformation (Figure 6H), nuclear length deformation (Figure 6I) and strain curves in WT and *Mybpc3^insG/insG^* mice (Figure 6J, S24, S25, S26). There was a significant effect of nocodazole on nuclear length deformation, probably due to decreased sarcomere shortening (Figure 6L). Furthermore, nocodazole also prevented the hysteresis in nuclear length deformability in *Mybpc3^insG/insG^* cardiomyocytes (Figure 6N). However, it is unknown whether the sarcomere-nuclear coupling would also be restored using nocodazole at a similar sarcomere shortening as in untreated cardiomyocytes (Figure S20B). These findings imply that inhibition of detyrosination of microtubules can rescue impaired relaxation and nuclear-cytoskeletal coupling and therefore nuclear deformation.

**Figure 6:**
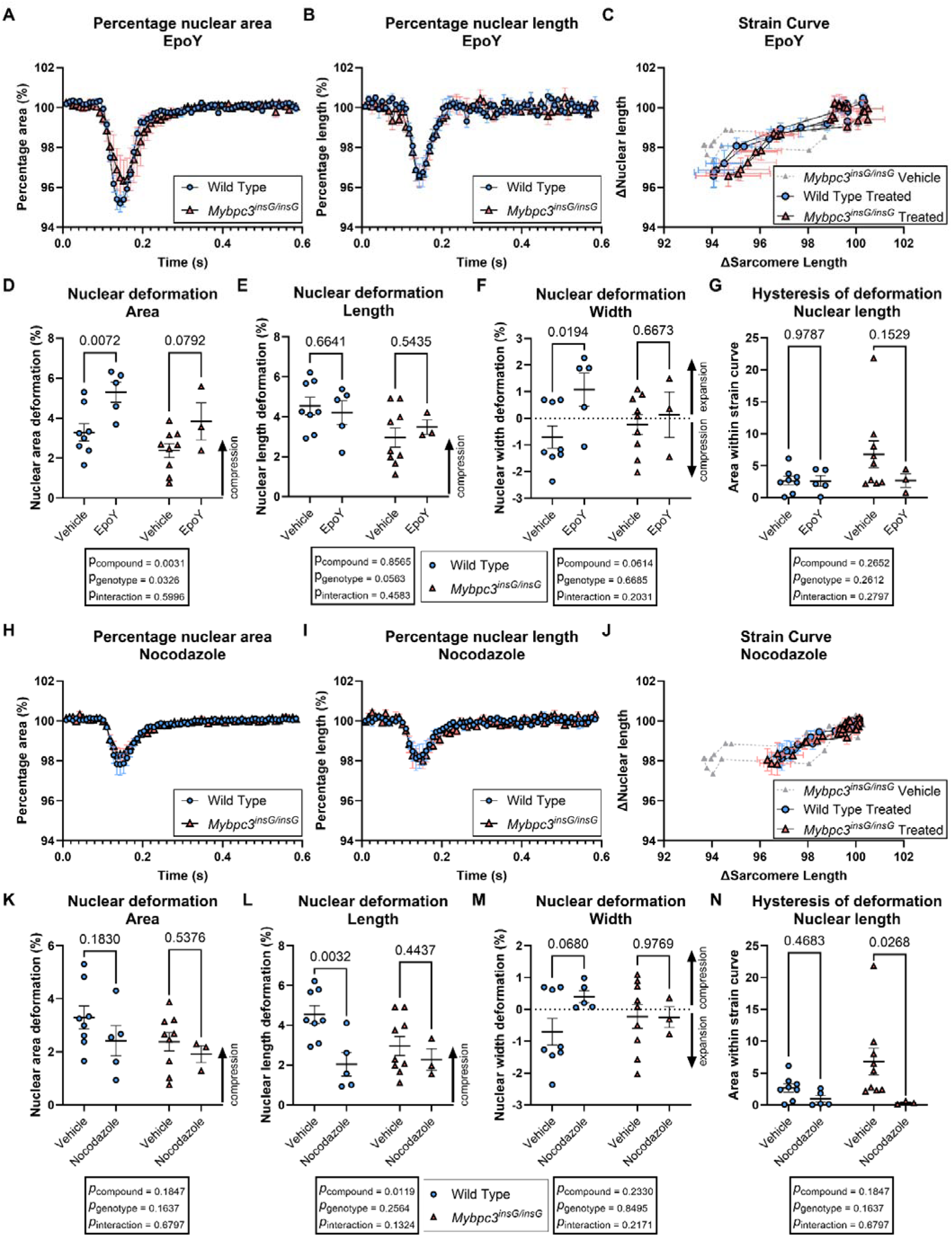
Nuclear deformation during contraction in WT and *Mybpc3^insG/insG^* mice upon the addition of microtubule modifiers nocodazole and epoY. Representation of the nuclear area (A) and nuclear length (B) over time in WT and *Mybpc3^insG/insG^*cardiomyocytes with addition of epoY. Sarcomere-nuclear strain curve generated by calculating the change in sarcomere length versus the change in nuclear length for the addition of epoY (C). Quantification of the maximum nuclear area deformation (D), nuclear length deformation (E), nuclear width deformation (F) and hysteresis in the deformation of the nuclear length during a contraction (G) with the addition of epoY. The hysteresis was determined by calculating the area within the strain curve. Representation of the nuclear area (H) and nuclear length (I) over time in WT and *Mybpc3^insG/insG^*cardiomyocytes with addition of nocodazole. Sarcomere-nuclear strain curve generated by calculating the change in sarcomere length versus the change in nuclear length for the addition of nocodazole (J). Quantification of the maximum nuclear area deformation (K), nuclear length deformation (L), nuclear width deformation (M) and hysteresis in the deformation of the nuclear length during a contraction (N) with the addition of nocodazole. Data are expressed as mean ± standard error of the mean. Every colored symbol represents the value of the average of the mice in their group (A-C, H-J). For the quantifications every colored symbol represents the averaged value per single mouse, N (D-G, K-N). (A, B) *Mybpc3^insG/insG^*, vehicle: *N*=9 *n*=145; *Mybpc3^insG/insG^*, epoY: *N*=3 *n*=29. (C) *Mybpc3^insG/insG^*, vehicle: *N*=9 *n*=145; WT, epoY: *N*=5 *n*=119; *Mybpc3^insG/insG^*, epoY: *N*=3 *n*=29. (D-G) WT, vehicle: *N*=8 *n*=223; WT, epoY: *N*=5 *n*=119; *Mybpc3^insG/insG^*, vehicle: *N*=9 *n*=145; *Mybpc3^insG/insG^*, epoY: *N*=3 *n*=29. (H, I) *Mybpc3^insG/insG^*, vehicle: *N*=9 *n*=145; *Mybpc3^insG/insG^*, nocodazole: *N*=3 *n*=62. (J) *Mybpc3^insG/insG^*, vehicle: *N*=9 *n*=145; WT, nocodazole: *N*=5 *n*=125; *Mybpc3^insG/insG^*, nocodazole: *N*=3 *n*=62. (K-N) WT, vehicle: *N*=8 *n*=223; WT, nocodazole: *N*=5 *n*=125; *Mybpc3^insG/insG^*, vehicle: *N*=9 *n*=145; *Mybpc3^insG/insG^*, nocodazole: *N*=3 *n*=62. Statistical test: two-way ANOVA (D-G, K-N).

### Microtubule detyrosination inhibitor epoY reduces nuclear invagination and induces YAP1 cytoplasmic retention

Since microtubule-modifying compounds can rescue nuclear deformability, we wondered whether they could also reduce nuclear invagination and have an effect on YAP1 localization. We treated isolated mouse cardiomyocytes with epoY for 2 hours and saw a reduced invagination length (Figure 7A). On the other hand, nocodazole treatment did not lead to reduced invagination of the nucleus (Figure 7B). EpoY treatment showed a trend towards lowered YAP1 nuclear to cytoplasm ratio: however, only in the *Mybpc3^insG/insG^*mice (Figure 7C). Nocodazole did not change YAP1 nuclear to cytoplasmic ratio in *Mybpc3^insG/insG^* mice, but led to an increased ratio in WT. Simple linear regression showed a good correlation between nuclear invagination and YAP1 nuclear to cytoplasmic ratio upon epoY treatment (*p*=0.03, *r*^2^=93) in *Mybpc3^insG/insG^* mice (Figure 7E). This correlation is absent upon nocodazole treatment (*p*=0.45, *r*^2^=0.30) (Figure 7F). EpoY and nocodazole do not change other modulators of nuclear stiffness, lamin A/C deposition or chromatin condensation (Figure S27), however, they do alter nuclear morphology by significantly reducing the circularity of the nuclei (Figure S28, S29). Taken together, this demonstrates that in HCM, microtubule detyrosination plays an important role in nuclear cytoskeleton coupling and signaling. Treatment with compounds targeting microtubule detyrosination can rescue nuclear deformation, invagination, and strain, and results in a possible rescue of pathological signaling by transcription factors like YAP1 that depend on nuclear translocation.

**Figure 7:**
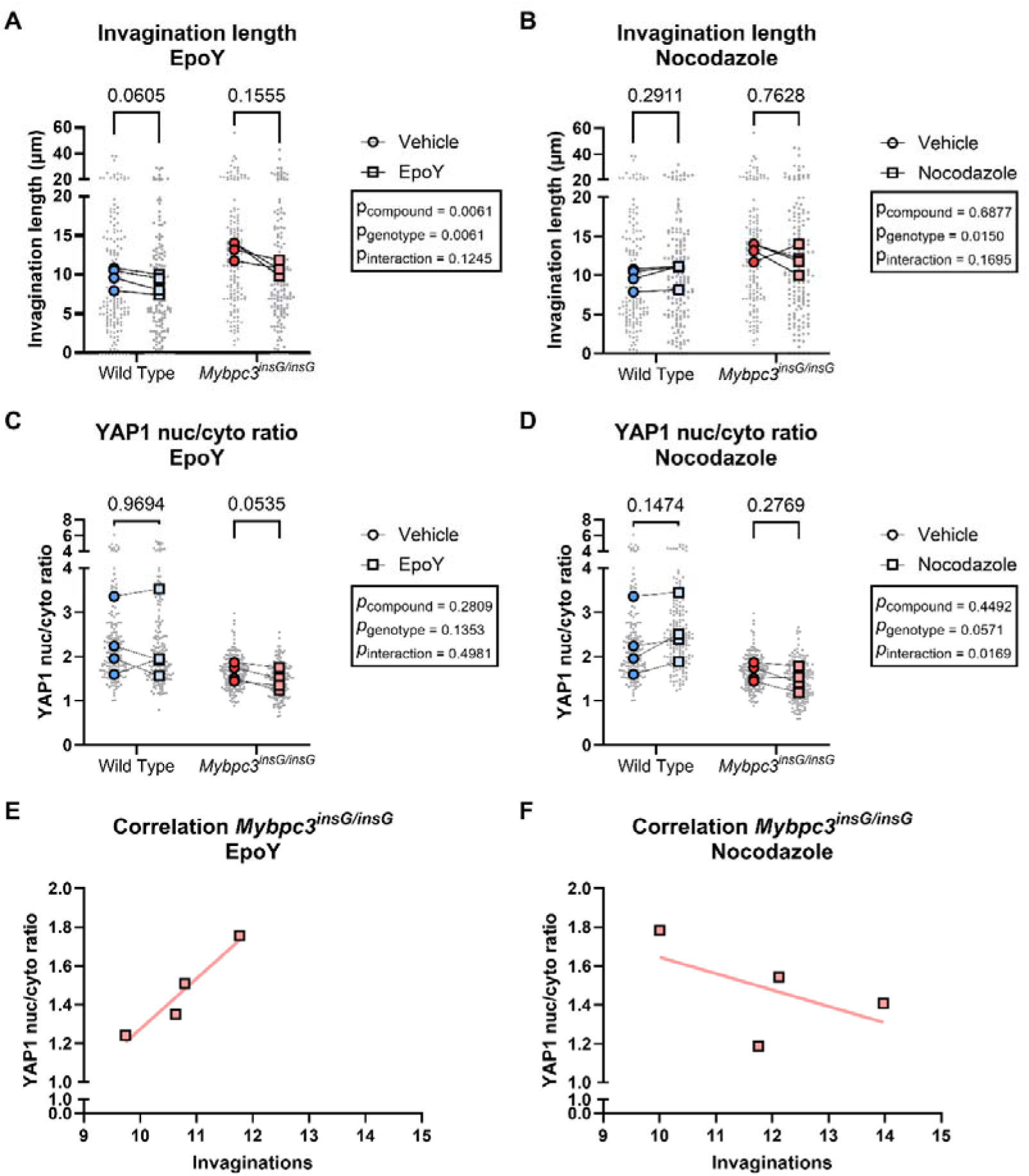
Invagination length and YAP1 nuc/cyto ratio upon the addition of microtubule modifiers nocodazole and epoY. Quantification of invagination length of WT and *Mybpc3^insG/insG^* cardiomyocytes with the addition of vehicle compared to epoY (A). Quantification of invagination length of WT and *Mybpc3^insG/insG^* cardiomyocytes with the addition of vehicle compared to nocodazole (B). Invagination length was calculated as the sum of every separate length of lamin A/C protruding into the nucleus (A, B). Quantification of the ratio of nuclear and cytoplasmic YAP1 in WT and *Mybpc3^insG/insG^* cardiomyocytes with the addition of vehicle compared to epoY (C). Quantification of the ratio of nuclear and cytoplasmic YAP1 in WT and *Mybpc3^insG/insG^* cardiomyocytes with the addition of vehicle compared to nocodazole (D). The nuc/cyto ratio was calculated by dividing the mean intensity of YAP1 in the nucleus by the mean intensity of YAP1 in the cytoplasm (C, D). Simple linear regression of nuclear invaginations and YAP1 nuc/cyto ratio upon epoY treatment in *Mybpc3^insG/insG^* cardiomyocytes (E, *p*=0.03, *r*^2^=0.93). Simple linear regression of nuclear invaginations and YAP1 nuc/cyto ratio upon nocodazole treatment in *Mybpc3^insG/insG^* cardiomyocytes (F, *p*=0.45, *r*^2^=0.30). Data are expressed as mean ± standard error of the mean. Every grey symbol represents the value of a single nucleus, *n*, and every colored symbol represents the average value per single mouse, *N*. Colored symbols are linked when the vehicle and epoY or Nocodazole treated cardiomyocytes come from the same mouse. (A) WT, vehicle: *N*=4 *n*=162; WT, epoY: *N*=4 *n*=157; *Mybpc3^insG/insG^*, vehicle: *N*=4 *n*=151; *Mybpc3^insG/insG^*, epoY: *N*=4 *n*=147, (B) WT, vehicle: *N*=4 *n*=162; WT, nocodazole: *N*=4 *n*=167; *Mybpc3^insG/insG^*, vehicle: *N*=4 *n*=151; *Mybpc3^insG/insG^*, nocodazole: *N*=4 *n*=154, (C) WT, vehicle: *N*=4 *n*=173; WT, epoY: *N*=4 *n*=174; *Mybpc3^insG/insG^*, vehicle: *N*=4 *n*=171; *Mybpc3^insG/insG^*, epoY: *N*=4 *n*=163, (D) WT, vehicle: *N*=4 *n*=173; WT, nocodazole: *N*=4 *n*=177; *Mybpc3^insG/insG^*, vehicle: *N*=4 *n*=171; *Mybpc3^insG/insG^*, nocodazole: *N*=4 *n*=173, (E) *Mybpc3^insG/insG^*, epoY: *N*=4, (F) *Mybpc3^insG/insG^*, nocodazole: *N*=4. Statistical test: two-way ANOVA (A-D).

## Discussion

In this study, we related cytoskeletal changes in HCM to alterations in nuclear morphology, strain and mechanotransduction. We found that HCM patients and a HCM mouse model display hypertrophic, highly invaginated and less deformable nuclei that may create a permissive environment for nuclear translocation of transcription factors, such as YAP1, which might have a considerable impact on hypertrophic signaling. The addition of microtubule destabilizer nocodazole or detyrosination inhibitor epoY restored nuclear deformability to non-diseased levels (Figure 8). Only epoY led to reduced nuclear invagination and reduced YAP1 nuclear levels. Taken together, this is the first report to comprehensively characterize nuclear abnormalities and mechanistically couple microtubule remodeling to impaired mechanotransduction and altered signaling in both HCM patients and a HCM mouse model and demonstrates the therapeutic potential of specifically microtubule detyrosination inhibitors.

The observation that nuclei are less deformable in *Mybpc3^insG/insG^*mice indicates a potential role for nuclear mechanotransduction in the pathogenesis of HCM through an altered strain on the nucleus, which can induce changes in protein conformation, nuclear envelope stretching and chromatin accessibility ^26,62^, and can, therefore, alter downstream signaling ^25,63,64^. However, it is not obvious whether the strain on the nucleus is increased or decreased in HCM based on our results. The upward shift of the strain curve at peak systole suggests a decrease in strain transfer of the sarcomere to the nucleus in *Mybpc3^insG/insG^* mice (Figure 3H)^65^. In addition, we also observed an increase in nuclear invagination, which could indicate that the nucleus is collapsing in association with reduced nuclear tensile strain ^66^. On the contrary, an alternative hypothesis suggests there is an increase of compressive strain on the nucleus by the protrusion of microtubules into the nucleus leading to nuclear envelope stretching ^25,47,66^. Using advanced imaging techniques, recent research has also shown that deep nuclear invaginations are linked to cytoskeletal filaments ^67^. In addition, it has been demonstrated that areas of higher membrane curvature encounter higher nuclear strain and increased active and passive nuclear import through nuclear pores ^48^. In that case, it would be expected that YAP1 is translocated to the nucleus ^45,46,68^. We indeed observed that YAP1, a protein involved in hypertrophic signaling via the Hippo pathway ^45,46^, was translocated to the nucleus in HCM patients, inducing transcription of downstream target genes. This was in line with another study where human HCM tissue was assessed on YAP expression ^46^. Consequently, we showed that returning sarcomere-nuclear strain transfer to non-diseased levels (Figure 6E), resulted in a reduction of nuclear invagination and YAP1 translocation to the nucleus (Figure 7A). All in all, this suggests that the strain on the nucleus is more likely increased in HCM. Further studies using for example a nuclear strain sensor ^69^ may aid in unraveling the strain discrepancy.

Even though nuclear deformability is dependent on multiple factors, such as chromatin condensation ^55,64^, lamin deposition ^70^ and the cytoskeletal network, in our model nuclear mechanics appeared to be dominated by detyrosinated α-tubulin, as treatment of cardiomyocytes with the microtubule detyrosination inhibitor epoY fully rescued nuclear deformability. Forces generated by sarcomere contraction are known to be transmitted to the nucleus through a surrounding microtubule cage ^71^. Detyrosinated microtubules form more stable attachments to the sarcomere than tyrosinated microtubules and act as compression-resistant elements that buckle less readily ^12^. In line with this, our strain measurements revealed that after approximately one-third of total sarcomere shortening the nucleus in *Mybpc3^insG/insG^* cardiomyocytes no longer shortened further (Figure 3H). We propose that stiff, detyrosinated microtubules tethered to the sarcomere reach a mechanical limit beyond which further buckling is prevented, thereby mechanically buffering the nucleus and restricting additional global nuclear deformation detectable with our approach. In this scenario, contractile energy would be absorbed by the microtubule network and subsequently released during diastole, explaining the faster nuclear re-lengthening observed in mutant cells. Although this buffering may reduce measurable whole-nucleus deformation, it could simultaneously generate localized envelope curvature and strain at sites of microtubule-nuclear contact that promote nuclear import of mechanosensitive factors such as YAP1, suggesting that our strain measurements may underestimate the local nuclear stresses driving downstream signaling. These findings are consistent with previous reports showing that hypertrophic cardiomyocytes exhibit elevated viscosity and stiffness due to detyrosinated α-tubulin ^13,72^, leading to altered mechanotransduction ^22^. Moreover, detyrosinated microtubules impair relaxation, and inhibition of detyrosination improves contractility ^9^. While detyrosinated α-tubulin has already been proposed as a therapeutic target because of its beneficial effects on contractile dysfunction, our data further suggest that modulating microtubule post-translational modifications may also restore aberrant sarcomere-nuclear mechanotransduction and nuclear signaling in HCM.

Beyond its effects on global nuclear mechanics, microtubule remodeling also emerged as a key determinant of YAP1 localization. We identified a strong association between microtubule abundance and nuclear invagination, as well as between nuclear invagination and YAP1 nuclear accumulation. Moreover, pharmacological reduction of microtubule detyrosination prevented YAP1 nuclear enrichment, suggesting that microtubule-dependent nuclear remodeling contributes to YAP1 localization in HCM cardiomyocytes. We therefore hypothesized that YAP1 nuclear entry may be facilitated by increased nuclear curvature and deformation associated with invaginations ^48^. However, YAP1 localization is likely regulated by multiple, intersecting mechanotransduction pathways. Consistent with this concept, Vanni et al. ^73^ recently demonstrated that microtubules modulate YAP1 nuclear accumulation through regulation of angiomotin (AMOT) stability. In their study, AMOT functioned as a mechanosensitive cytoplasmic anchor for YAP1. Under conditions of low nuclear strain, AMOT sequestered YAP1 and prevented its nuclear translocation, whereas increased nuclear strain promoted AMOT degradation, thereby permitting YAP1 nuclear accumulation. Importantly, AMOT stability depended on intact and mechanically coupled microtubules. Disruption of microtubule–nuclear envelope connections or pharmacological destabilization of microtubules preserved AMOT and prevented YAP1 nuclear accumulation. Taken together with our findings, these observations support a model in which microtubules regulate YAP1 localization through at least two complementary mechanisms. First, microtubule-dependent forces acting on the nucleus may drive nuclear deformation and invagination, increasing nuclear tension and curvature that facilitate YAP1 import. Second, microtubules may regulate YAP1 indirectly by controlling AMOT turnover in response to nuclear strain. Thus, altered microtubule organization and post-translational modification in HCM could promote sustained YAP1 nuclear accumulation by simultaneously remodeling nuclear architecture and modulating cytoplasmic retention pathways, thereby providing a mechanistic link between cytoskeletal remodeling, nuclear architecture, and maladaptive transcriptional reprogramming in HCM.

Our analysis revealed that not only cardiomyocytes are hypertrophic, but also cardiomyocyte nuclei are hypertrophic and show nuclear invaginations in our HCM mouse model. This is in line with previous research that has shown nuclear hypertrophy and abnormal nuclear shapes in different cardiomyopathies, including HCM ^29^. Polyploidy has been proposed as a mechanism to induce nuclear hypertrophy ^40,74–76^. The correlation between nuclear size and DNA content found in this study supports this theory. In HCM, there is a change in mechanical workload to meet its increased demands. Cardiomyocytes respond to this increase in demand with expansion in size through the addition of sarcomeres. This suggests that the increased ploidy observed in our study provides the cell with its required increase in biosynthetic activities, such as transcription, translation, and DNA repair and synthesis ^29,40,77^.

## Study limitations

A limitation of this study was that nuclear characterization was only done at one specific point in the development of HCM. This included septal myectomy tissue of HCM patients and *Mybpc3^insG/insG^*mice of 4-6 months of age, which both resemble overt HCM. It is, however, very relevant to study HCM at different stages of the disease to be able to unravel a causal relationship between nuclear abnormalities and the development of HCM. Due to the inability to obtain patient tissue at earlier stages of the disease, this cannot be studied in humans. The mouse model, on the other hand, has shown to possess stage II characteristics of HCM at 3-4 weeks of age ^15^ and would, therefore, be an ideal model to investigate causality. Furthermore, only short-term effects of epoY and Nocodazole have been assessed. In future research, it would be interesting to see if modulating microtubules in the same mouse model could potentially alter mechanosensing pathways and their downstream effects, such as hypertrophy and impaired relaxation, in the long term without encountering adverse effects.

## Conclusion

Overall, extensive nuclear changes are observed in the HCM mouse model. Cytoskeletal remodeling results in altered sarcomere-nuclear coupling and nuclear invaginations that might contribute to impaired mechanotransduction. Nuclear deformability and invagination can be restored by microtubule modifying drugs, specifically those that decrease microtubule detyrosination. Our study provides proof for the involvement of dysregulated mechanotransduction in HCM and that intervening through the inhibition of detyrosination of microtubules or targeting downstream signaling pathways might be a promising therapeutic strategy.

## Clinical perspectives

### Core clinical competencies

Hypertrophic cardiomyopathy (HCM) is characterized by left ventricular hypertrophy and diastolic dysfunction and is caused by sarcomere mutations in approximately half of patients (genotype-positive), while the etiology in genotype-negative patients remains unclear. The disease mechanisms of HCM is still incompletely understood. This study identifies abnormal nuclear morphology and impaired nuclear mechanotransduction in cardiomyocytes from both genotype-positive and genotype-negative HCM patients, associated with cytoskeletal remodeling. These nuclear alterations were linked to enhanced nuclear localization of the mechanosensitive transcription factor YAP1 and hypertrophic gene expression, suggesting that disrupted cytoskeleton–nucleus coupling represents an important pathogenic pathway in HCM.

### Translational outlook

Targeting cytoskeletal remodeling in cardiomyocytes may represent a novel therapeutic strategy for hypertrophic cardiomyopathy. This study demonstrates that pharmacological reduction of microtubule detyrosination attenuates abnormal nuclear morphology and decreases nuclear localization of the mechanosensitive transcription factor YAP1. Future investigations employing sustained modulation of detyrosinated tubulin levels in preclinical models are needed to determine whether this approach can improve cardiac hypertrophy, diastolic function, and overall myocardial performance on the long-term.

## Acknowledgments

We thank the Core Facility Cellular Imaging and the Microscopy Unit MCBI of the Amsterdam UMC for technical assistance, use of equipment and advise. Figures were created with BioRender.com.

## Author contributions

ID designed and performed experiments, analyzed and interpreted data and drafted the manuscript. LRM designed and performed experiments and analyzed data. EEN and VJ performed experiments. JAM and JSB performed experiments and analyzed data. KB and KBM collected, processed and provided non-failing donor material. SACS and MM collected and provided patient material. NNW assisted TEM experiments. JV conceptualized research. TJK and DWDK conceptualized research, interpreted data and supervised the project. All authors critically read, edited and approved the manuscript.

## Data availability

Data is available upon reasonable request. Correspondence and requests for materials or data should be addressed to TJK and DWDK.

## Abbreviations

HCM: Hypertrophic cardiomyopathy
G_Positive_: Genotype positive
G_Negative_: Genotype negative
LV: left ventricle
LINC: Linker of Nucleoskeleton and Cytoskeleton
YAP1: Yes-associated protein 1
TEM: Transmission electron microscopy
BSA: Bovine serum albumin
TBS-T: Tris-buffered Saline with 0.1 (v/v%) Tween
*Mybpc3^+/InsG^* /*Mybpc3^InsG/^*^InsG^: heterozygous/homozygous MYBPC3_c*.2373InsG*_ mouse model

