## Supplemental data for "Microtubule detyrosination alters nuclear mechanotransduction and leads to pro-hypertrophic signaling in hypertrophic cardiomyopathy"

**Supplemental appendix**

Table S1	Patient characteristics of tissue used	p. 3

Table S2	Comparison of clinical characteristics of non-failing donor and hypertrophic cardiomyopathy patients to show equality of groups for transmission electron	microscopy in figure 1.	P. 3

Table S3	Comparison of clinical characteristics of non-failing donor and hypertrophic cardiomyopathy patients to show equality of groups for immunohistological	stainings in figure 1 and 4.	P. 4

Table S4	Comparison of clinical characteristics of non-failing donor and hypertrophic cardiomyopathy patients to show equality of groups for qRT-PCRs figure 4.	P. 5

Table S5	Comparison of clinical characteristics of non-failing donor and hypertrophic cardiomyopathy patients to show equality of groups for western blots in	figure 5.	P. 6

Table S6	Overview antibody concentrations used for western blots.	P. 7

Table S7	Overview primer sequences qRT-PCR.	P. 8

Table S8	Characteristics of mice used for embedding of whole mouse hearts.	P. 9

Table S9	Comparison of characteristics of WT, Mybpc3^+/insG^ and Mybpc3^insG/insG^ mice	to show equality of groups for general parameters and the consistency of	the model.	P. 9

Table S10	Characteristics of mice used for cardiomyocyte isolation.	P. 9

Table S11	Overview antibodies used for immunofluorescent staining.	P. 9

Figure S1	Nuclear morphology in human patients with hypertrophic cardiomyopathy.	P. 10

Figure S2	Cell and nuclear morphology in male and female patients with hypertrophic	cardiomyopathy.	P. 11

Table S12	Uncorrected *p*-values of the Pearson R correlation of nuclear invaginations	with several clinical parameter in Figure 1F.	P. 12

Figure S3	2D nuclear morphology of WT, *Mybpc3^+/insG^* and *Mybpc3^insG/insG^* mice.	P. 13

Figure S4	3D nuclear morphology of the WT, *Mybpc3^+/insG^* and *Mybpc3^insG/insG^* mouse	model.	P. 14

Figure S5	Correlations of 2D nuclear parameters with clinical parameters of WT,	*Mybpc3^+/insG^* and *Mybpc3^insG/insG^* mice.	P. 15

Table S13	Uncorrected *p*-values of the Pearson R correlation of 2D nuclear parameters	with clinical parameters of WT, *Mybpc3^+/insG^* and *Mybpc3^insG/insG^* mice in	Figure S5.	P. 16

Figure S6	Correlations of 3D nuclear parameters with clinical parameters of WT,	*Mybpc3^+/insG^* and *Mybpc3^insG/insG^* mice.	P. 17

Table S14	Uncorrected p-values of the Pearson R correlation of 3D nuclear parameters	with clinical parameters of WT, Mybpc3^+/insG^ and Mybpc3^insG/insG^ mice in Figure	S6.	P. 18

Figure S7	The relationship between nuclear size and nuclear invagination.	P. 19

Table S15	Uncorrected *p*-values of the Pearson R correlation between nuclear	morphology and invagination parameters in WT mice in Figure S7A.	P. 20

Table S16	Uncorrected *p*-values of the Pearson R correlation between nuclear	morphology and invagination parameters in *Mybpc3^insG/insG^* mice in Figure S7B.	P. 20

Figure S8	Contraction and relaxation time of isolated cardiomyocytes of WT and	*Mybpc3^insG/insG^* mice.	P. 21

Figure S9	Correlations of nuclear parameters with deformability.	P. 22

Table S17	Uncorrected *p*-values of the Pearson R correlation of nuclear parameters	with deformability in WT mice in Figure S9A.	P. 23

Table S18	Uncorrected *p*-values of the Pearson R correlation of nuclear parameters	with deformability in *Mybpc3^insG/insG^* mice in Figure S9B.	P. 24

Figure S10	YAP localization and protein levels of FFPE heart tissue of WT and	*Mybpc3^insG/insG^* mice.	P. 25

Figure S11	Images of western blots p-S127 YAP1 and YAP1.	P. 26

Figure S12	Relative gene expression *YAP1*, *NPPA*, *NPPB*, *MYH6* and *MYH7*.	P. 27

Figure S13	Lamin A/C deposition and chromatin condensation of WT and *Mybpc3^insG/insG^* cardiomyocytes.	P. 28

Figure S14	Images of western blots of α-tubulin.	P. 29

Figure S15	Images of western blots of detyrosinated α-tubulin.	P. 29

Figure S16	Images of western blots of acetylated α-tubulin.	P. 29

Figure S17	Images of western blots of desmin.	P. 30

Figure S18	Images of western blots of Lamin A/C.	P. 30

Table S19	Uncorrected *p*-values of the Pearson R correlation of the microtubule code	with nuclear invaginations in non-failing donors and HCM patients in Figure	5G.	P. 30

Figure S19	Cytoskeletal protein expression in NF, G_Negative_ and G_Positive_ tissue and its	correlation with nuclear invaginations.	P. 31

Figure S20	Contraction and relaxation time of isolated cardiomyocytes of WT and	*Mybpc3^insG/insG^* mice upon addition of epoY.	P. 33

Figure S21	Contraction and relaxation time of isolated cardiomyocytes of WT and	*Mybpc3^insG/insG^* mice upon addition of nocodazole.	P. 34

Figure S22	Nuclear morphology at full relaxation upon addition of microtubule modifier	epoY.	P. 35

Figure S23	Nuclear morphology at full relaxation upon addition of microtubule modifier	nocodazole.	P. 36

Figure S24	Nuclear deformation percentage curves during contraction in WT and	*Mybpc3^insG/insG^* cardiomyocytes with vehicle or microtubule modifying	compounds.	P. 37

Figure S25	Strain curves of nuclear area and nuclear width of WT versus *Mybpc3^insG/insG^* cardiomyocytes.	P. 38

Figure S26	Strain curves of nuclear area, nuclear length and nuclear width of Vehicle	versus epoY versus Nocodazole in WT or Mybpc3^insG/insG^ cardiomyocytes.	P. 39

Figure S27	Lamin A/C deposition and chromatin condensation upon the addition of	microtubule modifiers nocodazole and epoY.	P. 40

Figure S28	Cell area and nuclear morphology upon the addition of microtubule modifier	epoY.	P. 41

Figure S29	Cell area and nuclear morphology upon the addition of microtubule modifier nocodazole.	P. 42

Table S1: Patient characteristics of tissue used. See separate excel sheet.

Table S2: Comparison of clinical characteristics of non-failing donor and hypertrophic cardiomyopathy patients to show equality of groups for transmission electron microscopy in figure 1. BSA, body surface area; IVS, interventricular septum thickness; LAD, left atrial diameter; LVPW, left ventricular posterior wall thickness; LVOTg, left ventricular outflow tract gradient.

|  | **NF** (***N*=4)** | **HCM all patients** **(*N*=19)** | ***P* value** |
| --- | --- | --- | --- |
| Sex, male | 25.0% (*N*=1) | 68.4% (*N*=13) | 0.11 |
| Age at myectomy | 39.0 ± 23.3 | 55.9 ± 18.4 | 0.11 |
| BSA (m²) | 1.70 ± 0.57 | 1.96 ± 0.22 | 0.13 |
|  | **Gnegative (*N*=9)** | **Gpostive (*N*=10)** | ***P* value** |
| **General** | | | |
| Mutation | - | *MYBPC3*: *N*=5 | - |
|  |  | *MYH7*: *N*=2 |  |
|  |  | *MYH6*: *N*=1 |  |
|  |  | *TNNT2*: *N*=1 |  |
|  |  | *MYL2*: *N*=1 |  |
| Sex, male | 66.7% (*N*=6) | 70.0% (*N*=7) | 0.88 |
| Age at myectomy | 63.9 ± 13.8 | 48.7 ± 19.7 | 0.04 |
| BSA (m²) | 1.89 ± 0.21 | 2.01 ± 0.23 | 0.41 |
| **Dimensions** | | | |
| IVS (mm) | 18.7 ± 3.2 | 22.8 ± 6.8 | 0.12 |
| LAD (mm) | 46.2 ± 6.8 | 46.2 ± 8.2 | >0.99 |
| LVPW (mm) | 10.4 ± 1.6 | 11.3 ± 3.0 | 0.46 |
| **Diastolic parameters** | | | |
| E/A | 1.29 ± 0.41 | 1.44 ± 0.61 | 0.54 |
| E/e' | 21.3 ± 7.3 | 16.5±6.3 | 0.15 |
| Decel. Time (ms) | 249 ± 87 | 242 ± 33 | 0.61 |
| **Obstruction parameters** | | | |
| Rest LVOTg (mmHG) | 79.4 ± 48.2 | 46.7 ± 31.8 | 0.10 |
| Provoked LVOTg (mmHG) | 77.2 ± 37.0 | 66.7 ± 20.4 | 0.56 |
| **Medication** | | | |
| β-blocker | 66.7% (*N*=6) | 80.0% (*N*=8) | 0.51 |
| Calcium channel blocker | 33.3% (*N*=3) | 30.0% (*N*=3) | 0.88 |
| Statins | 11.1% (*N*=1) | 20.0% (*N*=1) | 0.60 |

Table S3: Comparison of clinical characteristics of non-failing donor and hypertrophic cardiomyopathy patients to show equality of groups for immunohistological stainings in figure 1 and 4. BSA, body surface area; IVS, interventricular septum thickness; LAD, left atrial diameter; LVPW, left ventricular posterior wall thickness; LVOTg, left ventricular outflow tract gradient.

|  | **NF** (***N*=6)** | **HCM all patients** (***N*=11)** | ***P* value** |
| --- | --- | --- | --- |
| Sex, male | 33.3% (*N*=2) | 45.5% (*N*=5) | 0.63 |
| Age at myectomy | 40.3 ± 25.2 | 44.8 ± 20.5 | 0.70 |
| BSA (m²) | 1.68 ± 0.46 | 1.89 ± 0.39 | 0.33 |
|  | **Gnegative (*N*=5)** | **Gpostive (*N*=6)** | ***P* value** |
| **General** | | | |
| Mutation | - | MYBPC3: *N*=2 | - |
|  |  | MYH7: *N*=3 |  |
|  |  | MYL2: *N*=1 |  |
| Sex, male | 80.0% (*N*=4) | 16.7% (*N*=1) | 0.04***** |
| Age at myectomy | 48.2 ± 23.1 | 42.0 ± 19.9 | 0.64 |
| BSA (m²) | 1.98 ± 0.58 | 1.82 ± 0.17 | 0.17 |
| **Dimensions** | | | |
| IVS (mm) | 20.5 ± 3.9 | 21.2 ± 4.5 | 0.76 |
| LAD (mm) | 45.1 ± 12.2 | 46.2 ± 4.5 | 0.85 |
| LVPW (mm) | 19.4 ± 13.2 | 11.8 ± 2.6 | 0.24 |
| **Diastolic parameters** | | | |
| E/A | 1.26 ± 0.38 | 1.64 ± 1.36 | 0.84 |
| E/e' | 21.2 ± 8.5 | 20.2 ± 7.7 | 0.84 |
| Decel. Time (ms) | 281.8 ± 58.6 | 236.4 ± 103.2 | 0.41 |
| **Obstruction parameters** | | | |
| Rest LVOTg (mmHG) | 73.2 ± 29.4 | 90.2 ± 45.8 | 0.49 |
| Provoked LVOTg (mmHG) | N too small | | |
| **Medication** | | | |
| β-blocker | 80.0% (*N*=4) | 100.0% (*N*=6) | 0.25 |
| Calcium channel blocker | 20.0% (*N*=1) | 0.0% (*N*=0) | 0.25 |
| Statins | 20.0% (*N*=1) | 33.3% (*N*=2) | 0.62 |

Table S4: Comparison of clinical characteristics of non-failing donor and hypertrophic cardiomyopathy patients to show equality of groups for qRT-PCRs figure 4. BSA, body surface area; IVS, interventricular septum thickness; LAD, left atrial diameter; LVPW, left ventricular posterior wall thickness; LVOTg, left ventricular outflow tract gradient.

|  | **NF** (***N*=8)** | **HCM all patients** (***N*=15)** | ***P* value** |
| --- | --- | --- | --- |
| Sex, male | 50.0% (*N*=4) | 53.3% (*N*=8) | 0.88 |
| Age at myectomy | 54.4 ± 10.5 | 48.1 ± 17.4 | 0.36 |
| BSA (m²) | 1.99 ± 0.24 | 1.96 ± 0.26 | 0.80 |
|  | **Gnegative (*N*=5)** | **Gpostive (*N*=10)** | ***P* value** |
| **General** | | | |
| Mutation | - | MYBPC3: *N*=5 | - |
|  |  | MYH7: *N*=4 |  |
|  |  | MYL2: *N*=1 |  |
| Sex, male | 60.0% (*N*=3) | 50.0% (*N*=5) | 0.71 |
| Age at myectomy | 59.4 ± 7.8 | 42.4 ± 18.3 | 0.07 |
| BSA (m²) | 2.12 ± 0.30 | 1.88 ± 0.20 | 0.09 |
| **Dimensions** | | | |
| IVS (mm) | 20.5 ± 3.9 | 24.2 ± 6.4 | 0.33 |
| LAD (mm) | 47.4 ± 9.6 | 44.4 ± 4.7 | 0.42 |
| LVPW (mm) | 13.3 ± 3.3 | 11.9 ± 2.6 | 0.43 |
| **Diastolic parameters** | | | |
| E/A | 1.20 ± 0.49 | 1.72 ± 1.00 | 0.44 |
| E/e' | 20.0 ± 9.0 | 18.5 ± 7.8 | 0.76 |
| Decel. Time (ms) | 267.0 ± 60.5 | 245.0 ± 79.4 | 0.61 |
| **Obstruction parameters** | | | |
| Rest LVOTg (mmHG) | 77.0 ± 35.3 | 71.3 ± 47.3 | 0.82 |
| Provoked LVOTg (mmHG) | N too small | | |
| **Medication** | | | |
| β-blocker | 80.0% (*N*=4) | 100.0% (*N*=10) | 0.14 |
| Calcium channel blocker | 20.0% (*N*=1) | 10.0% (*N*=1) | 0.59 |
| Statins | 40.0% (*N*=2) | 20.0% (*N*=2) | 0.41 |

Table S5: Comparison of clinical characteristics of non-failing donor and hypertrophic cardiomyopathy patients to show equality of groups for western blots in figure 5. BSA, body surface area; IVS, interventricular septum thickness; LAD, left atrial diameter; LVPW, left ventricular posterior wall thickness; LVOTg, left ventricular outflow tract gradient.

|  | **NF** (***N*=8)** | **HCM all patients** **(*N*=19)** | ***P* value** |
| --- | --- | --- | --- |
| Sex, male | 37.5% (*N*=3) | 68.4% (*N*=13) | 0.14 |
| Age at myectomy | 45.5 ± 23.4 | 55.9 ± 18.4 | 0.18 |
| BSA (m²) | 1.75 ± 0.46 | 1.96 ± 0.22 | 0.13 |
|  | **Gnegative (*N*=9)** | **Gpostive (*N*=10)** | ***P* value** |
| **General** | | | |
| Mutation | - | *MYBPC3*: *N*=5 | - |
|  |  | *MYH7*: *N*=2 |  |
|  |  | *MYH6*: *N*=1 |  |
|  |  | *TNNT2*: *N*=1 |  |
|  |  | *MYL2*: *N*=1 |  |
| Sex, male | 66.7% (*N*=6) | 70.0% (*N*=7) | 0.88 |
| Age at myectomy | 63.9 ± 13.8 | 48.7 ± 19.7 | 0.04 |
| BSA (m²) | 1.89 ± 0.21 | 2.01 ± 0.23 | 0.41 |
| **Dimensions** | | | |
| IVS (mm) | 18.7 ± 3.2 | 22.8 ± 6.8 | 0.12 |
| LAD (mm) | 46.2 ± 6.8 | 46.2 ± 8.2 | >0.99 |
| LVPW (mm) | 10.4 ± 1.6 | 11.3 ± 3.0 | 0.46 |
| **Diastolic parameters** | | | |
| E/A | 1.29 ± 0.41 | 1.44 ± 0.61 | 0.54 |
| E/e' | 21.3 ± 7.3 | 16.5±6.3 | 0.15 |
| Decel. Time (ms) | 249 ± 87 | 242 ± 33 | 0.61 |
| **Obstruction parameters** | | | |
| Rest LVOTg (mmHG) | 79.4 ± 48.2 | 46.7 ± 31.8 | 0.10 |
| Provoked LVOTg (mmHG) | 77.2 ± 37.0 | 66.7 ± 20.4 | 0.56 |
| **Medication** | | | |
| β-blocker | 66.7% (*N*=6) | 80.0% (*N*=8) | 0.51 |
| Calcium channel blocker | 33.3% (*N*=3) | 30.0% (*N*=3) | 0.88 |
| Statins | 11.1% (*N*=1) | 20.0% (*N*=1) | 0.60 |

Table S6: Overview antibody concentrations used for western blots.

| **Target** | **Isotype/host** | **Company/product number** | **Dilution** | **Blocking reagent** |  | **Secondary antibody** | **Dilution** |
| --- | --- | --- | --- | --- | --- | --- | --- |
| GAPDH | Rabbit | Cell Signaling #2118 | 1:5000 | 5% (w/v) milk |  | Goat anti-rabbit IgG-HRP, Dako, P0448 | 1:5000 |
| α-tubulin | Mouse | Sigma-Aldrich T9026 | 1:10000 | 5% (w/v) milk |  | Goat anti-mouse IgG-HRP, Dako, P0447 | 1:2000 |
| Detyrosinated tubulin | Rabbit | Abcam ab48389 | 1:1000 | 5% (w/v) milk |  | Goat anti-rabbit IgG-HRP, Dako, P0448 | 1:2000 |
| Acetylated tubulin | Mouse | Sigma-Aldrich T7451 | 1:20000 | 5% (w/v) milk |  | Goat anti-mouse IgG-HRP, Dako, P0447 | 1:2000 |
| Desmin | Rabbit | Cell Signaling #5332 | 1:1000 | 3% (w/v) BSA |  | Goat anti-rabbit IgG-HRP, Dako, P0448 | 1:5000 |
| Lamin A/C | Rabbit | Abcam ab108595 | 1:5000 | 5% (w/v) milk |  | Goat anti-rabbit IgG-HRP, Dako, P0448 | 1:2000 |
| YAP1 | Rabbit | Invitrogen PA1-46189 | 1:1000 | 5% (w/v) milk |  | Goat anti-rabbit IgG-HRP, Dako, P0448 | 1:5000 |
| p-S127 YAP1 | Rabbit | Antibodies.com A94631 | 1:500 | 3% (w/v) BSA |  | Goat anti-rabbit IgG-HRP, Dako, P0448 | 1:5000 |

Table S7: Overview primer sequences qRT-PCR.

| **Gene** | | **Sequence (5’-3’)** | **Annealing temperature (°C)** |
| --- | --- | --- | --- |
| *YAP1* | Forward | CCTTCTTCAAGCCGCCGGAG | 72.2 |
|  | Reverse | CAGTGTCCCAGGAGAAACAG | 62.4 |
| *CTGF* | Forward | CCTGCAGGCTAGAGAAGCA | 63.5 |
|  | Reverse | GATGCACTTTTTGCCCTTCT | 63.1 |
| *CYR61* | Forward | AGCCTCGATCCTATACAACC | 64.1 |
|  | Reverse | TTCTTTCACAAGGCGGCATC | 68.6 |
| *ANKRD1* | Forward | ATGTGGCGGTGAGGACTGG | 68.7 |
|  | Reverse | GTCGGATCATCTTATAGCGGTTCAG | 67.4 |
| *MYH6* | Forward | GCCCTTTGACATTCGCACTG | 67.9 |
|  | Reverse | GGTTTCAGCAATGACCTTGCC | 67.7 |
| *MYH7* | Forward | GGAGTTCACACGCCTCAAAGAGG | 70.1 |
|  | Reverse | TCCTCAGCATCTGCCAGGTTGT | 70.3 |
| *NPPA* | Forward | CCGTGAGCTTCCTCCTTTTA | 63.2 |
|  | Reverse | CCAAATGGTCCAGCAAATTC | 64.1 |
| *NPPB* | Forward | GCTTTGGGAGGAAGATGGAC | 64.8 |
|  | Reverse | GCAGCCAGGACTTCCTCTTA | 63.4 |
| *PSMB2* | Forward | AGAGGGCAGTGGAACTCCTT | 64.1 |
|  | Reverse | AGGTTGGCAGATTCAGGATG | 64.0 |
| *OARD1* | Forward | GGCTTCGCACAAGCCAACTTATG | 70.4 |
|  | Reverse | GCAGACGATCAAGACCACATCC | 68.0 |

Table S9: Comparison of characteristics of WT, Mybpc3^+/insG^ and Mybpc3^insG/insG^ mice to show equality of groups for general parameters and the consistency of the model. VW/BW, ventricle weight over body weight; VW/TL, ventricle weight over tibia length; LVAW_d_, diastolic left ventricular anterior wall thickness; LVPW_d_, diastolic left ventricular posterior wall thickness; IVRT, isovolumetric relaxation time.

Table S8: Characteristics of mice used for embedding of whole mouse hearts. See separate excel sheet.

|  | **WT (*N*=6)** | ***Mybpc3^+/insG^ (N=6)*** | ***Mybpc3^insG/insG^ (N=6)*** | ***P* value** |
| --- | --- | --- | --- | --- |
|  | **General** | | | |
| Age (weeks) | 26.2 ± 1.0 | 25.8 ± 2.1 | 25.2 ± 0.41 | 0.20 |
| Sex, male | 66.7% (*N*=4) | 50% (*N*=3) | 50% (*N*=3) | 0.80 |
| Body weight (g) | 33.1 ± 7.5 | 30.2 ± 6.4 | 32.5 ± 7.9 | 0.78 |
|  | **Model validation** | | | |
| VW/BW (mg/g) | 4.0 ± 0.68 | 3.9 ± 0.42 | 7.4 ± 0.85 | 0.0005 |
| VW/TL (mg/mm) | 6.9 ± 0.71 | 6.2 ± 0.87 | 12.8 ± 2.6 | <0.0001 |
| LVAW_d_ (mm) | 1.0 ± 0.07 | 1.0 ± 0.07 | 1.4 ± 0.17 | <0.0001 |
| LVPW_d_ (mm) | 0.9 ± 0.18 | 1.0 ± 0.15 | 1.2 ± 0.28 | 0.075 |
| Fractional shortening (%) | 39.0 ± 3.2 | 41.3 ± 7.9 | 7.1 ± 1.0 | <0.0001 |
| IVRT (ms) | 8.5 ± 2.5 | 8.2 ± 3.6 | 32.5 ± 6.4 | <0.0001 |

Table S10: Characteristics of mice used for cardiomyocyte isolation. See separate excel sheet.

Table S11: Overview antibodies used for immunofluorescent staining.

| **Target** | **Isotype/host** | **Company/product number** | **Dilution** |  | **Secondary antibody** | **Dilution** |
| --- | --- | --- | --- | --- | --- | --- |
| PCM-1 | Rabbit | Invitrogen PA5-76769 | 1:100 |  | 555 | 1:250 |
| Lamin A/C | Mouse | Santa Cruz sc-376248 | 1:200 |  | 647 | 1:250 |
| YAP1 | Rabbit | Invitrogen PA1-46189 | 1:500 |  | 488 | 1:250 |

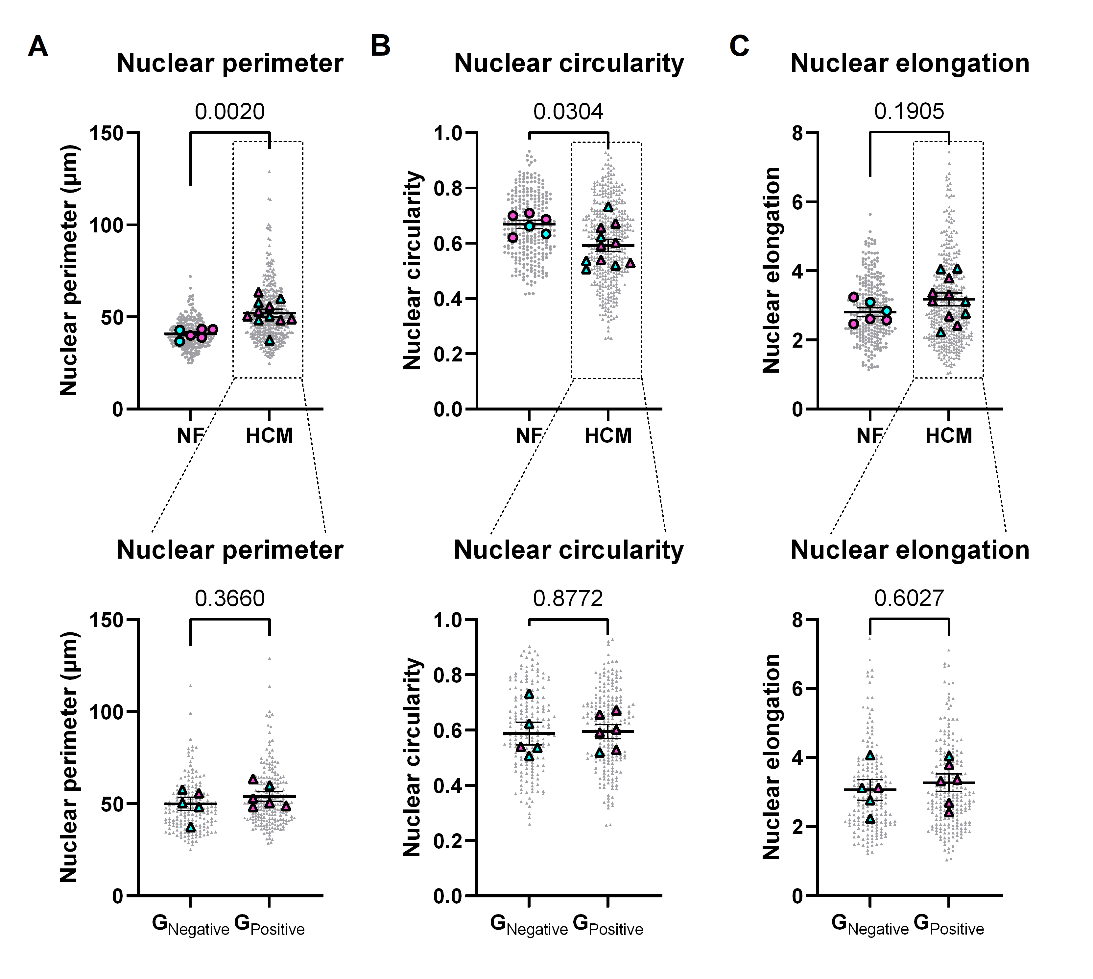

**Figure S1: Nuclear morphology in human patients with hypertrophic cardiomyopathy.** Quantification of the nuclear perimeter (A), nuclear circularity (B) and nuclear elongation (C) of NF donors and HCM patients (top panel) and G_Negative_ and G_Positive_ (bottom panel). Data are expressed as mean ± standard error of the mean. Every grey symbol represents the value of a single nucleus, n, and every colored symbol represents the average value per non-failing donor or HCM patient, N. Female are shown in magenta, male are shown in cyan, NF are shown as circles and HCM are shown as triangles. (A-C) NF: N=6 n=281; HCM: N=11 n=410; G_Negative_: N=5 n=190; G_Positive_: N=6 n=220. Statistical tests: unpaired t-test (A-C).

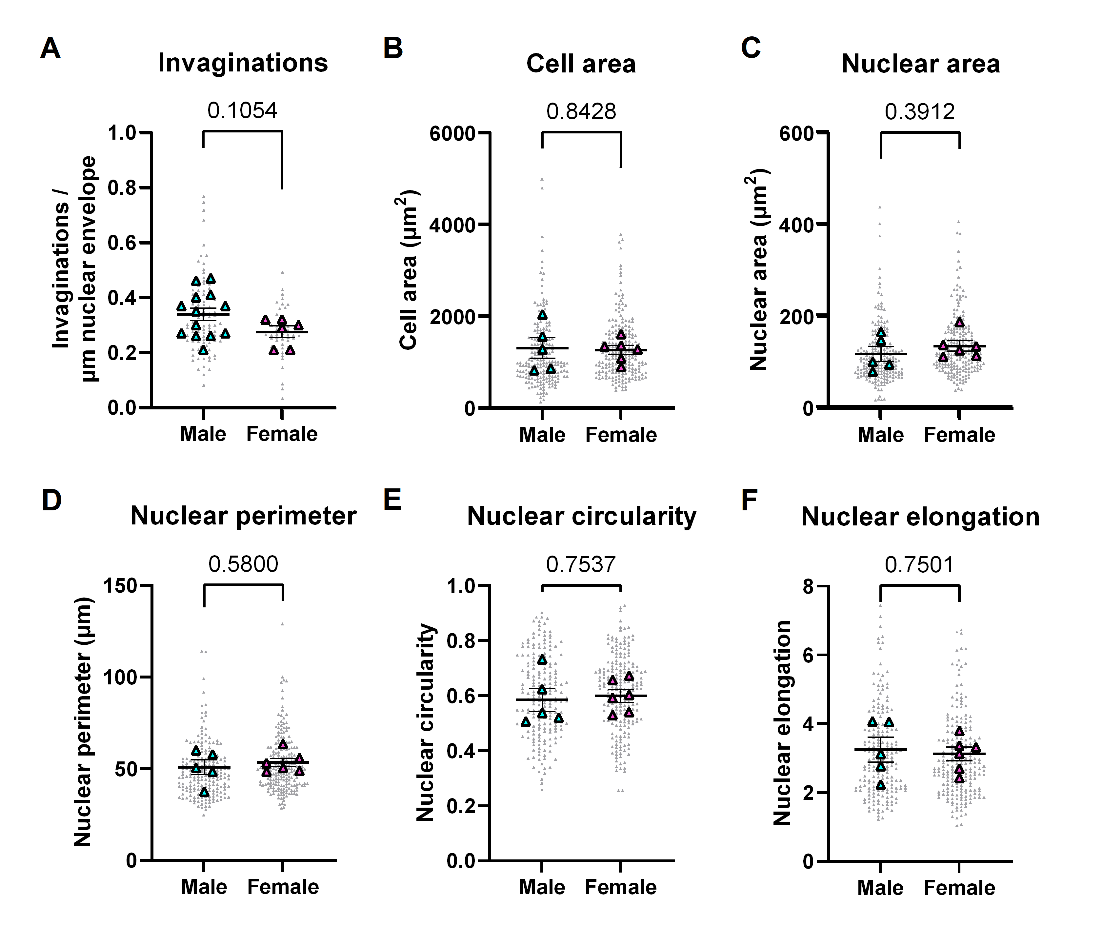

**Figure S2: Cell and nuclear morphology in male and female patients with hypertrophic cardiomyopathy.** Quantification of nuclear invagination (A), cell area (B), nuclear area (C), nuclear perimeter (D), nuclear circularity (E) and nuclear elongation (F) of male and female HCM patients. Data are expressed as mean ± standard error of the mean. Every grey symbol represents the value of a single nucleus, n, and every colored symbol represents the average value per non-failing donor or HCM patient, N. Female are shown in magenta, male are shown in cyan. (A) Male: N=13 n=101 ; Female: N=6 n=46, (B-F) Male: N=5 n=198 ; Female: N=6 n=212. Statistical tests: unpaired t-test (A-F).

Table S12: Uncorrected p-values of the Pearson R correlation of nuclear invaginations with several clinical parameter in Figure 1F.

|  | Invaginations | Age (at operation) | IVS | IVS/BSA | Lad/BSA | lvpw/BSA |
| --- | --- | --- | --- | --- | --- | --- |
| Invaginations |  | 0.1623 | 0.0083 | 0.0577 | 0.0872 | 0.2970 |
| Age (at operation) | 0.1623 |  | 0.0022 | 0.0061 | 0.0575 | 0.6837 |
| IVS | 0.0083 | 0.0022 |  | <0.0001 | 0.8551 | 0.0382 |
| IVS/BSA | 0.0577 | 0.0061 | <0.0001 |  | 0.7563 | 0.0025 |
| Lad/BSA | 0.0872 | 0.0575 | 0.8551 | 0.7563 |  | 0.3176 |
| lvpw/BSA | 0.2970 | 0.6837 | 0.0382 | 0.0025 | 0.3176 |  |

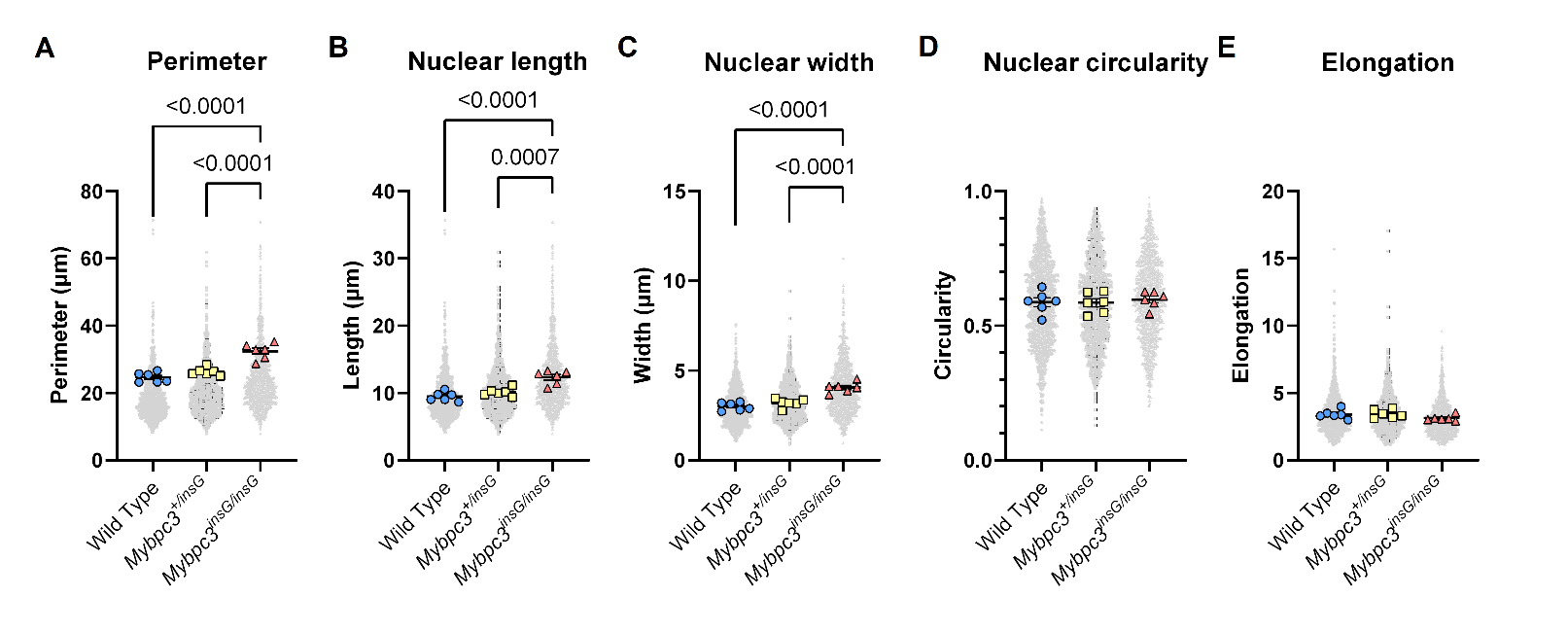

**Figure S3:** **2D nuclear morphology of WT, Mybpc3^+/insG^** **and Mybpc3^insG/insG^** **mice.** Quantification of the nuclear perimeter (A), nuclear length (B), nuclear width (C), nuclear circularity (D) and nuclear elongation (E). Nuclear elongation is calculated as the nuclear length divided by the nuclear width (E). Data are expressed as mean ± standard error of the mean. Every grey symbol represents the value of a single nucleus, n, and every colored symbol represents the average value per single mouse, N. (A-E) Wild Type: N=6 n=1371; Mybpc3^+/insG^: N=6 n=1371; Mybpc3^insG/insG^: N=6 n=1064. Statistical tests: Ordinary one-way ANOVA (A-E).

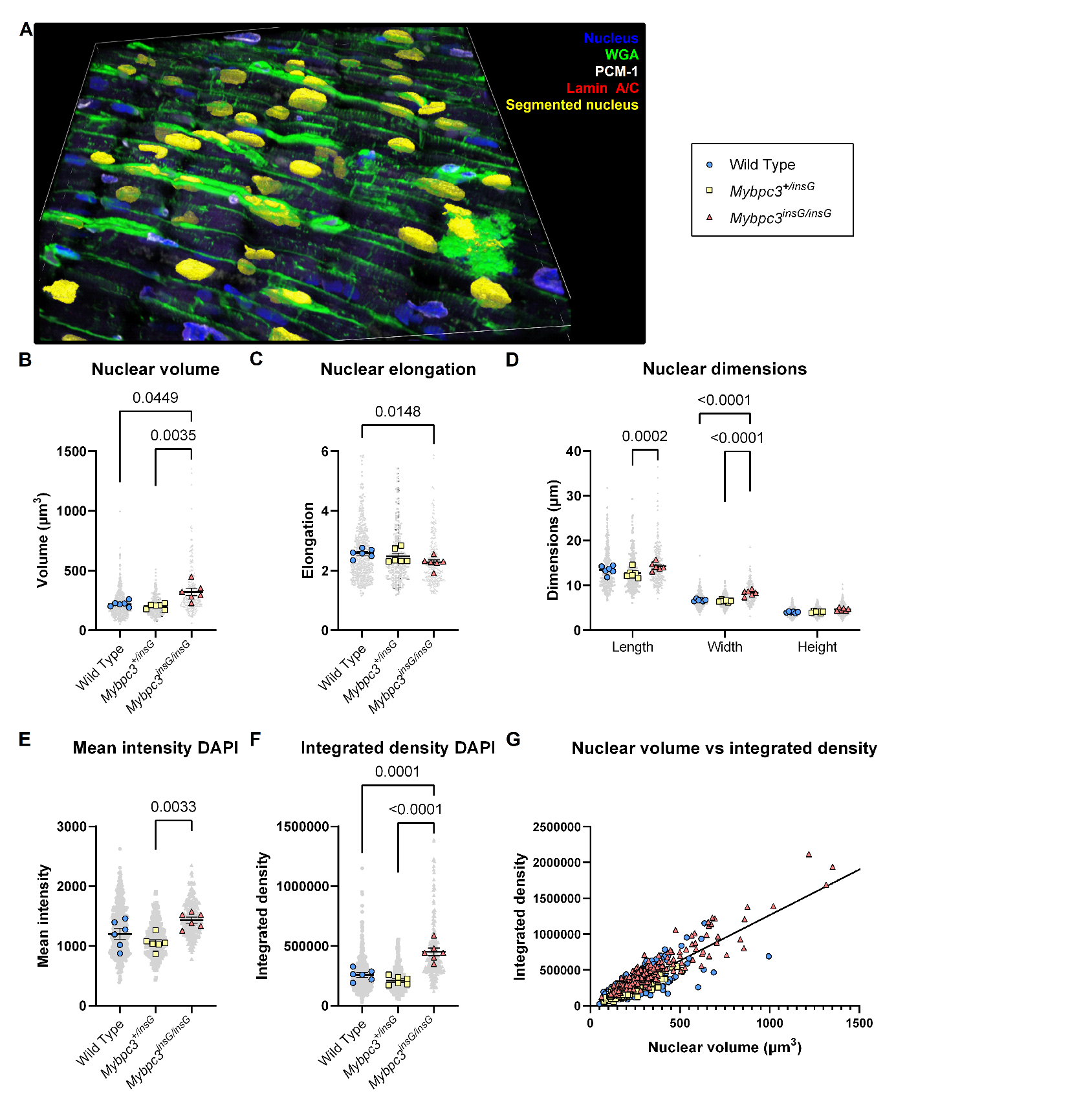

**Figure S4: 3D nuclear morphology of the WT, Mybpc3^+/insG^ and Mybpc3^insG/insG^ mouse model.** Nuclear segmentation of FFPE mouse model heart tissue stained with DAPI (blue), WGA (green), PCM-1 (white) and Lamin A/C (red) (A). Quantification of nuclear volume (B), nuclear elongation (C), nuclear dimensions (D), mean intensity of DAPI (E) and integrated density of DAPI (F) in WT, Mybpc3^+/insG^ and Mybpc3^insG/insG^ cardiomyocytes. Nuclear elongation is calculated as the nuclear length divided by the nuclear width (C). Simple linear regression of nuclear volume vs integrated density in WT, Mybpc3^+/insG^ and Mybpc3^insG/insG^ cardiomyocytes (p=<0.0001) (G). Data are expressed as mean ± standard error of the mean. Every grey symbol represents the value of a single nucleus, n, and every colored symbol represents the average value per single mouse, N. (A-G) WT: N=6 n=505; Mybpc3^+/insG^: N=6 n=259; Mybpc3^insG/insG^: N=6 n=238. Statistical tests: Kruskal-Wallis test (B,C), 2way ANOVA (D) and ordinary one-way ANOVA (E, F).

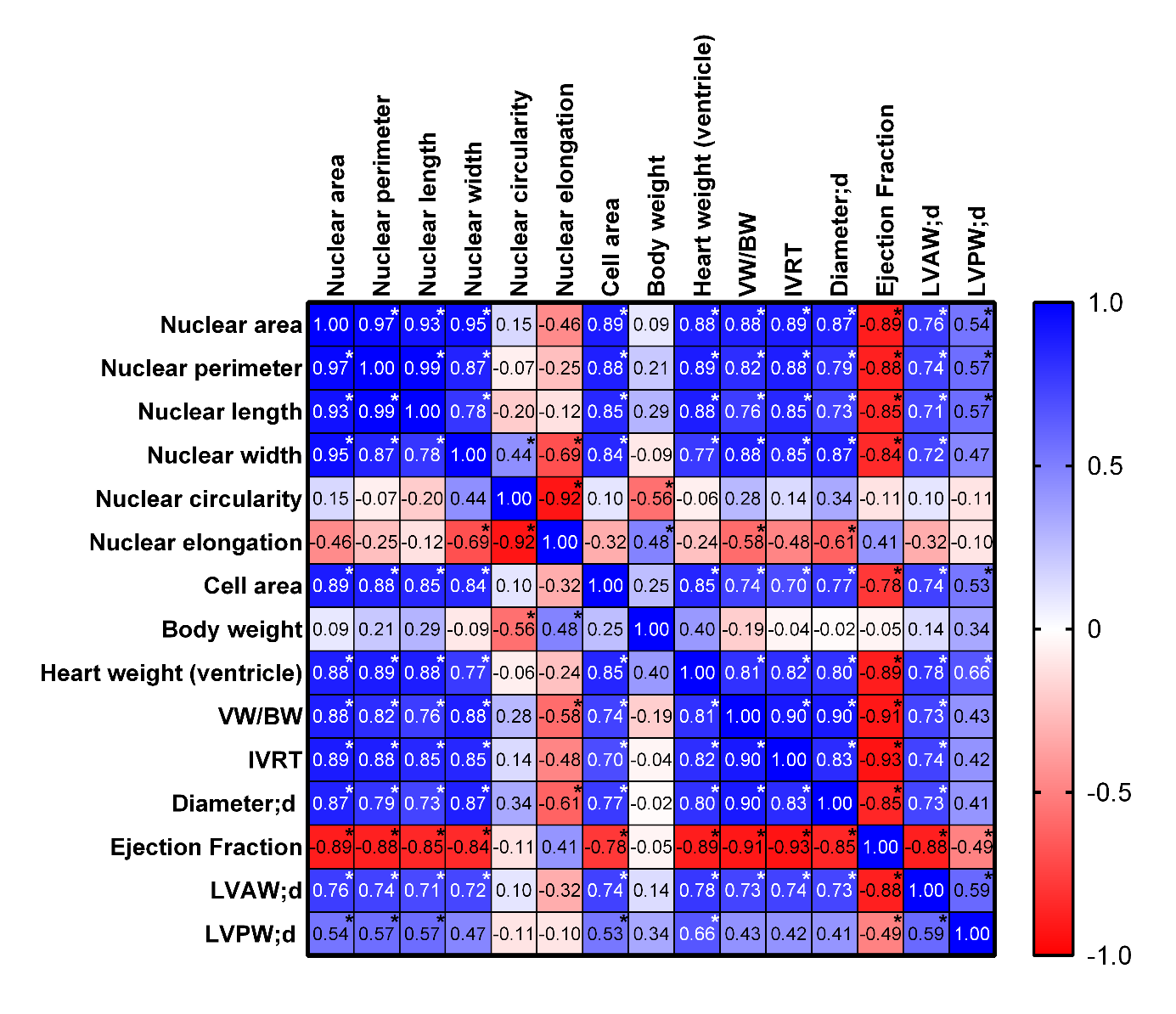

**Figure S5: Correlations of** **2D nuclear parameters with clinical parameters of WT, Mybpc3^+/insG^ and Mybpc3^insG/insG^ mice.** Wild Type: N=6; Mybpc3^+/insG^: N=6; Mybpc3^insG/insG^: N=6. (Pearson R correlation, *=p<0.05).

Table S13: Uncorrected p-values of the Pearson R correlation of 2D nuclear parameters with clinical parameters of WT, Mybpc3^+/insG^ and Mybpc3^insG/insG^ mice in Figure S5.

|  | Nuclear area | Nuclear perimeter | Nuclear length | Nuclear width | Nuclear circularity | Nuclear elongation | Cell area | Body weight | Heart weight (ventricle) | VW/BW | IVRT | Diameter;d | Ejection Fraction | LVAW;d |
| --- | --- | --- | --- | --- | --- | --- | --- | --- | --- | --- | --- | --- | --- | --- |
| Nuclear area |  | <0.0001 | <0.0001 | <0.0001 | 0.5479 | 0.0569 | <0.0001 | 0.7253 | <0.0001 | <0.0001 | <0.0001 | <0.0001 | <0.0001 | 0.0004 |
| Nuclear perimeter | <0.0001 |  | <0.0001 | <0.0001 | 0.7959 | 0.3099 | <0.0001 | 0.3985 | <0.0001 | <0.0001 | <0.0001 | 0.0001 | <0.0001 | 0.0007 |
| Nuclear length | <0.0001 | <0.0001 |  | 0.0001 | 0.4202 | 0.6410 | <0.0001 | 0.2370 | <0.0001 | 0.0003 | <0.0001 | 0.0009 | <0.0001 | 0.0013 |
| Nuclear width | <0.0001 | <0.0001 | 0.0001 |  | 0.0699 | 0.0016 | <0.0001 | 0.7121 | 0.0002 | <0.0001 | <0.0001 | <0.0001 | <0.0001 | 0.0010 |
| Nuclear circularity | 0.5479 | 0.7959 | 0.4202 | 0.0699 |  | <0.0001 | 0.6966 | 0.0148 | 0.7995 | 0.2640 | 0.5849 | 0.1857 | 0.6731 | 0.6980 |
| Nuclear elongation | 0.0569 | 0.3099 | 0.6410 | 0.0016 | <0.0001 |  | 0.1912 | 0.0419 | 0.3288 | 0.0122 | 0.0509 | 0.0086 | 0.0991 | 0.2069 |
| Cell area | <0.0001 | <0.0001 | <0.0001 | <0.0001 | 0.6966 | 0.1912 |  | 0.3141 | <0.0001 | 0.0004 | 0.0017 | 0.0003 | 0.0002 | 0.0006 |
| Body weight | 0.7253 | 0.3985 | 0.2370 | 0.7121 | 0.0148 | 0.0419 | 0.3141 |  | 0.1015 | 0.4411 | 0.8762 | 0.9503 | 0.8583 | 0.6047 |
| Heart weight (ventricle) | <0.0001 | <0.0001 | <0.0001 | 0.0002 | 0.7995 | 0.3288 | <0.0001 | 0.1015 |  | 0.0001 | 0.0001 | 0.0001 | <0.0001 | 0.0002 |
| VW/BW | <0.0001 | <0.0001 | 0.0003 | <0.0001 | 0.2640 | 0.0122 | 0.0004 | 0.4411 | 0.0001 |  | <0.0001 | <0.0001 | <0.0001 | 0.0008 |
| IVRT | <0.0001 | <0.0001 | <0.0001 | <0.0001 | 0.5849 | 0.0509 | 0.0017 | 0.8762 | 0.0001 | <0.0001 |  | <0.0001 | <0.0001 | 0.0007 |
| Diameter;d | <0.0001 | 0.0001 | 0.0009 | <0.0001 | 0.1857 | 0.0086 | 0.0003 | 0.9503 | 0.0001 | <0.0001 | <0.0001 |  | <0.0001 | 0.0009 |
| Ejection Fraction | <0.0001 | <0.0001 | <0.0001 | <0.0001 | 0.6731 | 0.0991 | 0.0002 | 0.8583 | <0.0001 | <0.0001 | <0.0001 | <0.0001 |  | <0.0001 |
| LVAW;d | 0.0004 | 0.0007 | 0.0013 | 0.0010 | 0.6980 | 0.2069 | 0.0006 | 0.6047 | 0.0002 | 0.0008 | 0.0007 | 0.0009 | <0.0001 |  |

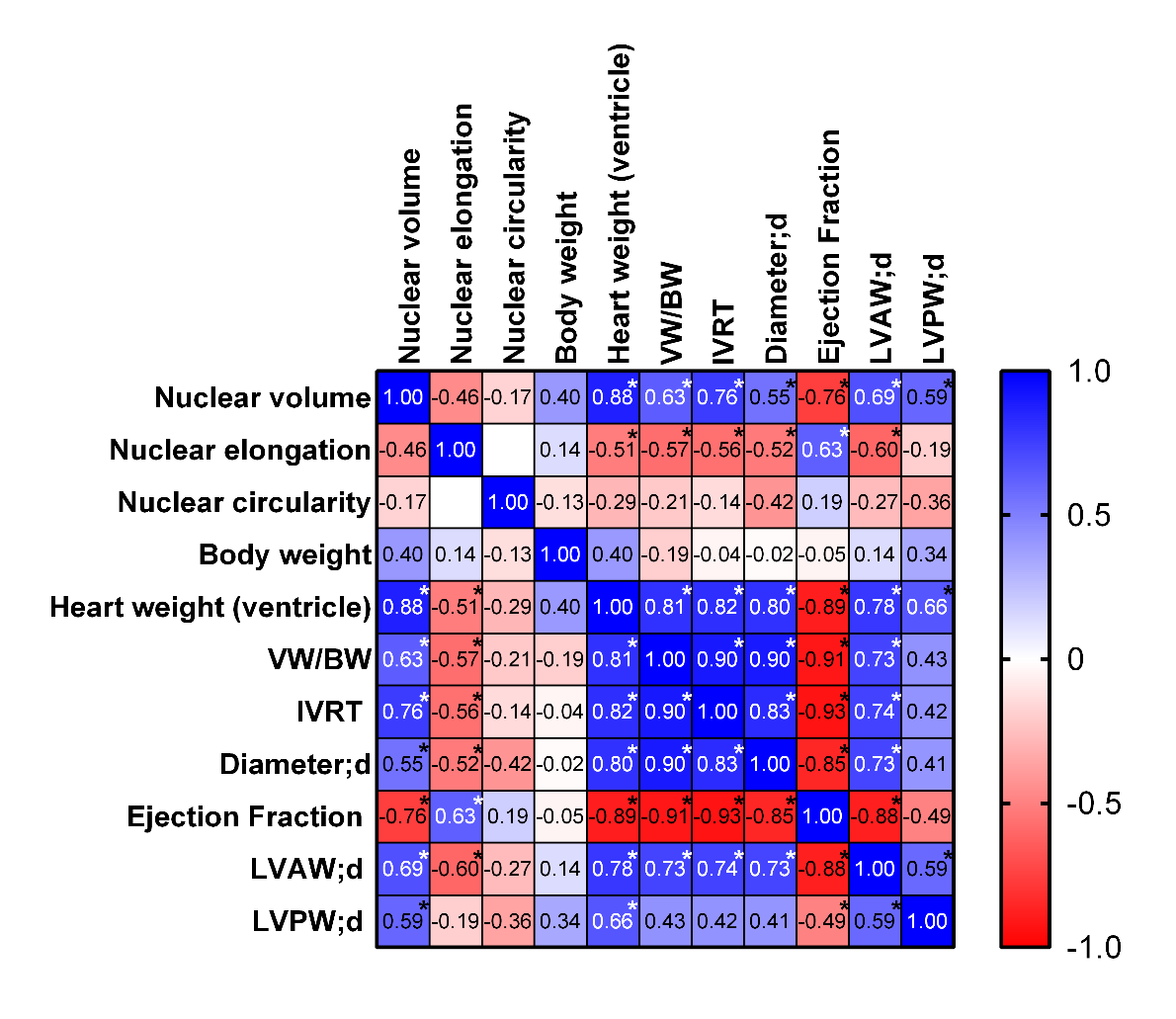

**Figure S6: Correlations of 3D nuclear parameters with clinical parameters of WT, Mybpc3^+/insG^ and Mybpc3^insG/insG^ mice.** Wild Type: N=6; Mybpc3^+/insG^: N=6; Mybpc3^insG/insG^: N=6. (Pearson R correlation, *=p<0.05).

Table S14: Uncorrected p-values of the Pearson R correlation of 3D nuclear parameters with clinical parameters of WT, Mybpc3^+/insG^ and Mybpc3^insG/insG^ mice in Figure S6.

|  | Nuclear volume | Nuclear elongation | Nuclear circularity | Body weight | Heart weight (ventricle) | VW/BW | IVRT | Diameter;d | Ejection Fraction | LVAW;d | LVPW;d |
| --- | --- | --- | --- | --- | --- | --- | --- | --- | --- | --- | --- |
| Nuclear volume |  | 0.0578 | 0.4930 | 0.0997 | <0.0001 | 0.0048 | 0.0004 | 0.0224 | 0.0004 | 0.0024 | 0.0119 |
| Nuclear elongation | 0.0578 |  | 0.9945 | 0.5882 | 0.0303 | 0.0131 | 0.0185 | 0.0310 | 0.0072 | 0.0113 | 0.4643 |
| Nuclear circularity | 0.4930 | 0.9945 |  | 0.6073 | 0.2461 | 0.3919 | 0.5798 | 0.0903 | 0.4621 | 0.2976 | 0.1514 |
| Body weight | 0.0997 | 0.5882 | 0.6073 |  | 0.1015 | 0.4411 | 0.8762 | 0.9503 | 0.8583 | 0.6047 | 0.1804 |
| Heart weight (ventricle) | <0.0001 | 0.0303 | 0.2461 | 0.1015 |  | 0.0001 | 0.0001 | 0.0001 | <0.0001 | 0.0002 | 0.0039 |
| VW/BW | 0.0048 | 0.0131 | 0.3919 | 0.4411 | 0.0001 |  | <0.0001 | <0.0001 | <0.0001 | 0.0008 | 0.0885 |
| IVRT | 0.0004 | 0.0185 | 0.5798 | 0.8762 | 0.0001 | <0.0001 |  | <0.0001 | <0.0001 | 0.0007 | 0.0911 |
| Diameter;d | 0.0224 | 0.0310 | 0.0903 | 0.9503 | 0.0001 | <0.0001 | <0.0001 |  | <0.0001 | 0.0009 | 0.0985 |
| Ejection Fraction | 0.0004 | 0.0072 | 0.4621 | 0.8583 | <0.0001 | <0.0001 | <0.0001 | <0.0001 |  | <0.0001 | 0.0446 |
| LVAW;d | 0.0024 | 0.0113 | 0.2976 | 0.6047 | 0.0002 | 0.0008 | 0.0007 | 0.0009 | <0.0001 |  | 0.0130 |
| LVPW;d | 0.0119 | 0.4643 | 0.1514 | 0.1804 | 0.0039 | 0.0885 | 0.0911 | 0.0985 | 0.0446 | 0.0130 |  |

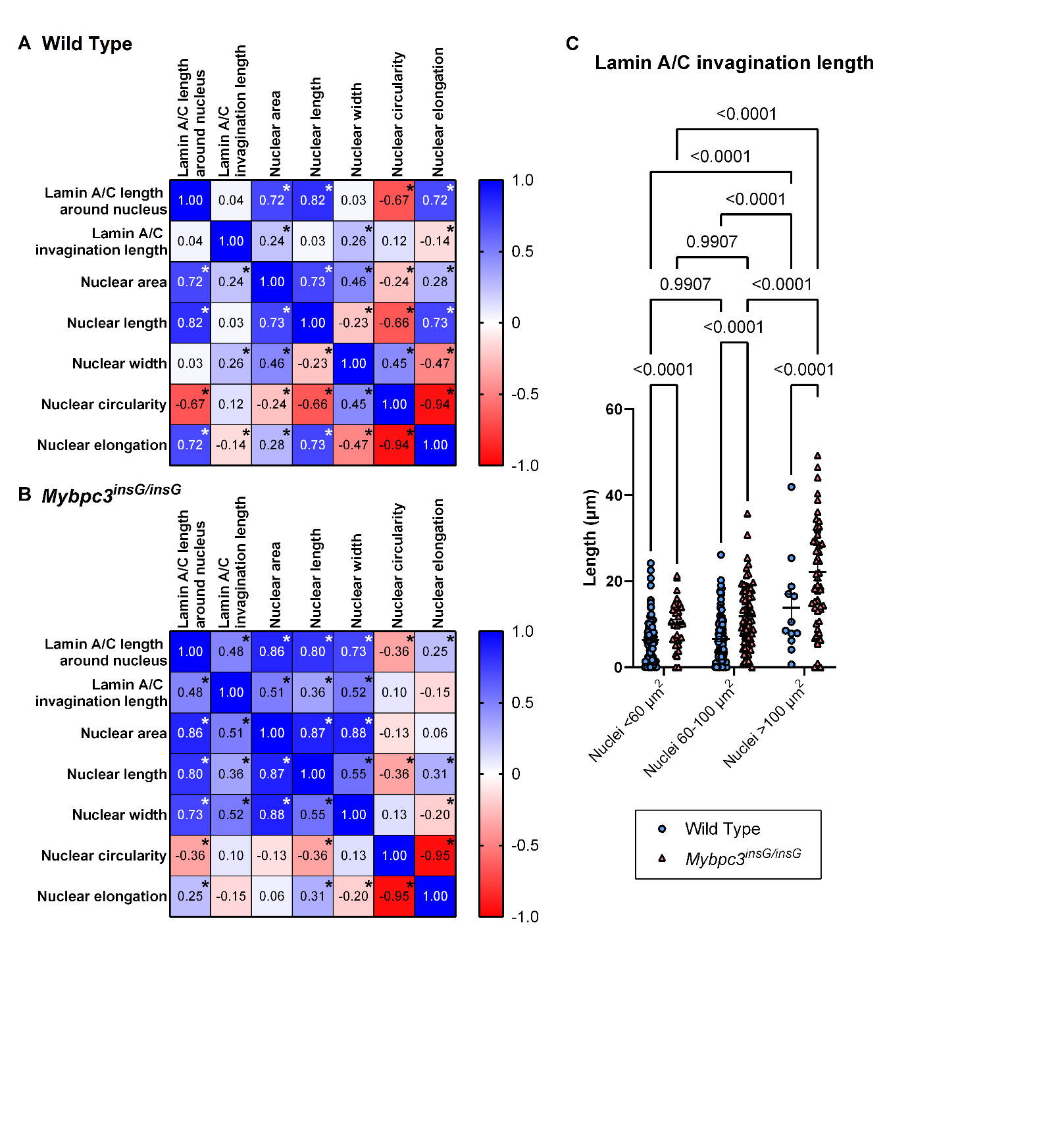

**Figure S7: The relationship between nuclear size and nuclear invagination.** Pearson R correlation between nuclear morphology and invagination parameters in WT (A) and Mybpc3^insG/insG^ (B). Quantification of total nuclear invagination length of WT and Mybpc3^insG/insG^ for 3 bins of different nuclear size: <60 µm^2^, 60-100 µm^2^ and >100 µm^2^ (C). Data are expressed as mean ± standard error of the mean. Every colored symbol represents the value for a single nucleus, n. (A) N=6 n=206, (B) N=6 n=153, (C) WT, <60 µm^2^: n=95; WT, 60-100 µm^2^: n=99; WT, >100 µm^2^: n=12; Mybpc3^insG/insG^ , <60 µm^2^: n=29, Mybpc3^insG/insG^, 60-100 µm^2^: n=68, Mybpc3^insG/insG^, >100 µm^2^: n=50. Statistical tests: Pearson R correlation (A, B, *=p<0.05) and 2-way ANOVA (C).

Table S15: Uncorrected p-values of the Pearson R correlation between nuclear morphology and invagination parameters in WT mice in Figure S7A.

|  | Lamin A/C length  around nucleus | Lamin A/C  invagination length | Nuclear area | Nuclear length | Nuclear width | Nuclear circularity | Nuclear elongation |
| --- | --- | --- | --- | --- | --- | --- | --- |
| Lamin A/C length  around nucleus |  | 0.5660 | <0.0001 | <0.0001 | 0.6894 | <0.0001 | <0.0001 |
| Lamin A/C  invagination length | 0.5660 |  | 0.0006 | 0.6607 | 0.0002 | 0.0765 | 0.0442 |
| Nuclear area | <0.0001 | 0.0006 |  | <0.0001 | <0.0001 | 0.0004 | <0.0001 |
| Nuclear length | <0.0001 | 0.6607 | <0.0001 |  | 0.0010 | <0.0001 | <0.0001 |
| Nuclear width | 0.6894 | 0.0002 | <0.0001 | 0.0010 |  | <0.0001 | <0.0001 |
| Nuclear circularity | <0.0001 | 0.0765 | 0.0004 | <0.0001 | <0.0001 |  | <0.0001 |
| Nuclear elongation | <0.0001 | 0.0442 | <0.0001 | <0.0001 | <0.0001 | <0.0001 |  |

Table S16: Uncorrected p-values of the Pearson R correlation between nuclear morphology and invagination parameters in Mybpc3^insG/insG^ mice in Figure S7B.

|  | Lamin A/C length  around nucleus | Lamin A/C  invagination length | Nuclear area | Nuclear length | Nuclear width | Nuclear circularity | Nuclear elongation |
| --- | --- | --- | --- | --- | --- | --- | --- |
| Lamin A/C length  around nucleus |  | <0.0001 | <0.0001 | <0.0001 | <0.0001 | <0.0001 | 0.0017 |
| Lamin A/C  invagination length | <0.0001 |  | <0.0001 | <0.0001 | <0.0001 | 0.2346 | 0.0591 |
| Nuclear area | <0.0001 | <0.0001 |  | <0.0001 | <0.0001 | 0.1235 | 0.4610 |
| Nuclear length | <0.0001 | <0.0001 | <0.0001 |  | <0.0001 | <0.0001 | 0.0001 |
| Nuclear width | <0.0001 | <0.0001 | <0.0001 | <0.0001 |  | 0.1114 | 0.0155 |
| Nuclear circularity | <0.0001 | 0.2346 | 0.1235 | <0.0001 | 0.1114 |  | <0.0001 |
| Nuclear elongation | 0.0017 | 0.0591 | 0.4610 | 0.0001 | 0.0155 | <0.0001 |  |

**Video S1: Representative video of nuclear deformation while the cardiomyocyte is contracting.**

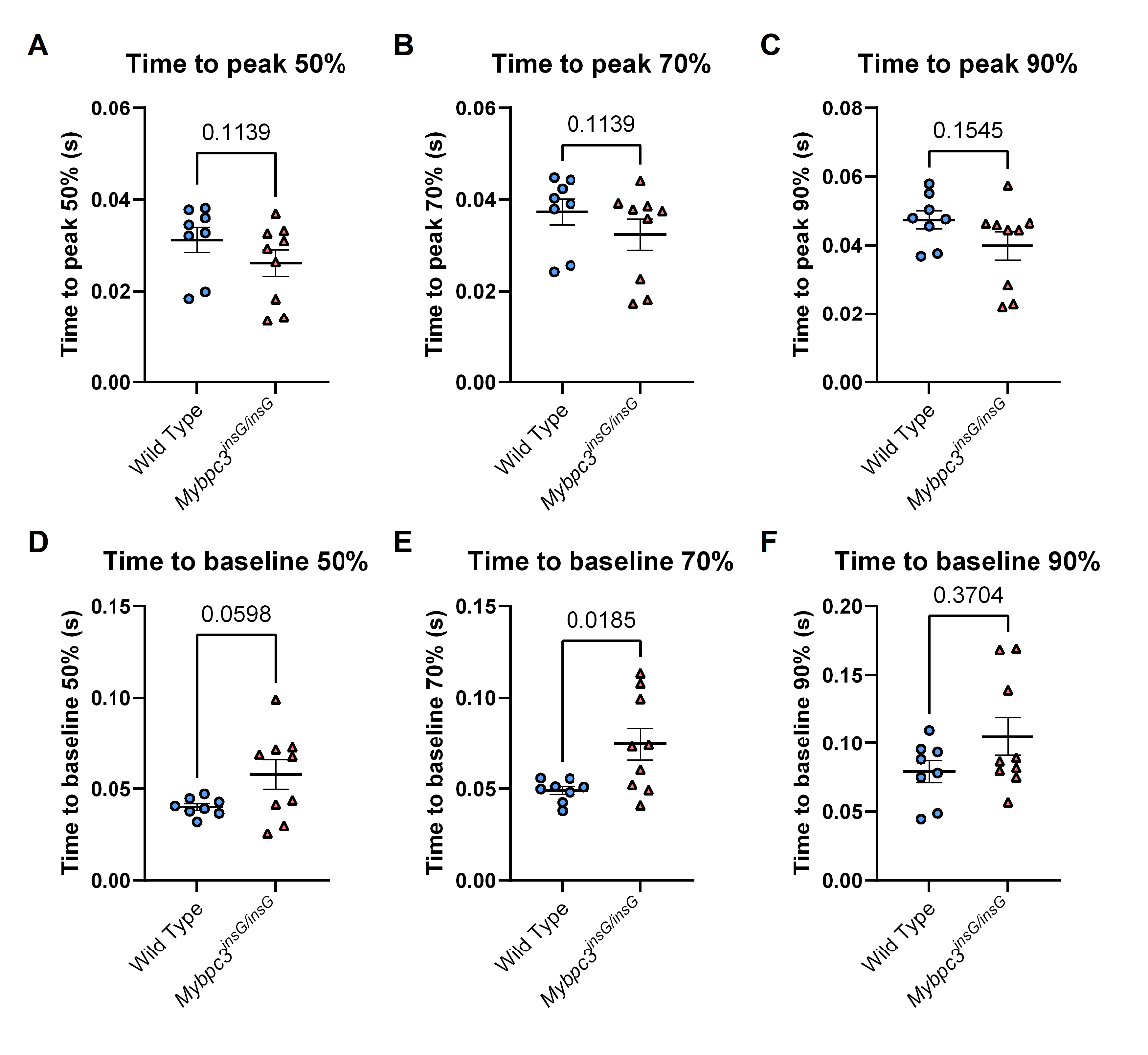

**Figure S8: Contraction and relaxation time of isolated cardiomyocytes of WT and Mybpc3^insG/insG^ mice.** Quantification of the time to peak at a sarcomere shortening of 50% (A), 70% (B) and 90% (C) of maximum sarcomere shortening. Quantification of the relaxation using time to baseline at a sarcomere shortening of 50% (D), 70% (E) and 90% (F) of maximum sarcomere shortening. Every colored symbol represents the average value per single mouse, N. The data are expressed as mean ± standard error of the mean. Wild Type: N=8 n=223; Mybpc3^insG/insG^: N=9 n=145, Statistical tests: Mann-Whitney test (A, B, F) unpaired t-test (C-E).

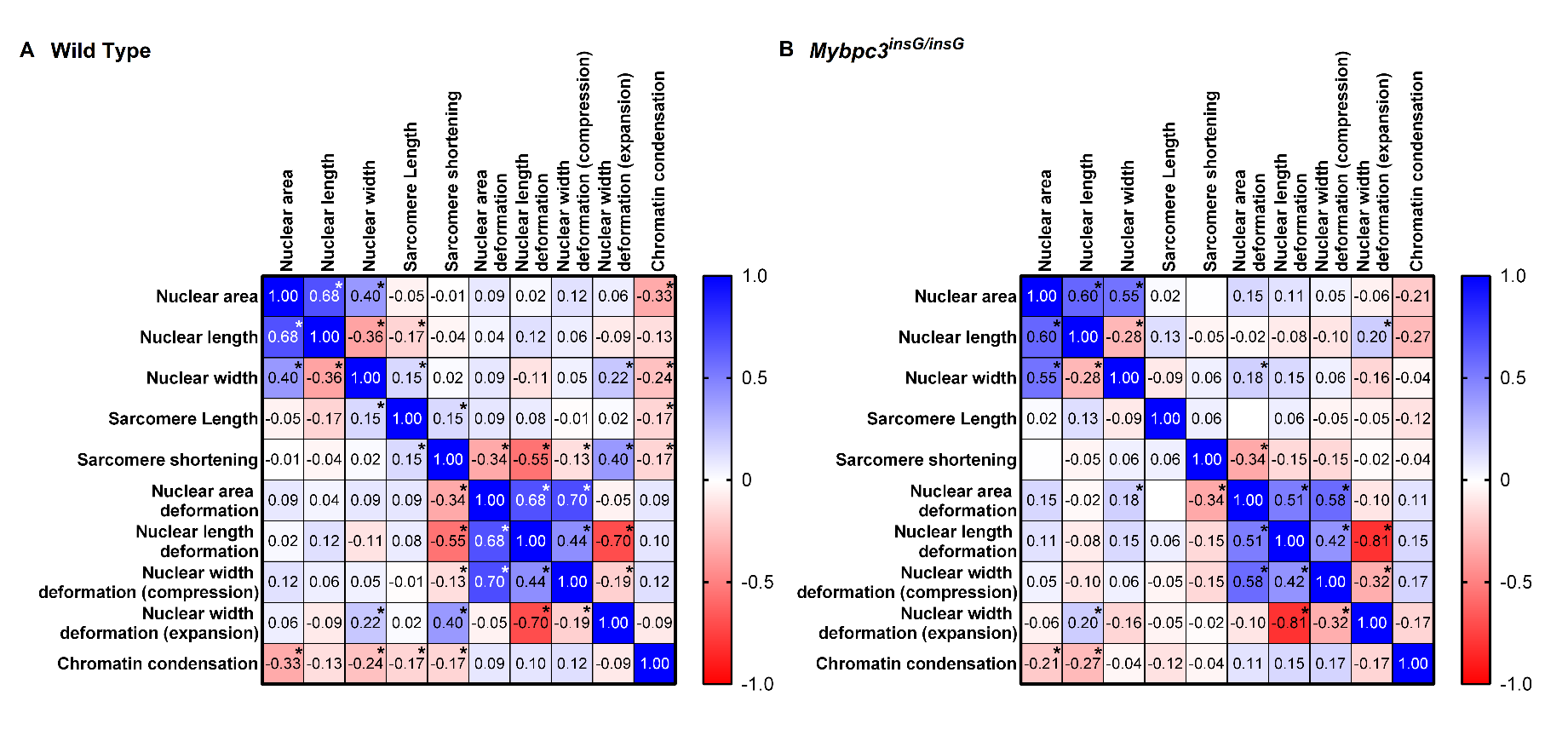

**Figure S9: Correlations of nuclear parameters with deformability.** Pearson R correlation of nuclear morphology and nuclear deformation parameters in WT (A) and in Mybpc3^insG/insG^ cardiomyocytes (B). (A) WT: N=8 n=223; Mybpc3^insG/insG^: N=9 n=145. Statistical tests: Pearson R correlation (A, B, *=p<0.05).

Table S17: Uncorrected p-values of the Pearson R correlation of nuclear parameters with deformability in WT mice in Figure S9A.

|  | Nuclear length | Nuclear width | Sarcomere Length | Sarcomere shortening | Nuclear area  deformation | Nuclear length  deformation | Nuclear width  deformation (compression) | Nuclear width  deformation (expansion) | Chromatin condensation | Nuclear length |
| --- | --- | --- | --- | --- | --- | --- | --- | --- | --- | --- |
| Nuclear area |  | <0.0001 | <0.0001 | 0.4341 | 0.9173 | 0.1953 | 0.7120 | 0.0852 | 0.3406 | <0.0001 |
| Nuclear length | <0.0001 |  | <0.0001 | 0.0114 | 0.5313 | 0.5381 | 0.0786 | 0.3503 | 0.2001 | 0.0517 |
| Nuclear width | <0.0001 | <0.0001 |  | 0.0248 | 0.7993 | 0.1628 | 0.1016 | 0.4289 | 0.0011 | 0.0003 |
| Sarcomere Length | 0.4341 | 0.0114 | 0.0248 |  | 0.0270 | 0.1609 | 0.2453 | 0.8396 | 0.8107 | 0.0147 |
| Sarcomere shortening | 0.9173 | 0.5313 | 0.7993 | 0.0270 |  | <0.0001 | <0.0001 | 0.0482 | <0.0001 | 0.0137 |
| Nuclear area  deformation | 0.1953 | 0.5381 | 0.1628 | 0.1609 | <0.0001 |  | <0.0001 | <0.0001 | 0.4397 | 0.1843 |
| Nuclear length  deformation | 0.7120 | 0.0786 | 0.1016 | 0.2453 | <0.0001 | <0.0001 |  | <0.0001 | <0.0001 | 0.1384 |
| Nuclear width  deformation (compression) | 0.0852 | 0.3503 | 0.4289 | 0.8396 | 0.0482 | <0.0001 | <0.0001 |  | 0.0049 | 0.0730 |
| Nuclear width  deformation (expansion) | 0.3406 | 0.2001 | 0.0011 | 0.8107 | <0.0001 | 0.4397 | <0.0001 | 0.0049 |  | 0.1975 |
| Chromatin condensation | <0.0001 | 0.0517 | 0.0003 | 0.0147 | 0.0137 | 0.1843 | 0.1384 | 0.0730 | 0.1975 |  |

Table S18: Uncorrected p-values of the Pearson R correlation of nuclear parameters with deformability in Mybpc3^insG/insG^ mice in Figure S9B.

|  | Nuclear length | Nuclear width | Sarcomere Length | Sarcomere shortening | Nuclear area  deformation | Nuclear length  deformation | Nuclear width  deformation (compression) | Nuclear width  deformation (expansion) | Chromatin condensation | Nuclear length |
| --- | --- | --- | --- | --- | --- | --- | --- | --- | --- | --- |
| Nuclear area |  | <0.0001 | <0.0001 | 0.8218 | 0.9949 | 0.0662 | 0.1871 | 0.5459 | 0.4914 | 0.0141 |
| Nuclear length | <0.0001 |  | 0.0006 | 0.1243 | 0.5901 | 0.7755 | 0.3275 | 0.2158 | 0.0185 | 0.0014 |
| Nuclear width | <0.0001 | 0.0006 |  | 0.2965 | 0.4914 | 0.0269 | 0.0783 | 0.4848 | 0.0529 | 0.6728 |
| Sarcomere Length | 0.8218 | 0.1243 | 0.2965 |  | 0.4551 | 0.9687 | 0.4442 | 0.5881 | 0.5681 | 0.1771 |
| Sarcomere shortening | 0.9949 | 0.5901 | 0.4914 | 0.4551 |  | <0.0001 | 0.0707 | 0.0651 | 0.8253 | 0.6270 |
| Nuclear area  deformation | 0.0662 | 0.7755 | 0.0269 | 0.9687 | <0.0001 |  | <0.0001 | <0.0001 | 0.2113 | 0.2128 |
| Nuclear length  deformation | 0.1871 | 0.3275 | 0.0783 | 0.4442 | 0.0707 | <0.0001 |  | <0.0001 | <0.0001 | 0.0798 |
| Nuclear width  deformation (compression) | 0.5459 | 0.2158 | 0.4848 | 0.5881 | 0.0651 | <0.0001 | <0.0001 |  | 0.0001 | 0.0545 |
| Nuclear width  deformation (expansion) | 0.4914 | 0.0185 | 0.0529 | 0.5681 | 0.8253 | 0.2113 | <0.0001 | 0.0001 |  | 0.0537 |
| Chromatin condensation | 0.0141 | 0.0014 | 0.6728 | 0.1771 | 0.6270 | 0.2128 | 0.0798 | 0.0545 | 0.0537 |  |

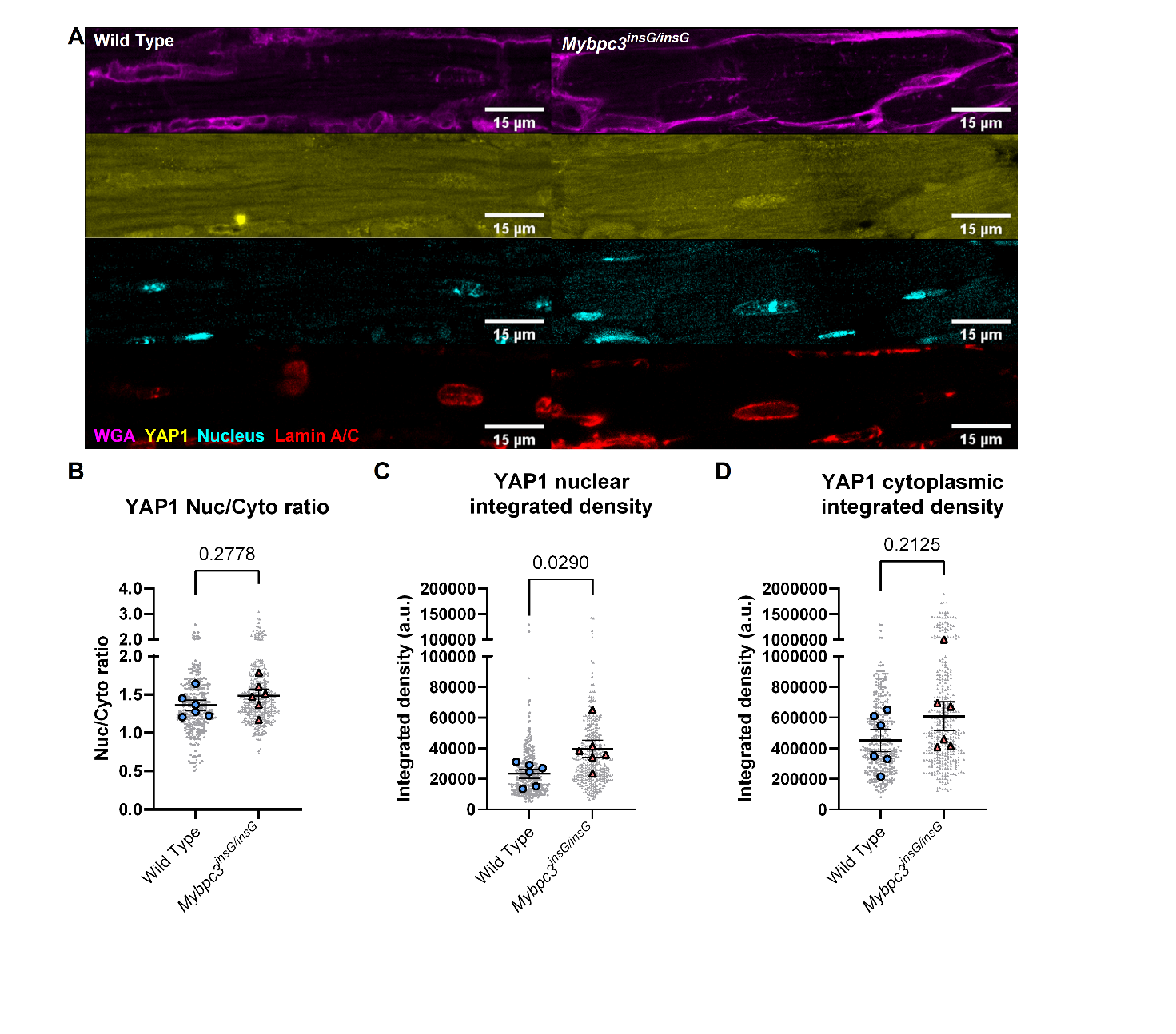

**Figure S10: YAP localization and protein levels of FFPE heart tissue of WT and Mybpc3^insG/insG^ mice.** Representative images of FFPE heart tissue of WT and Mybpc3^insG/insG^ mice stained with WGA (magenta), YAP1 (yellow), DAPI/nucleus (cyan) and lamin A/C (red) (A). Quantification of YAP 1 nuclear to cytoplasmic ratio in WT and Mybpc3^insG/insG^ cardiomyocytes. The ratio was calculated by dividing the mean intensity of YAP1 in the nucleus by the mean intensity in the cytoplasm (B). Quantification of the YAP integrated density of the nucleus (C) and the cytoplasm (D) in WT and Mybpc3^insG/insG^ cardiomyocytes. Data are expressed as mean ± standard error of the mean. Every grey symbol represents the value of a single nucleus, n, and every colored symbol represents the average value per single mouse, N. (A, B)) WT: N=6 n=330; Mybpc3^insG/insG^: N=6 n=361, (C) WT: N=6 n=271; Mybpc3^insG/insG^: N=6 n=302. Statistical tests: Unpaired t-test (A-C).

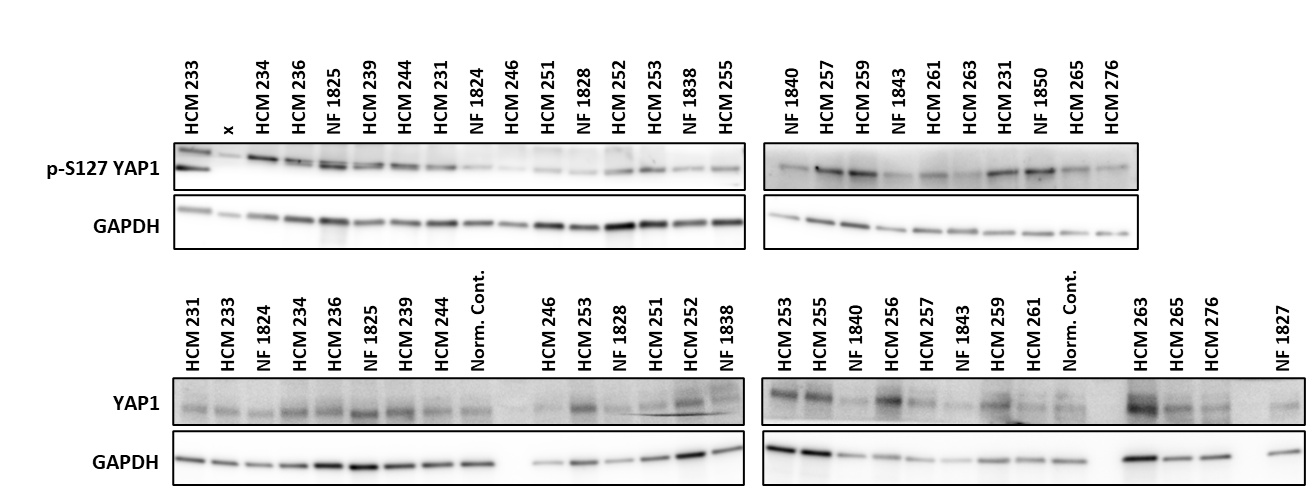

**Figure S11: Images of western blots p-S127 YAP1 and YAP1.**

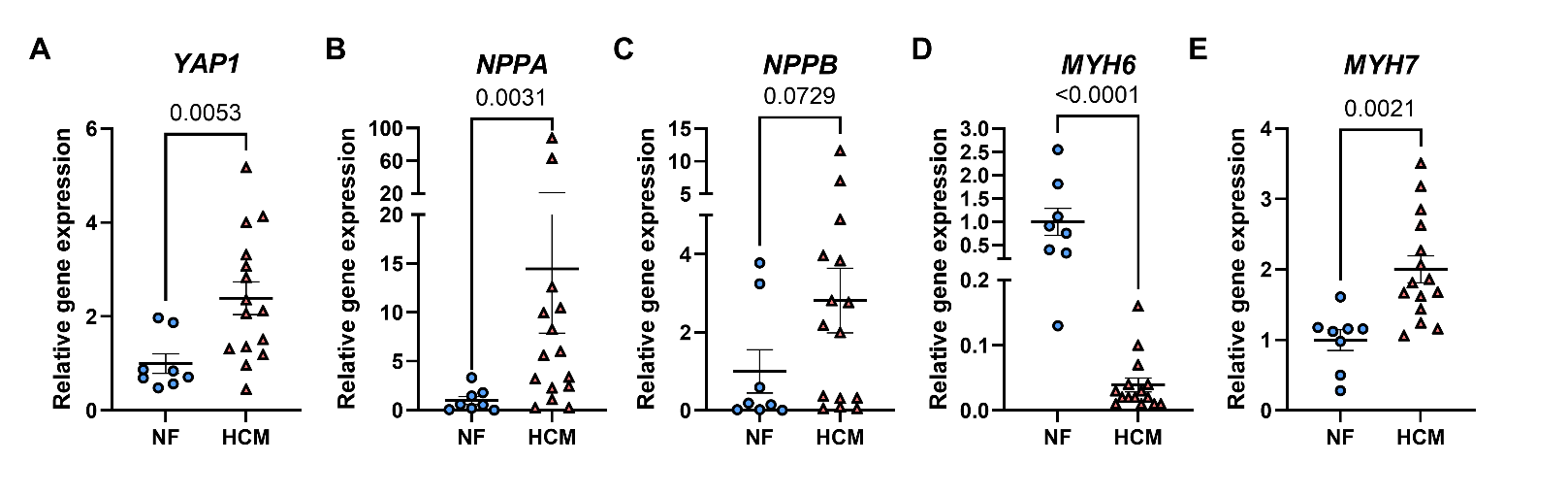

**Figure S12: Relative gene expression YAP1 (A) NPPA (B), NPPB (C), MYH6 (D) and MYH7 (E).** Quantification of gene expression determined by RT-qPCR of YAP1 (A) NPPA (B), NPPB (C), MYH6 (D) and MYH7 (E). Gene expression is normalized to the housekeeping genes PSMB2 and OARD1 (A-E). Data are expressed as mean ± standard error of the mean. Every colored symbol represents the average value per single donor or patient, N. (A-E) NF: N=8; HCM: N=15. Statistical tests: Mann-Whitney test (A- D), Unpaired t-test (E).

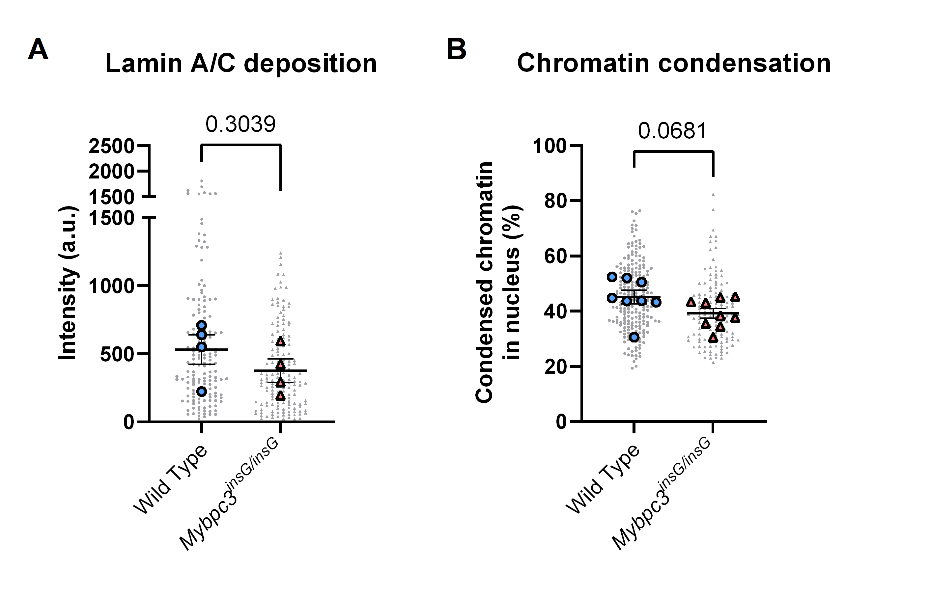

**Figure S13: Lamin A/C deposition and chromatin condensation of WT and Mybpc3^insG/insG^ cardiomyocytes.** Quantification of Lamin A/C deposition (A) and chromatin condensation (B) in WT and Mybpc3^insG/insG^ cardiomyocytes. Lamin A/C deposition is determined as the mean intensity of the Lamin A/C signal (A). Chromatin condensation is determined as the percentage of heterochromatin in the nucleus (B). Data are expressed as mean ± standard error of the mean. Every grey symbol represents the value of a single nucleus, n, and every colored symbol represents the average value per single mouse, N. (A) WT: N=4 n=156; Mybpc3^insG/insG^: N=4 n=154 (B) WT: N=8 n=218; Mybpc3^insG/insG^: N=9 n=137. Statistical tests: Unpaired t-test (A,B).

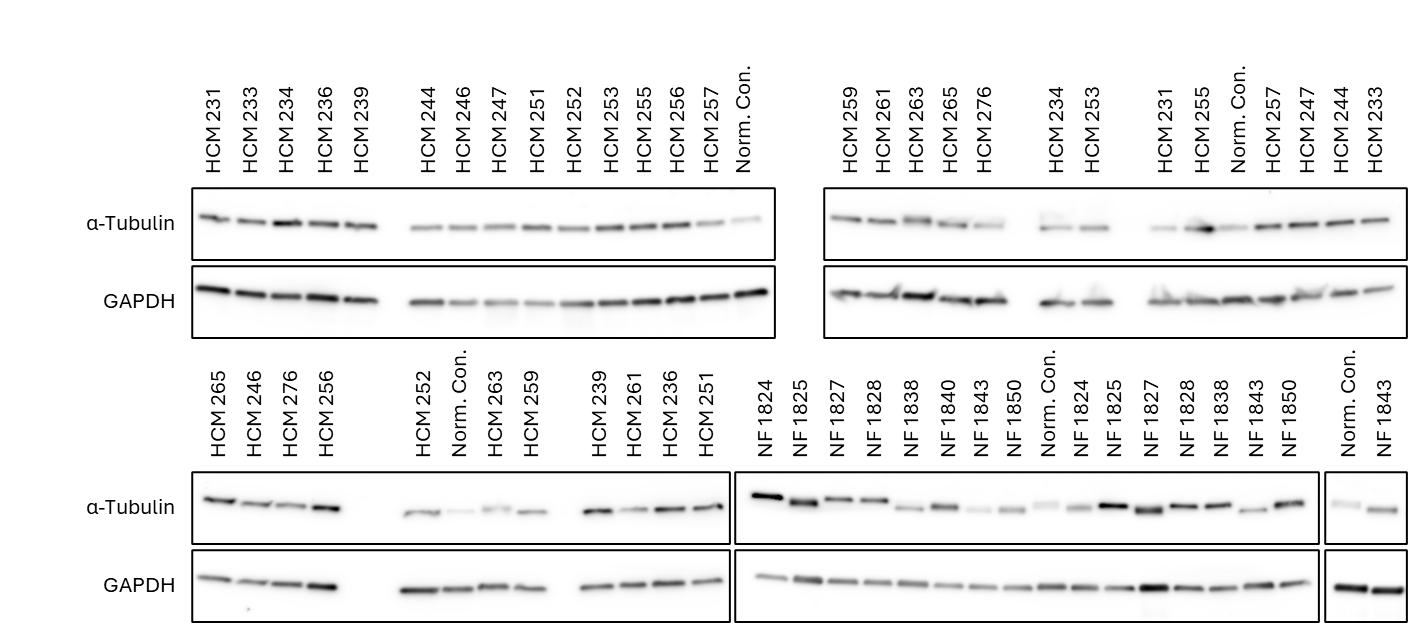

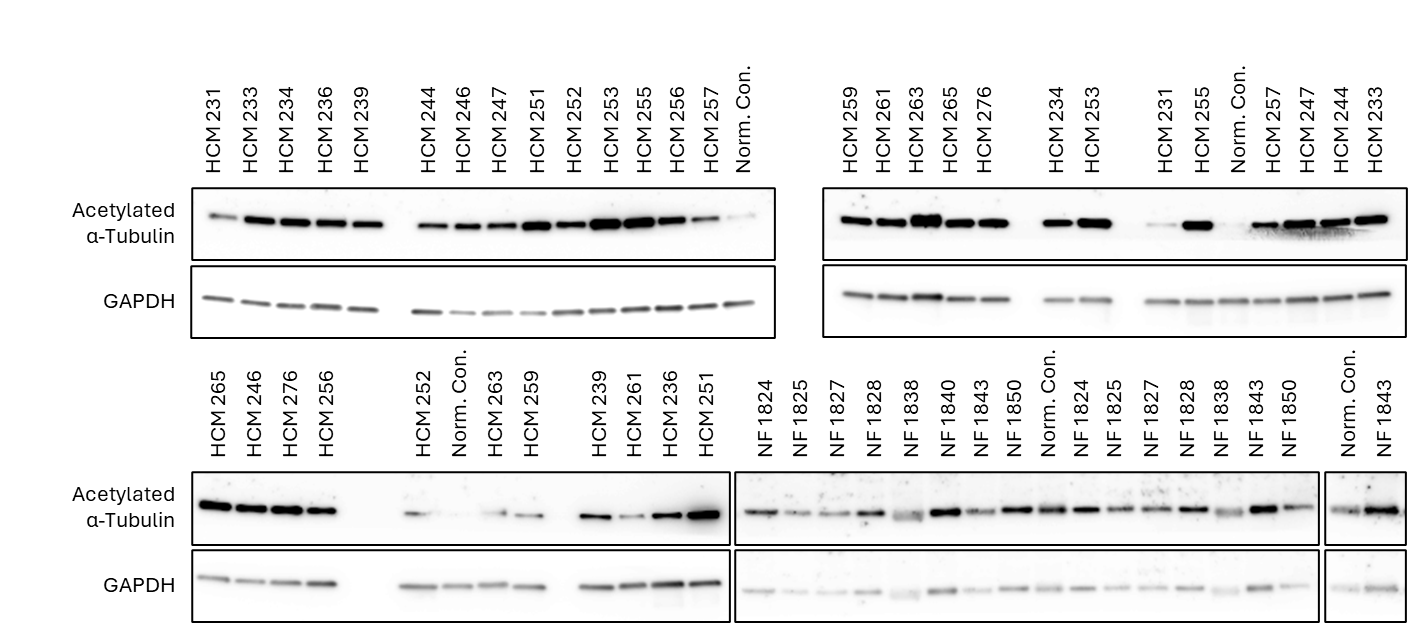

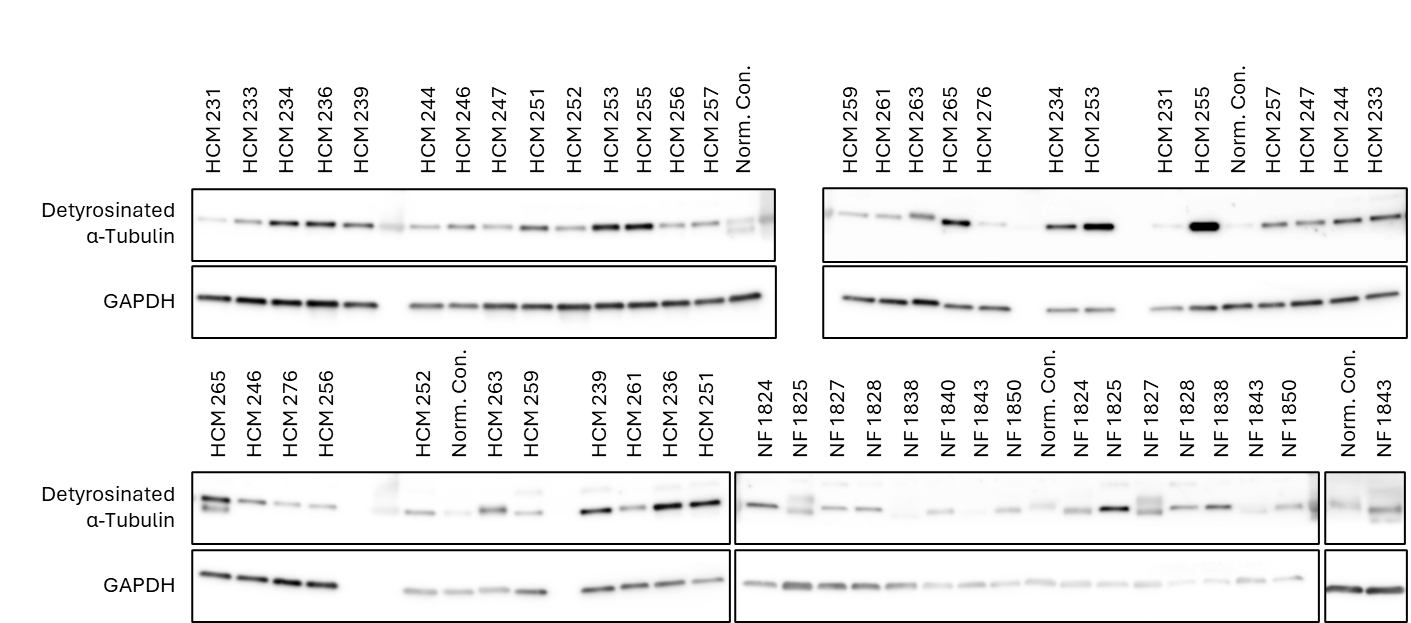

**Figure S16: Images of western blots of acetylated α-tubulin.**

**Figure S15: Images of western blots of detyrosinated α-tubulin.**

**Figure S14: Images of western blots of α-tubulin.**

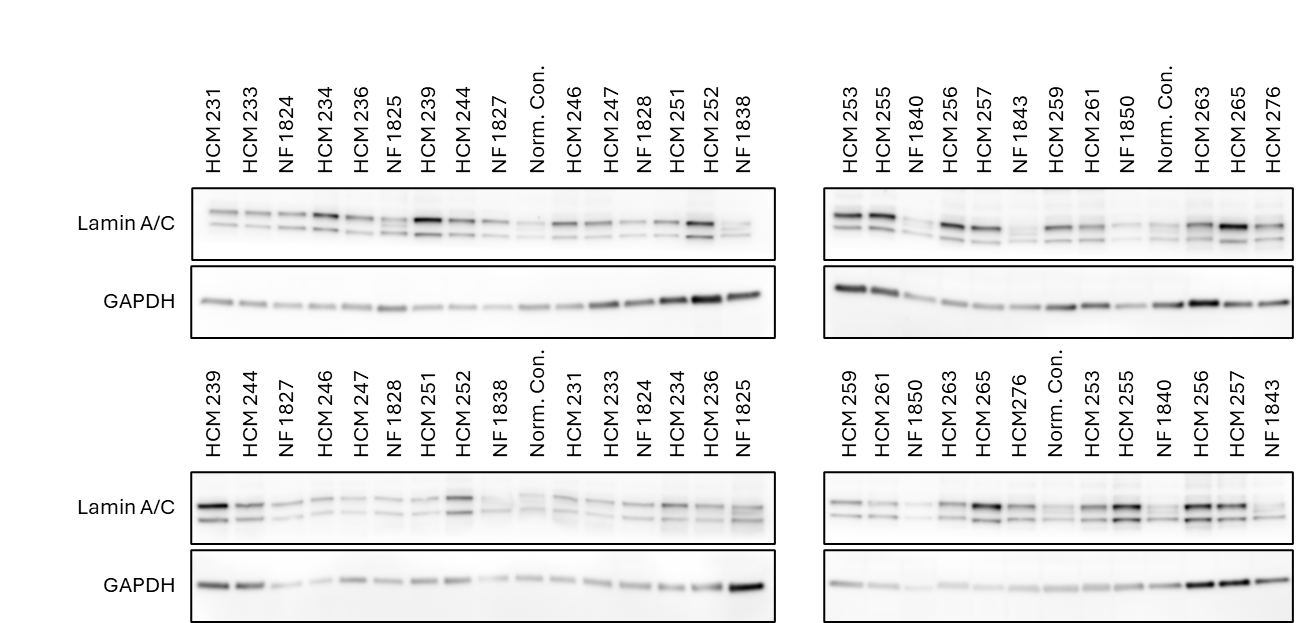

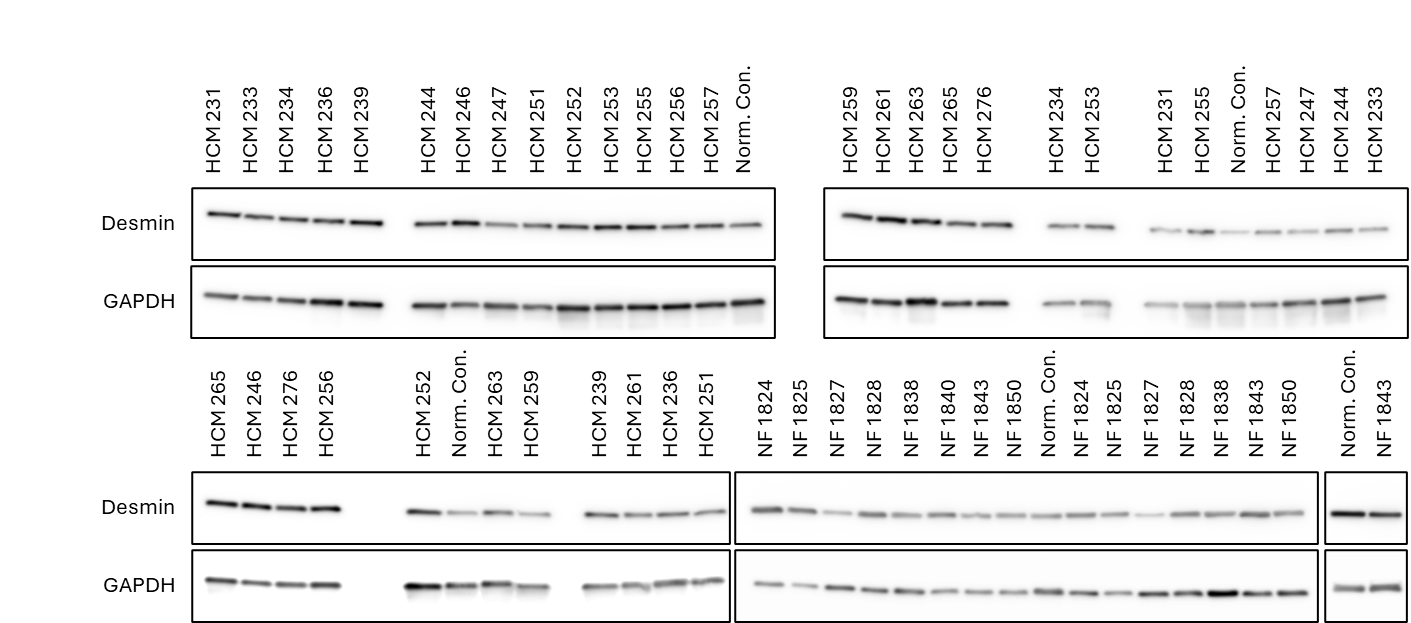

**Figure S18: Images of western blots of lamin A/C.**

**Figure S17: Images of western blots of desmin.**

Table S19: Uncorrected p-values of the Pearson R correlation of the microtubule code with nuclear invaginations in non-failing donors and HCM patients in Figure 5G.

|  | α-Tubulin | Detyr-Tubulin | Acetyl-Tubulin | Desmin | Lamin A/C | Invaginations |
| --- | --- | --- | --- | --- | --- | --- |
| α-Tubulin |  | 0.1142 | 0.0356 | 0.1381 | 0.1611 | 0.0613 |
| Detyr-Tubulin | 0.1142 |  | 0.0024 | 0.0543 | 0.0520 | 0.1245 |
| Acetyl-Tubulin | 0.0356 | 0.0024 |  | 0.0016 | 0.0190 | 0.0854 |
| Desmin | 0.1381 | 0.0543 | 0.0016 |  | 0.0156 | 0.1115 |
| Lamin A/C | 0.1611 | 0.0520 | 0.0190 | 0.0156 |  | 0.9149 |
| Invaginations | 0.0613 | 0.1245 | 0.0854 | 0.1115 | 0.9149 |  |

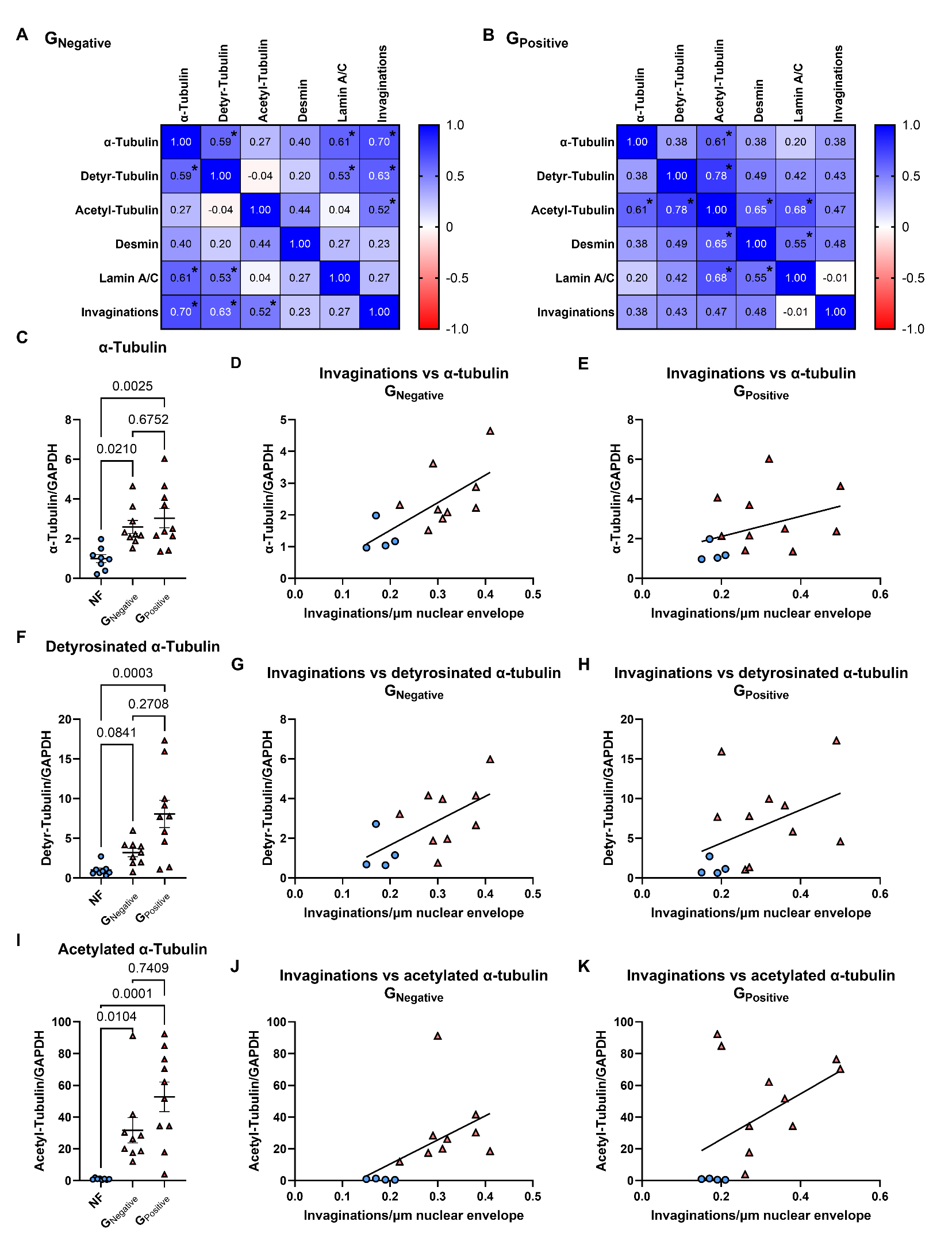

**Figure S19: Cytoskeletal protein expression in NF, G_Negative_ and G_Positive_ tissue and its correlation with nuclear invaginations.** Pearson R correlation of G_Negative_ (A) and G_Positive_ patients (B)(*= p<0.05). Quantification of protein levels of α-tubulin in NF donor, G_Negative_ and G_Positive_ patients (C). Simple linear regression of α-tubulin with nuclear invaginations in non-failing donors and G_Negative_ HCM patients (D, p=0.07, r^2^=0.49). Simple linear regression of α-tubulin with nuclear invaginations in non-failing donors and G_Positive_ HCM patients (E, p=0.18, r^2^=0.14). Quantification of protein levels of detyrosinated α-tubulin in NF donor, G_Negative_ and G_Positive_ patients (F). Simple linear regression of detyrosinated α-tubulin with nuclear invaginations in non-failing donors and G_Negative_ HCM patients (G, p=0.02, r^2^=0.40). Simple linear regression of detyrosinated α-tubulin with nuclear invaginations in non-failing donors and G_Positive_ HCM patients (H, p=0.12, r^2^=0.18). Quantification of protein levels of acetylated α-tubulin in NF donor, G_Negative_ and G_Positive_ patients (I). Simple linear regression of acetylated α-tubulin with nuclear invaginations in non-failing donors and G_Negative_ HCM patients (J, p=0.07, r^2^=0.27). Simple linear regression of acetylated α-tubulin with nuclear invaginations in non-failing donors and G_Positive_ HCM patients (K, p=0.09, r^2^=0.22). Protein levels were normalized to GAPDH. Every value is the average of two measurements per donor/patient (C, F, I). Data are expressed as mean ± standard error of the mean. Every colored symbol represents the average value per single donor or patient, N. (A, D, G, J) NF: N=4; G_Negative_: N=9. (B, E, H, K) NF: N=4; G_Positive_: N=10. (C, F, I) NF: N=8; G_Negative_: N=9; G_Positive_: N=10. Statistical tests: One-way ANOVA (C), Kruskal-Wallis test (F, I).

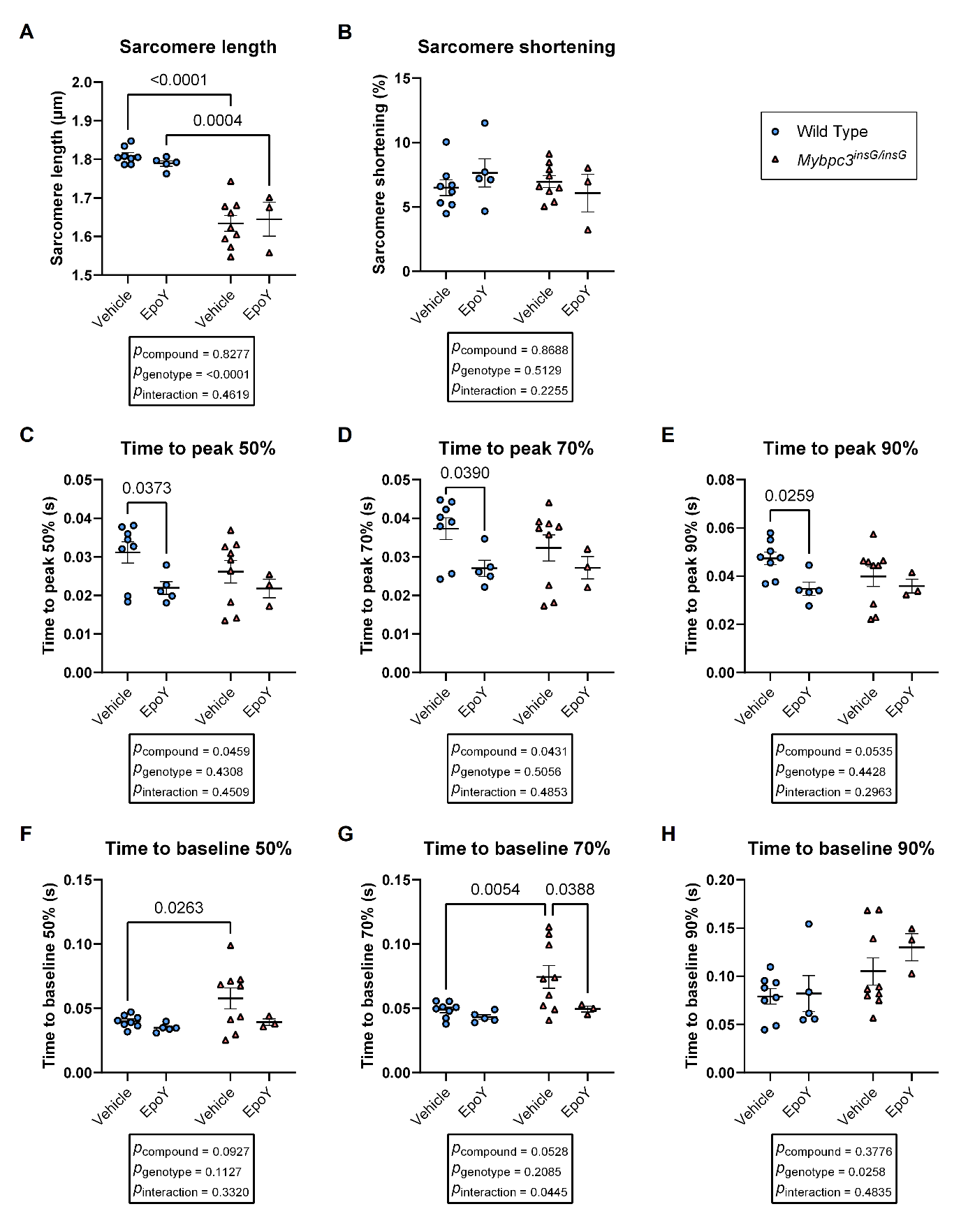

**Figure S20: Contraction and relaxation time of isolated cardiomyocytes of WT and Mybpc3^insG/insG^ mice upon addition of epoY.** Quantification of the resting sarcomere length (A) and maximum sarcomere shortening (B) in isolated cardiomyocytes of WT and Mybpc3^insG/insG^ mice upon addition of epoY in comparison with the vehicle. Quantification of the time to peak at a sarcomere shortening of 50% (A), 70% (B) and 90% (C) of maximum sarcomere shortening in isolated cardiomyocytes of WT and Mybpc3^insG/insG^ mice upon addition of epoY in comparison with the vehicle. Quantification of the relaxation using time to baseline at a sarcomere shortening of 50% (D), 70% (E) and 90% (F) of maximum sarcomere shortening in isolated cardiomyocytes of WT and Mybpc3^insG/insG^ mice upon addition of epoY in comparison with the vehicle. Every colored symbol represents the average value per single mouse, N. The data are expressed as mean ± standard error of the mean. Wild Type, vehicle: N=8 n=223; WT, epoY: N=5 n=119; Mybpc3^insG/insG^, vehicle: N=9 n=145; Mybpc3^insG/insG^, epoY: N=3 n=29. Statistical tests: two-way ANOVA (A-H).

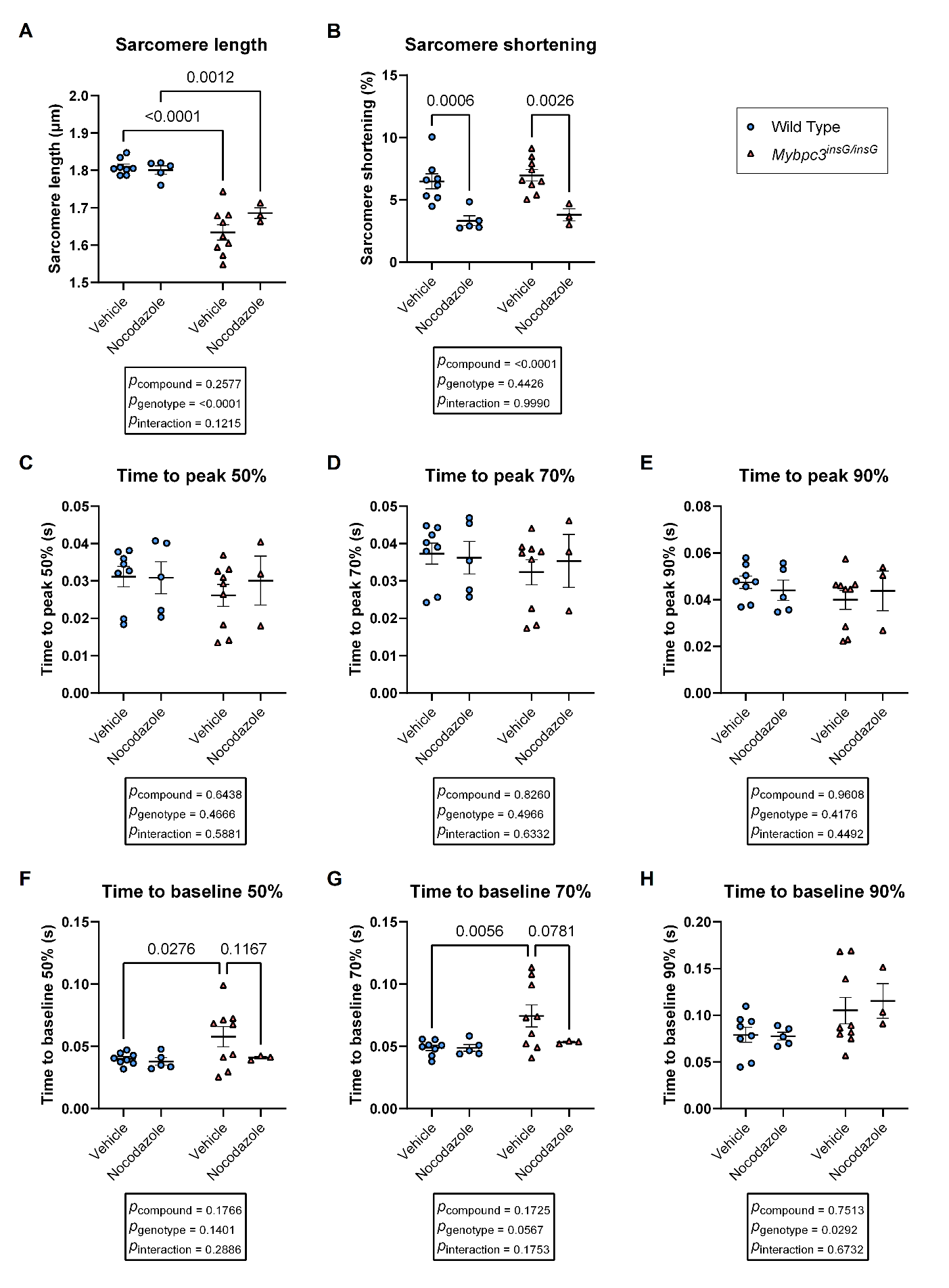

**Figure S21: Contraction and relaxation time of isolated cardiomyocytes of WT and Mybpc3^insG/insG^ mice upon addition of nocodazole.** Quantification of the resting sarcomere length (A) and maximum sarcomere shortening (B) in isolated cardiomyocytes of WT and Mybpc3^insG/insG^ mice upon addition of nocodazole in comparison with the vehicle. Quantification of the time to peak at a sarcomere shortening of 50% (A), 70% (B) and 90% (C) of maximum sarcomere shortening in isolated cardiomyocytes of WT and Mybpc3^insG/insG^ mice upon addition of nocodazole in comparison with the vehicle. Quantification of the relaxation using time to baseline at a sarcomere shortening of 50% (D), 70% (E) and 90% (F) of maximum sarcomere shortening in isolated cardiomyocytes of WT and Mybpc3^insG/insG^ mice upon addition of nocodazole in comparison with the vehicle. Every colored symbol represents the average value per single mouse, N. The data are expressed as mean ± standard error of the mean. Wild Type, vehicle: N=8 n=223; WT, nocodazole: N=5 n=125; Mybpc3^insG/insG^, vehicle: N=9 n=145; Mybpc3^insG/insG^, nocodazole: N=3 n=62. Statistical tests: two-way ANOVA (A-H).

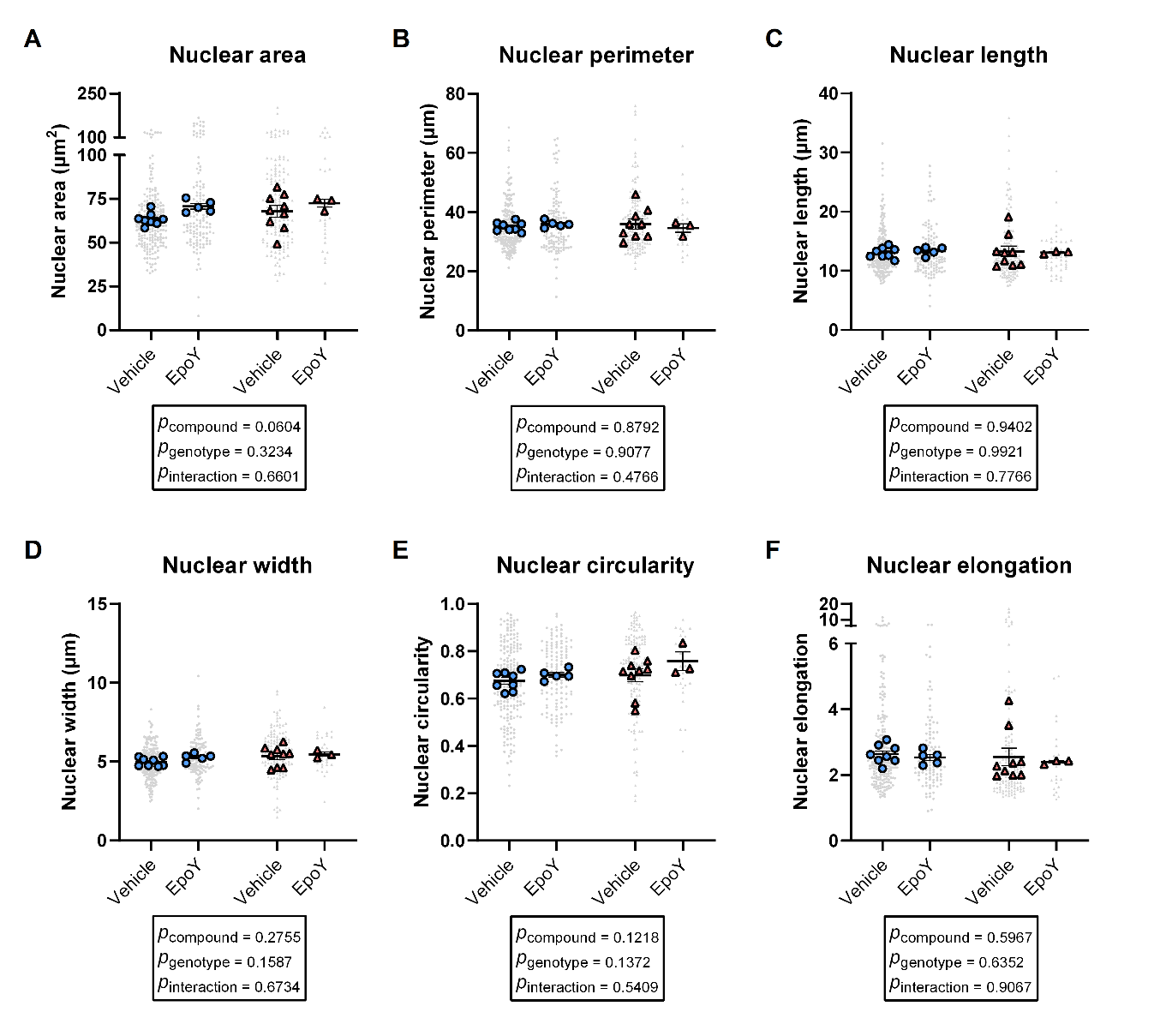

**Figure S22: Nuclear morphology at full relaxation upon addition of microtubule modifier epoY.** Quantification of nuclear area (A), nuclear perimeter (B), nuclear length (C), nuclear width (D), nuclear circularity (E) and nuclear elongation (F) upon the addition of vehicle or epoY in fully relaxed WT and Mybpc3^insG/insG^ cardiomyocytes. Data are expressed as mean ± standard error of the mean. Every grey symbol represents the value of a single nucleus, n, and every colored symbol represents the average value per single mouse, N. (A-F) WT, vehicle: N=8 n=223; WT, epoY: N=5 n=118; Mybpc3^insG/insG^, vehicle: N=9 n=145; Mybpc3^insG/insG^, epoY: N=3 n=29. Statistical tests: 2-way ANOVA (A-F).

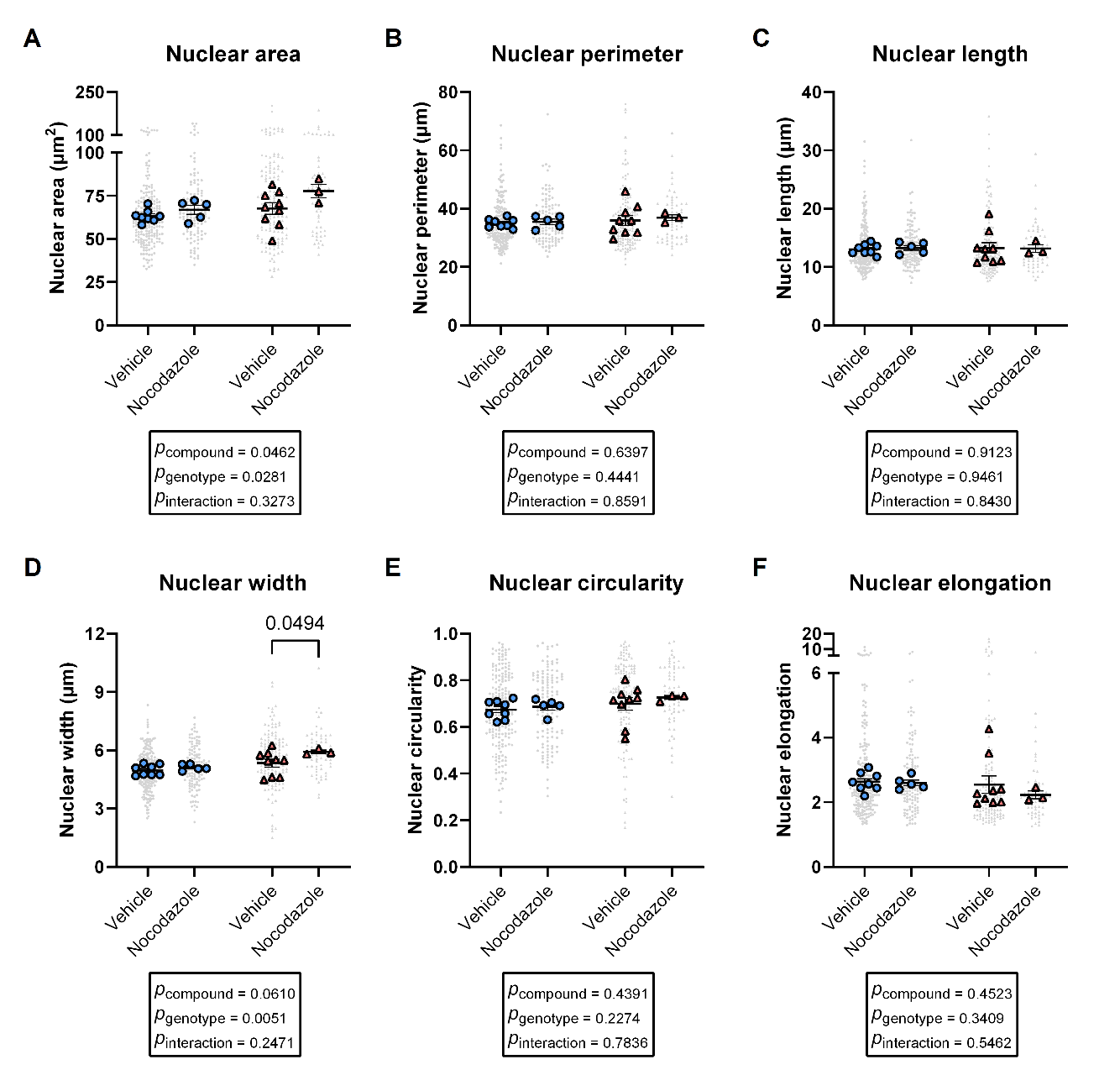

**Figure S23: Nuclear morphology at full relaxation upon addition of microtubule modifier nocodazole.** Quantification of nuclear area (A), nuclear perimeter (B), nuclear length (C), nuclear width (D), nuclear circularity (E) and nuclear elongation (F) upon the addition of vehicle or nocodazole in fully relaxed WT and Mybpc3^insG/insG^ cardiomyocytes. Data are expressed as mean ± standard error of the mean. Every grey symbol represents the value of a single nucleus, n, and every colored symbol represents the average value per single mouse, N. (A-F) WT, vehicle: N=8 n=223; WT, nocodazole: N=5 n=124; Mybpc3^insG/insG^, vehicle: N=9 n=145; Mybpc3^insG/insG^, nocodazole: N=3 n=62. Statistical tests: 2-way ANOVA (A-F).

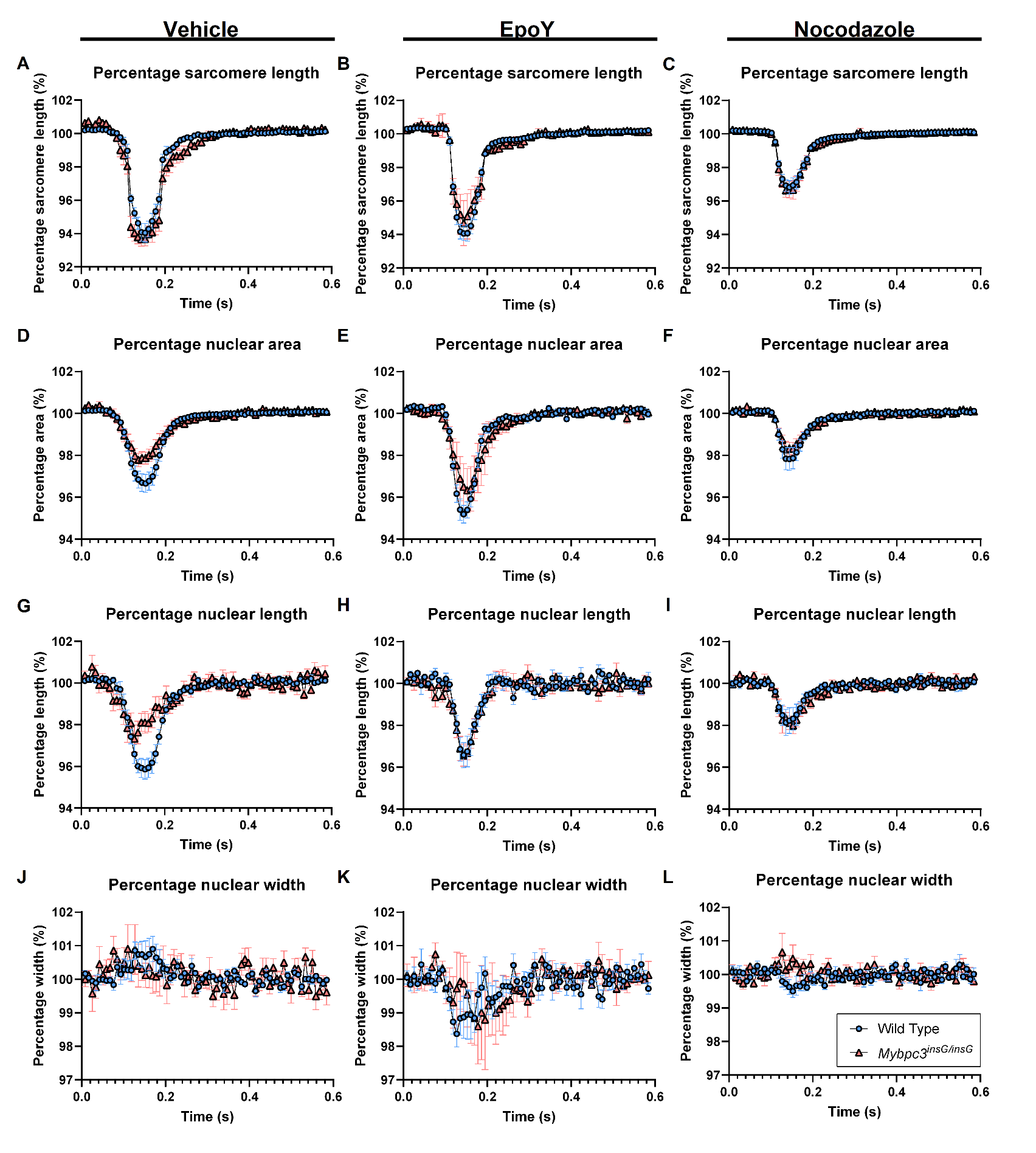

**Figure S24: Nuclear deformation percentage curves during contraction in WT and Mybpc3^insG/insG^ cardiomyocytes with vehicle or microtubule modifying compounds.** Quantification of the sarcomere length, nuclear area, nuclear length and nuclear width over time in WT and Mybpc3^insG/insG^ cardiomyocytes upon addition of vehicle (A, D, G, J), epoY (B, E, H, K) and nocodazole (C, F, I, L). Data are expressed as mean ± standard error of the mean. Every colored symbol represents the average value per single mouse, N. (A, D, G, J) WT, vehicle: N=8 n=223; (B, E, H, K) WT, epoY: N=5 n=118; (C, F, I, L) WT, nocodazole: N=5 n=124; (A, D, G, J) Mybpc3^insG/insG^, vehicle: N=9 n=145; (B, E, H, K) Mybpc3^insG/insG^, epoY: N=3 n=29; (C, F, I, L) Mybpc3^insG/insG^, nocodazole: N=3 n=62.

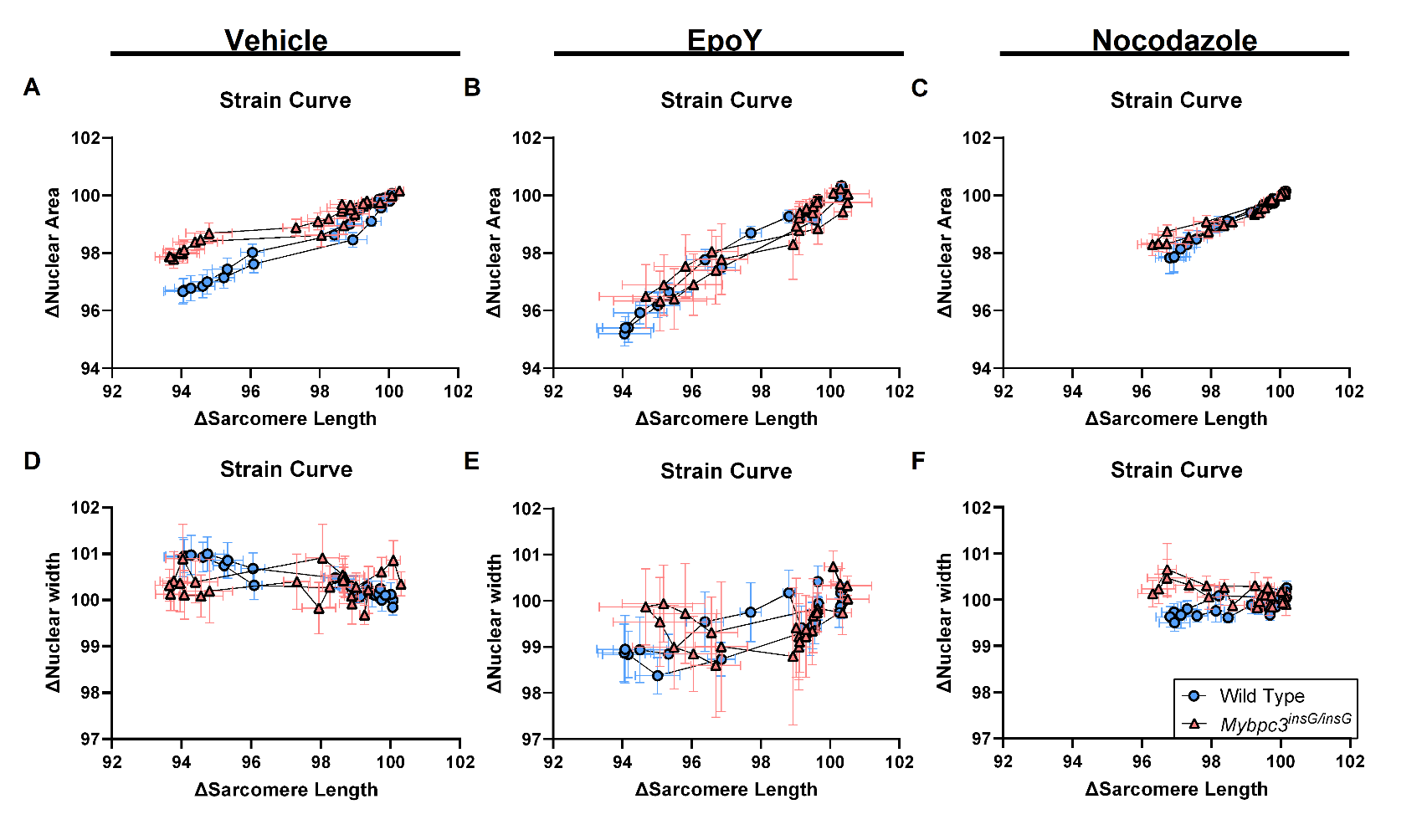

**Figure S25: Strain curves of nuclear area and nuclear width of WT versus Mybpc3^insG/insG^ cardiomyocytes.** Sarcomere-nuclear strain curves generated by calculating the change in sarcomere length versus the change in nuclear area upon the addition of vehicle (A), epoY (B) and nocodazole (C). Sarcomere-nuclear strain curves generated by calculating the change in sarcomere length versus the change in nuclear width upon the addition of vehicle (D), epoY (E) and nocodazole (F) Data are expressed as mean ± standard error of the mean. Every colored symbol represents the value of the average of the mice in their group, N. (A, D) WT, vehicle: N=8 n=223; (B, E) WT, epoY: N=5 n=118; (C, F) WT, nocodazole: N=5 n=124; (A, D) Mybpc3^insG/insG^, vehicle: N=9 n=145; (B, E) Mybpc3^insG/insG^, epoY: N=3 n=29; (C, F) Mybpc3^insG/insG^, nocodazole: N=3 n=62.

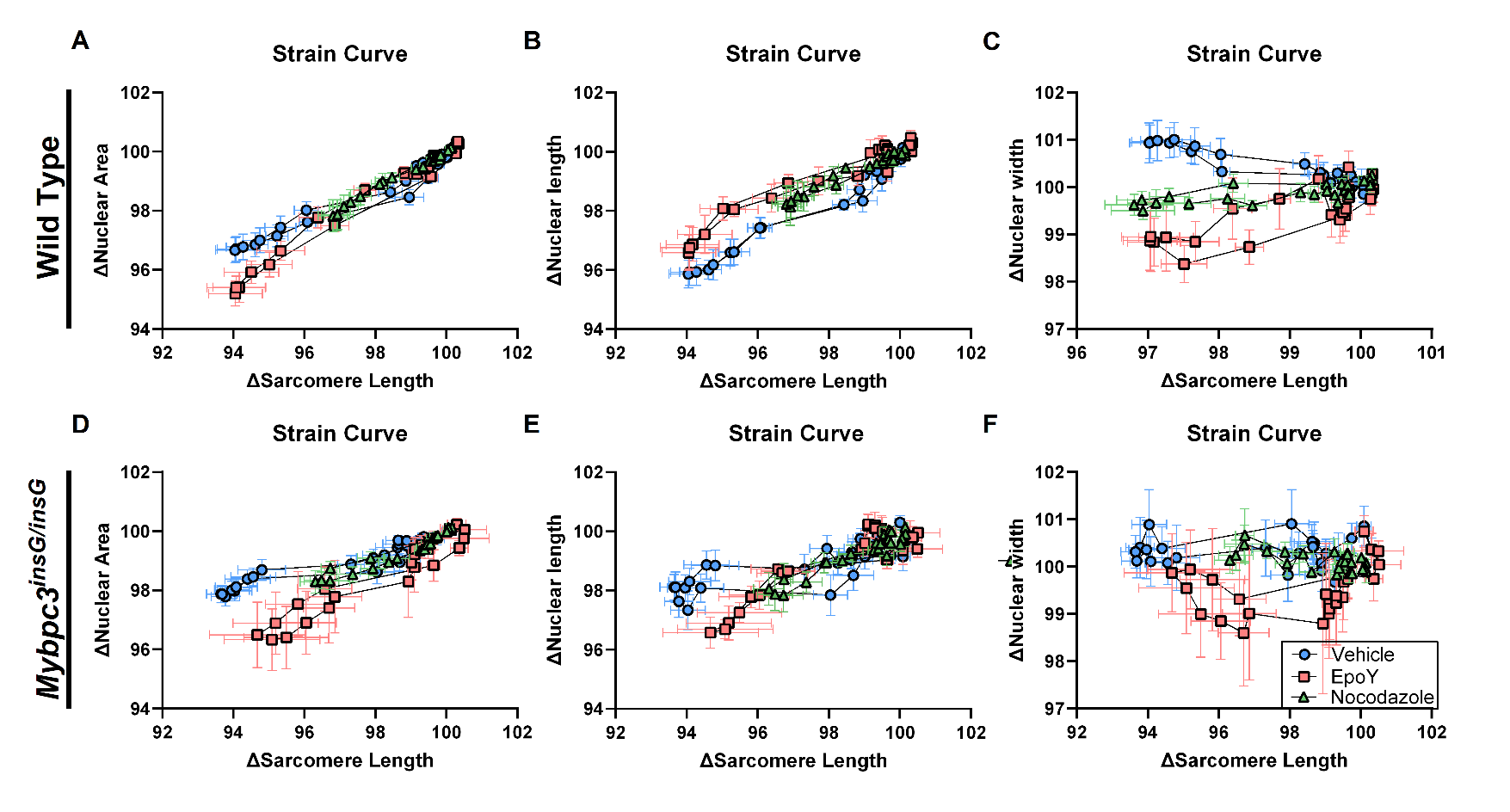

**Figure S26: Strain curves of nuclear area (A, D), nuclear length (B, E) and nuclear width (C, F) of Vehicle versus epoY versus Nocodazole in WT or Mybpc3^insG/insG^ cardiomyocytes.** Sarcomere-nuclear strain curves generated by calculating the change in sarcomere length versus the change in nuclear area upon the addition of vehicle, epoY and nocodazole in WT (A) and Mybpc3^insG/insG^ (D). Sarcomere-nuclear strain curves generated by calculating the change in sarcomere length versus the change in nuclear length upon the addition of vehicle, epoY and nocodazole in WT (B) and Mybpc3^insG/insG^ (E). Sarcomere-nuclear strain curves generated by calculating the change in sarcomere length versus the change in nuclear width upon the addition of vehicle, epoY and nocodazole in WT (C) and Mybpc3^insG/insG^ (F). Data are expressed as mean ± standard error of the mean. Every colored symbol represents the value of the average of the mice in their group, N. (A-C) WT, vehicle: N=8 n=223; (A-C) WT, epoY: N=5 n=118; (A-C)) WT, nocodazole: N=5 n=124; (D-F) Mybpc3^insG/insG^, vehicle: N=9 n=145; (D-F) Mybpc3^insG/insG^, epoY: N=3 n=29; (D-F) Mybpc3^insG/insG^, nocodazole: N=3 n=62.

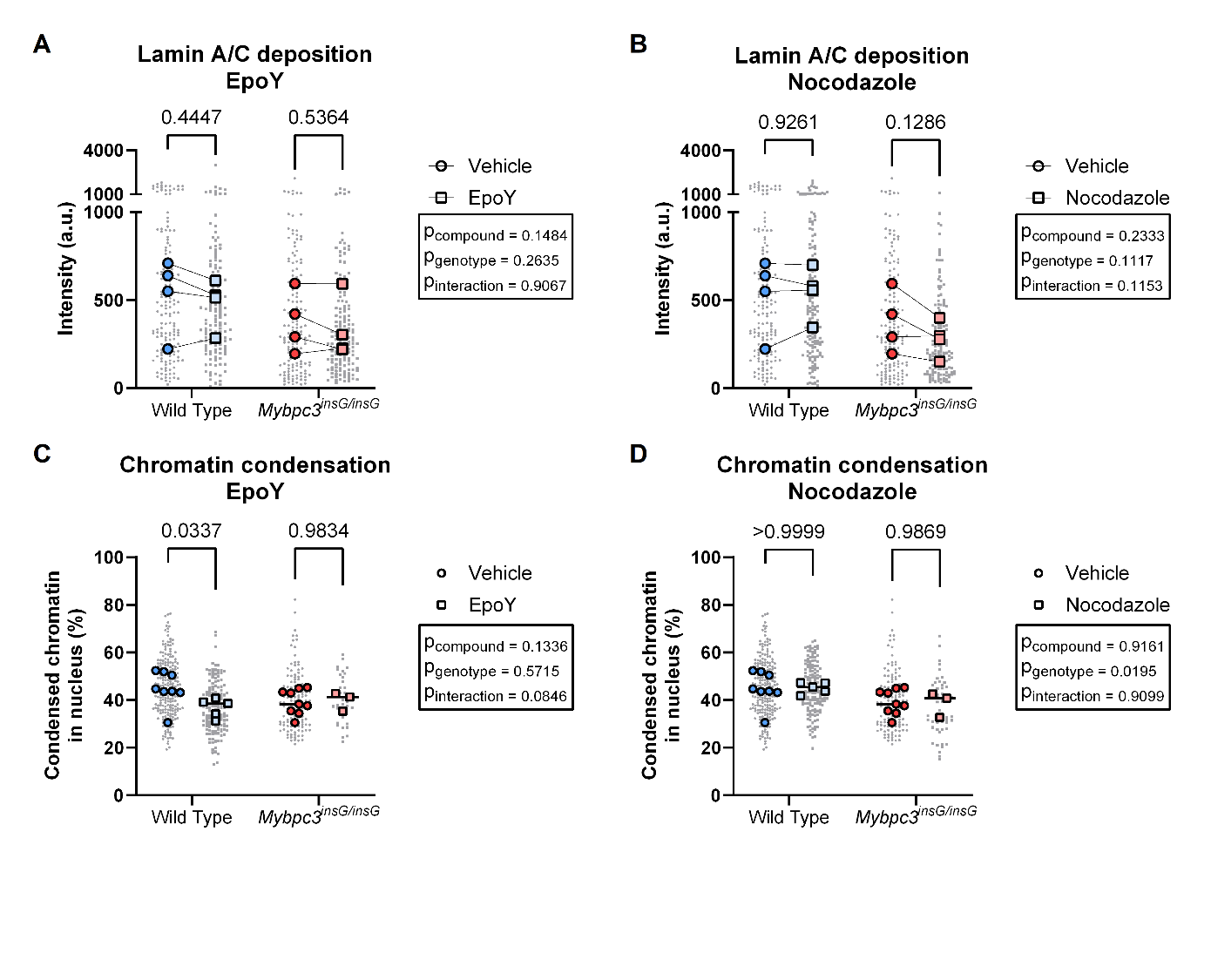

**Figure S27: Lamin A/C deposition and chromatin condensation upon the addition of microtubule modifiers nocodazole and epoY.** Quantification of Lamin A/C deposition (A, B) and chromatin condensation (C, D) in WT and Mybpc3^insG/insG^ cardiomyocytes upon the addition of epoY (A, C) or nocodazole (B, D). Lamin A/C deposition is determined as the mean intensity of the Lamin A/C signal (A). Chromatin condensation is determined as the percentage of heterochromatin in the nucleus (B). Data are expressed as mean ± standard error of the mean. Every grey symbol represents the value of a single nucleus, n, and every colored symbol represents the average value per single mouse, N. Colored symbols are linked when the vehicle and epoY or Nocodazole treated cardiomyocytes come from the same mouse. (A) WT, vehicle: N=4 n=156; WT, epoY: N=4 n=142; Mybpc3^insG/insG^, vehicle: N=4 n=154; Mybpc3^insG/insG^, epoY: N=4 n=150, (B) WT, vehicle: N=4 n=156; WT, nocodazole: N=4 n=161; Mybpc3^insG/insG^, vehicle: N=4 n=154; Mybpc3^insG/insG^, nocodazole: N=4 n=147, (C) WT, vehicle: N=8 n=218; WT, epoY: N=5 n=150; Mybpc3^insG/insG^, vehicle: N=9 n=137; Mybpc3^insG/insG^, epoY: N=3 n=34, (D) WT, vehicle: N=8 n=218; WT, nocodazole: N=5 n=149; Mybpc3^insG/insG^, vehicle: N=9 n=137; Mybpc3^insG/insG^, nocodazole: N=3 n=52. Statistical tests: two-way ANOVA (A-D).

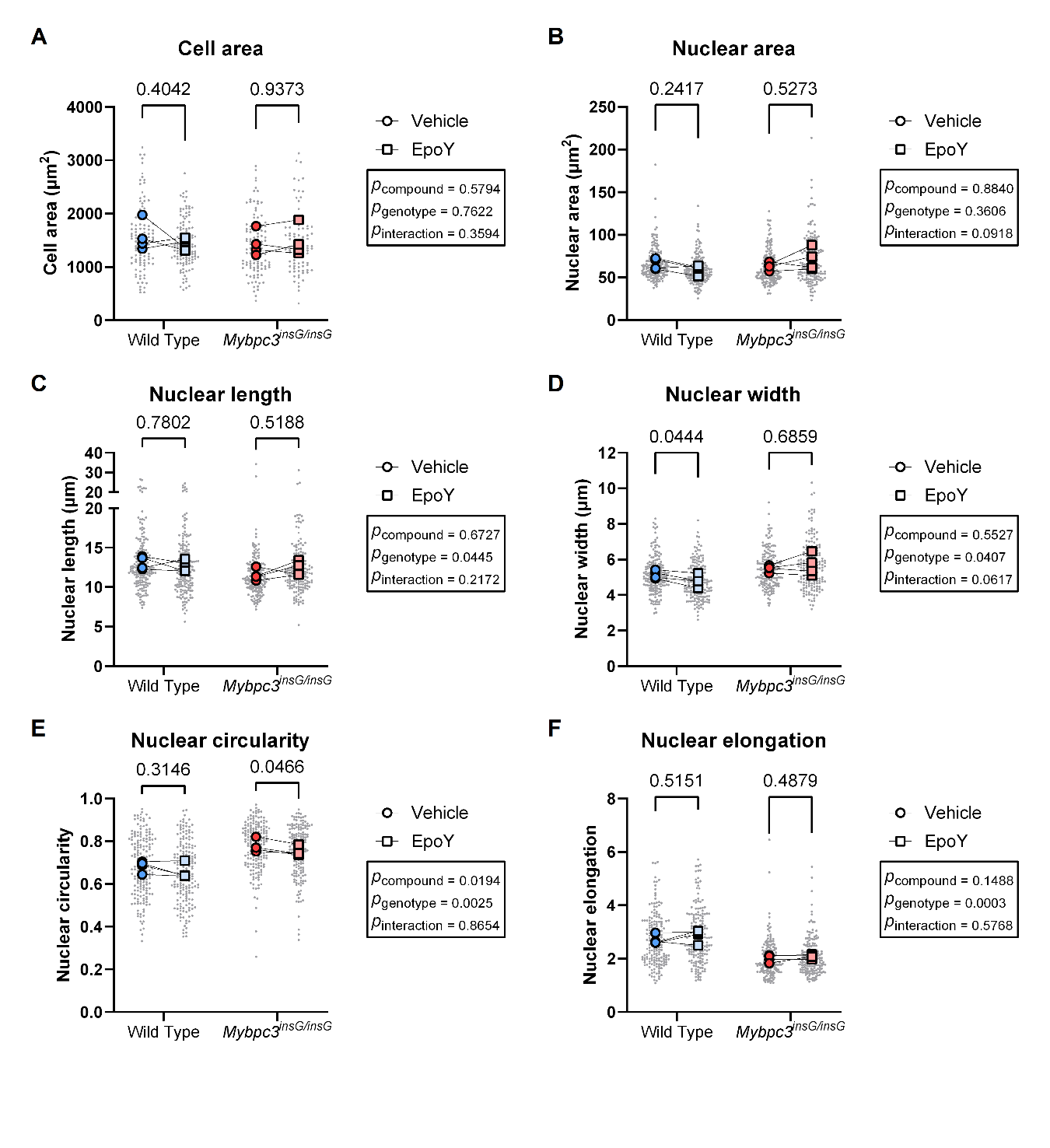

**Figure S28: Cell area and nuclear morphology upon the addition of microtubule modifier epoY.** Quantification of the cell area (A), nuclear area (B), nuclear length (C), nuclear width (D), nuclear circularity (E) and nuclear elongation (F) upon addition of vehicle or epoY. Data are expressed as mean ± standard error of the mean. Every grey symbol represents the value of a single nucleus, n, and every colored symbol represents the average value per single mouse, N. Colored symbols are linked when the vehicle and epoY treated cardiomyocytes come from the same mouse. (A) WT, vehicle: N=4 n=106; WT, epoY: N=4 n=114; Mybpc3^insG/insG^, vehicle: N=4 n=114; Mybpc3^insG/insG^, epoY: N=4 n=99, (B-F) WT, vehicle: N=4 n=174; WT, epoY: N=4 n=174; Mybpc3^insG/insG^, vehicle: N=4 n=172; Mybpc3^insG/insG^, epoY: N=4 n=164. Statistical tests: two-way ANOVA (A-F).

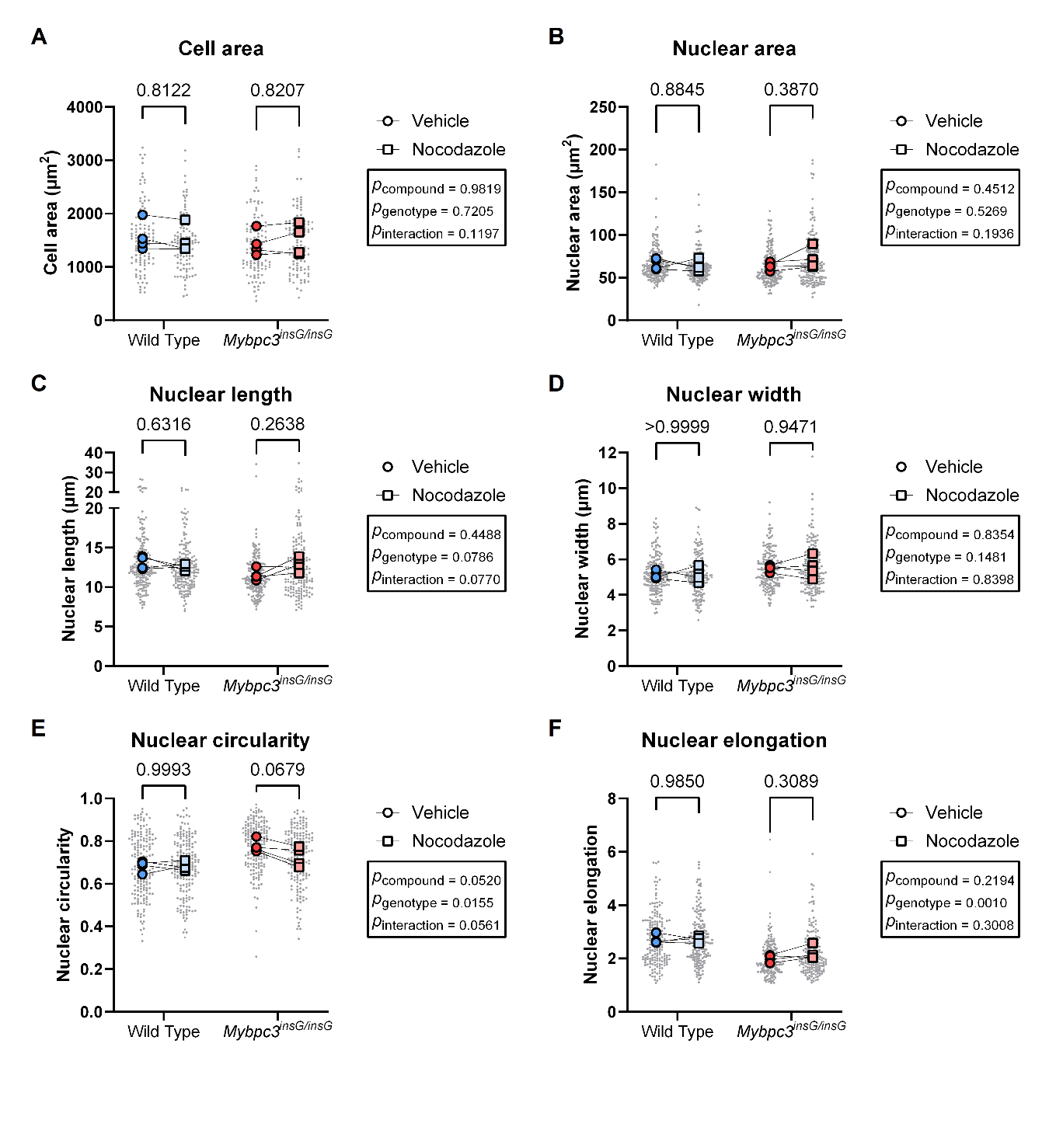

**Figure S29: Cell area and nuclear morphology upon the addition of microtubule modifier nocodazole.** Quantification of the cell area (A), nuclear area (B), nuclear length (C), nuclear width (D), nuclear circularity (E) and nuclear elongation (F) upon addition of vehicle or nocodazole. Data are expressed as mean ± standard error of the mean. Every grey symbol represents the value of a single nucleus, n, and every colored symbol represents the average value per single mouse, N. Colored symbols are linked when the vehicle and epoY treated cardiomyocytes come from the same mouse. (A) WT, vehicle: N=4 n=106; WT, nocodazole: N=4 n=114; Mybpc3^insG/insG^, vehicle: N=4 n=114; Mybpc3^insG/insG^, nocodazole: N=4 n=113, (B-F) WT, vehicle: N=4 n=174; WT, nocodazole: N=4 n=177; Mybpc3^insG/insG^, vehicle: N=4 n=172; Mybpc3^insG/insG^, nocodazole: N=4 n=173. Statistical tests: two-way ANOVA (A-F).
